# Cancer Driver Topologically Associated Domains identify oncogenic and tumor suppressive lncRNAs

**DOI:** 10.1101/2024.03.19.585685

**Authors:** Ziyan Rao, Min Zhang, Shaodong Huang, Chenyang Wu, Yuheng Zhou, Weijie Zhang, Xia Lin, Dongyu Zhao

**Affiliations:** Department of Biomedical Informatics, School of Basic Medical Sciences, Peking University, Beijing, 100191, China; State Key Laboratory of Vascular Homeostasis and Remodeling, Peking University, Beijing, 100191, China; Zhejiang Provincial Key Laboratory of Pancreatic Disease, the First Affiliated Hospital, Zhejiang University School of Medicine, Hangzhou, Zhejiang 310058, China; Zhejiang Provincial Key Laboratory of Cancer Molecular Cell Biology, Life Sciences Institute, Zhejiang University, Hangzhou, Zhejiang 310058, China; Department of Orthopaedic Surgery, Second Affiliated Hospital, School of Medicine, Zhejiang University, Hangzhou, Zhejiang 310058, China

## Abstract

Cancer long noncoding RNAs (lncRNAs) have been identified by experimental and in silico methods. However, current approaches for identifying cancer lncRNAs are not sufficient and effective. To uncover them, we focused on the core cancer driver lncRNAs, which directly interact with cancer driver protein-coding genes (PCGs). We investigated various aspects of cancer lncRNAs, including their expression patterns, genomic locations, and direct interactions with cancer driver PCGs, and developed a pipeline to unearth candidate cancer driver lncRNAs. Finally, we validated the reliability of potential cancer driver lncRNAs through functional analysis of bioinformatics data and CRISPR-Cas9 knockout experiments. We found that cancer lncRNAs were more concentrated in cancer driver topologically associated domains (CDTs), and CDT is an important feature in identifying cancer lncRNAs. Moreover, cancer lncRNAs showed a high tendency to co-express with and bind to cancer driver PCGs. Utilizing these distinctive characteristics, we developed a pipeline CADTAD to unearth candidate cancer driver lncRNAs in pan-cancer, including 256 oncogenic lncRNAs, 177 tumor suppressive lncRNAs, and 75 dual-function lncRNAs, as well as in three individual cancer types, and validated their cancer-related function. More importantly, the function of 10 putative cancer driver lncRNAs in prostate cancer was subsequently validated to influence cancer phenotype through cell studies. In light of these findings, our study offers a new perspective from the 3D genome to study the roles of lncRNAs in cancer. Furthermore, we provide a valuable set of potential lncRNAs that could deepen our understanding of the oncogenic mechanism of cancer driver lncRNAs.

## INTRODUCTION

Researchers often identify cancer driver protein-coding genes (PCGs) as cancer biomarkers (Martínez-Jiménez et al. 2020). However, coding genes represent less than 2% of the entire genome, while most of the genome is transcribed into noncoding RNAs, among which long noncoding RNAs (lncRNAs) form a major group. lncRNAs, typically longer than 200 base pairs with limited protein-coding potential (Derrien et al. 2012), can orchestrate the gene expression and chromatin architecture by engaging with DNA, RNA, or proteins (Melé and Rinn 2016; Statello et al. 2021; Gil and Ulitsky 2020; Herman et al. 2022; Mattick et al. 2023; Chang and Qi 2023). Numerous studies have reported the pivotal roles of the lncRNAs in cancer development (Huarte 2015; Schmitt and Chang 2016; Bhan et al. 2017).

The subcellular localization of lncRNAs largely dictates their function (Bridges et al. 2021). Most nuclear lncRNAs are strongly enriched within 3D proximity of their transcriptional loci (Quinodoz et al. 2021) and can exert a *cis*-regulatory influence on neighboring genes, exemplified by lncRNA *CDKN2B-AS1* mediates the epigenetic silencing of proximal genes *CDKN2A* and *CDKN2B* to increase cell proliferation (Yu et al. 2008; Kotake et al. 2011). Certain lncRNAs in the nucleus can also affect distant gene expression, such as *HOTAIR*, repressing the expression of a distant gene *HOXD* cluster through epigenetic silencing, resulting in enhanced tumor metastasis (Rinn et al. 2007; Gupta et al. 2010). Conversely, cytoplasmic lncRNAs, like *PTENP1*, can sequester microRNAs to stabilize *PTEN* (Poliseno et al. 2010). Hence, dissecting lncRNA functions offers key insights into cancer biology and therapeutic opportunities.

Traditional wet experimental methodologies for cancer lncRNA discovery are inefficient and costly, while most bioinformatic approaches have identified cancer-related lncRNAs with limited influence on tumorigenesis. Manual curation can collect cancer-related lncRNA datasets based on the existing literature (Vancura et al. 2021; Gao et al. 2021). However, this method exhibited some leniency in the inclusion criteria, such as including differentially expressed lncRNAs. The only evidence of expression difference is not sufficient to prove the pivotal roles of lncRNAs in tumorigenesis. Current machine learning models (Zhao et al. 2015; Zhang et al. 2018b, 2019a; Liu et al. 2020; Zhang et al. 2020; Yuan et al. 2021) only identified cancer-related lncRNAs reliant on the aforementioned collected datasets and failed to distinguish between oncogenic and tumor suppressive lncRNAs in silico. Therefore, current studies are not capable of effectively discerning cancer lncRNAs, which could play driver roles in cancer.

In eukaryotes, chromatin is organized into higher-order structures, one of which is the topologically associated domain (TAD). It represents a discrete region of gene regulation. Within TADs, DNA elements interact frequently, whereas inter-TAD chromatin interactions tend to be relatively less pronounced. Disruption of TAD boundaries can cause profound consequences, including long-range transcriptional dysregulation and aberrant gene expression (Nora et al. 2012; Valton and Dekker 2016).

Given the critical roles of lncRNAs’ strong correlations with the expression of proximate genes, potentially acting as *cis*-regulatory elements (Werner et al. 2017) and well-studied cancer driver PCGs, we hypothesized that linking lncRNAs to cancer driver PCGs would enable the effective identification of cancer driver lncRNAs, which are defined as those directly modulating the expression of cancer driver PCGs. They can be divided into oncogenic lncRNAs and tumor suppressive lncRNAs according to their association with cancer driver PCGs.

In this study, we first explored the relationship between cancer lncRNAs and cancer driver PCGs. Notably, cancer lncRNAs exhibited a pronounced propensity to localize within cancer driver topologically associated domains (CDTs). They also show higher levels of co-expression and lncRNA-DNA interactions with cancer driver PCGs. Based on these findings, we developed CADTAD, a computational pipeline to identify potential cancer driver lncRNAs across various cancer types. CADTAD identified 256 oncogenic lncRNAs, 177 tumor suppressive lncRNAs, and 75 dual-function lncRNAs for pan-cancer and putative cancer driver lncRNAs in three individual cancer types (prostate cancer, gastric cancer, and lung cancer). We validated the candidate driver lncRNAs utilizing cancer genomic, epigenomic, and phenotype data. Additionally, we conducted CRISPR-Cas9 knockout experiments to confirm the function of six oncogenic lncRNAs and four tumor suppressive lncRNAs specifically in prostate cancer.

## RESULTS

### Cancer lncRNAs tend to co-express with and close to cancer driver PCGs

We collected cancer driver PCGs from Cancer Gene Census (CGC) database v.94 (Sondka et al. 2018), and cancer lncRNAs from Cancer LncRNA Census 2 (CLC2) (Vancura et al. 2021) and Lnc2Cancer3.0 (Gao et al. 2021). Next, we explored the relationship between these cancer lncRNAs and cancer driver PCGs within the context of cancer (see Methods).

Our initial investigation focused on the associations between the expression of cancer lncRNAs and cancer driver PCGs. Cancer lncRNAs show both positive and negative correlations with cancer driver PCGs. To evaluate the levels of co-expression rather than the direction of the relationship, we used absolute values of expression correlation (Zhang and Horvath 2005; Langfelder and Horvath 2008). Here, co-expression refers to the phenomenon where two or more genes exhibit changes in expression levels under the same or similar conditions, and these changes can be positively or negatively correlated. We found that cancer lncRNAs obtained from both CLC2 and Lnc2Cancer exhibited significantly higher levels of co-expression with cancer driver PCGs compared with randomly selected lncRNAs or PCGs under identical expression levels (Fig. 1A, Supplemental Fig. S1A-C, see Methods). When considering the transcription start sites (TSSs) of cancer driver PCGs or randomly selected PCGs as the center, we observed a notable predominance of cancer lncRNAs in proximity to cancer driver PCGs compared with their distribution around randomly selected PCGs across a series of extended genomic regions (Fig. 1B, Supplemental Fig. S1D, see Methods). We calculated the distance to TSS when the cancer lncRNAs reached a certain quantile (Supplemental Table S1). We found that at the same quantile, cancer lncRNAs were located closer to the cancer driver PCGs than to randomly selected PCGs, whether in CLC2 or Lnc2Cancer. We also calculated the distance between cancer lncRNAs and the closest cancer driver/random PCGs and found that cancer lncRNAs were located nearer cancer driver PCGs than randomly selected PCGs (Supplemental Table S2, KS-test: CLC2 3.27×10^-18^, Lnc2Cancer 6.53 × 10^-36^). However, compared with randomly selected lncRNAs, the predominance of cancer lncRNAs in proximity to cancer driver PCGs is not obvious (Fig. 1B, Supplemental Table S1-2, KS-test: CLC2 6.94×10^-3^, Lnc2Cancer 4.11×10^-2^). At a 1 Mb distance, the number of cancer lncRNAs was higher than the number of randomly selected lncRNAs. To explore in detail the distribution difference between cancer lncRNAs and randomly selected lncRNAs around cancer driver PCGs, we calculated the quantity variation between them across a series of extended genomic regions. We found that cancer lncRNAs have a significantly higher number in close range (Supplemental Fig. S1E-F, H-I) at around 6Mb to cancer driver PCG TSS. We also obtained the closest lncRNAs around cancer driver PCGs and found that the selected lncRNAs were significantly enriched in the cancer lncRNA datasets (Supplemental Fig. S1G, J). However, if we replaced cancer driver PCGs with randomly selected PCGs or replaced cancer lncRNAs with randomly selected lncRNAs, the enrichments were not very significant (Supplemental Fig. S1G, J). Our findings indicate that cancer lncRNAs tend to be located near cancer driver PCGs. Their co-expression patterns suggest a potential interaction between their transcription, including that cancer lncRNAs possibly influence the expression of cancer driver PCGs.

**Fig. 1.**
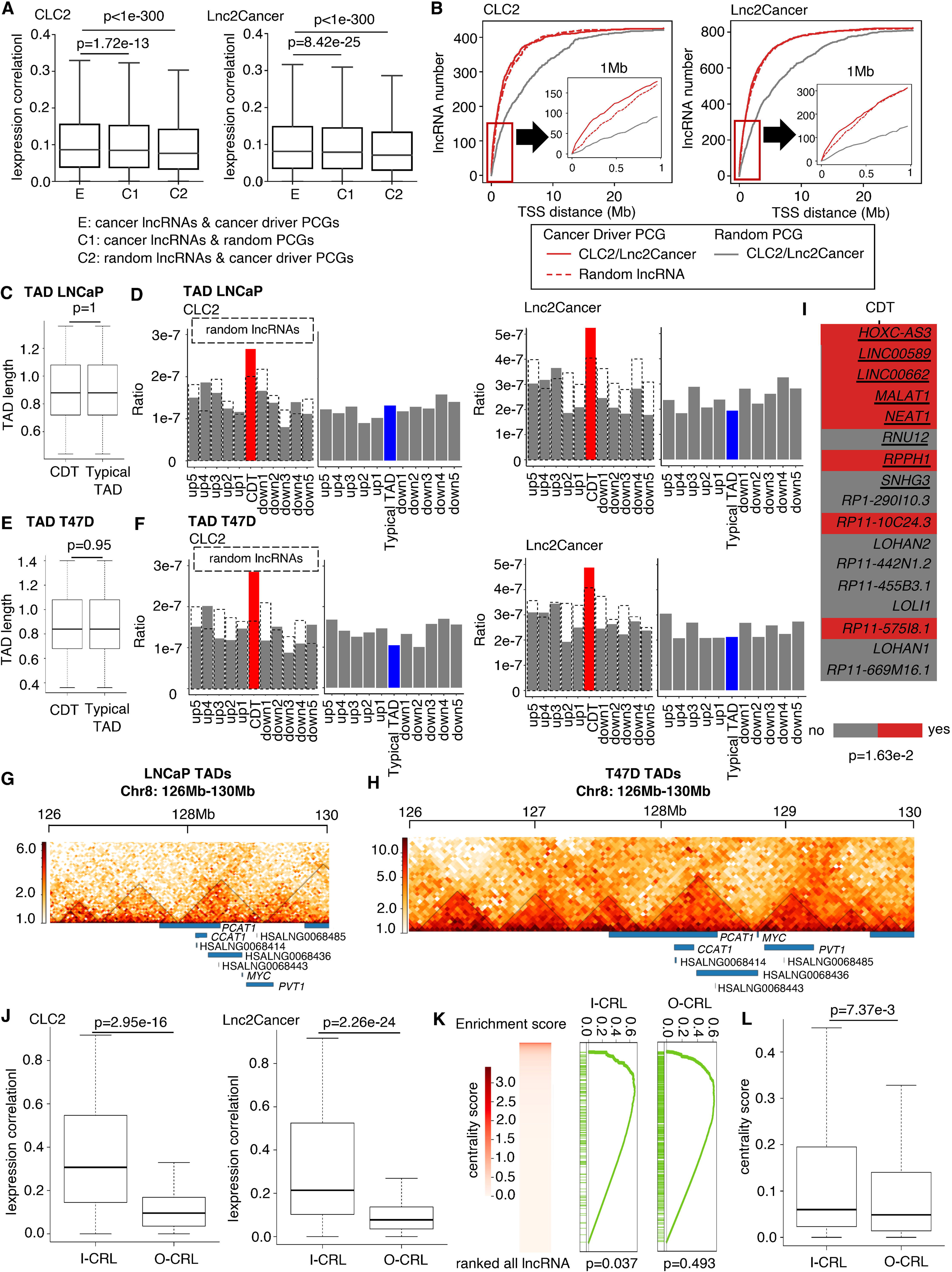
Cancer lncRNAs are more co-expressed with and located near cancer driver PCGs, especially in the same TAD. **A.** Boxplot to show the difference in absolute Spearman’s correlation between the experimental pair and two control pairs. P values are determined by one tail Wilcoxon test. **B.** Line plot of the statistics of the lncRNA number in CLC2/Lnc2Cancer in regions whose step size is 40kb extending from 40kb to 28Mb near the TSS of cancer driver/random PCGs. The red line indicates cancer driver PCGs, while the gray line indicates random PCGs. The solid line represents cancer lncRNA and the dashed line represents random lncRNA. **C.** Boxplot to indicate the TAD length of CDT and typical TAD are almost the same in LNCaP. P value is determined by two tail Wilcoxon test. **D.** Bar plot to show the density of cancer/random lncRNAs in each TAD of LNCaP. The dashed line indicates random lncRNAs. The red bar represents CDT, while the blue bar represents typical TAD. The grey bar represents flanking typical TAD. CDT: cancer driver TAD. **E.** Boxplot to indicate the TAD length of CDT and typical TAD are almost the same in T47D. P value is determined by two tail Wilcoxon test. **F.** Bar plot to show the density of cancer/random lncRNAs in each TAD of T47D. The dashed line indicates random lncRNAs. The red bar represents CDT, while the blue bar represents typical TAD. The grey bar represents flanking typical TAD. CDT: cancer driver TAD. **G.** The two-dimensional heatmap shows the normalized Hi-C interaction frequencies in LNCaP. Black line box out the calculated TAD. Blue rectangles represent genes. **H.** The two-dimensional heatmap shows the normalized Hi-C interaction frequencies in T47D. Black line box out the calculated TAD. Blue rectangles represent genes. **I.** Heatmap to show the CDT characteristic of the driver lncRNAs identified by ExInAtor2. The red box represents that lncRNA has this characteristic. The gray box represents that lncRNA doesn’t have this characteristic. Underlined lncRNAs represent known driver lncRNAs. P value is determined by one tail Fisher’s exact test. **J.** Boxplot to show the difference in absolute Spearman’s correlation between I-CRL pairs and O-CRL pairs. I-CRL: cancer lncRNAs in CDT. O-CRL: cancer lncRNAs outside CDT. P values are determined by one tail Wilcoxon test. **K.** GSEA analysis shows enrichment of the I-CRLs and O-CRLs in hub lncRNAs in a cancer-related lncRNA-mRNA network. **L.** Boxplot to indicate the difference in centrality score between I-CRLs and O-CRLs. P value is determined by one tail Wilcoxon test.

### CDT is an important feature in identifying cancer lncRNAs

As TADs are characterized by high-density chromatin contacts, and their alterations may cause aberrant gene expression (Valton and Dekker 2016), we considered TADs in cancer as functional units where lncRNAs may affect the expression of cancer driver PCGs. TADs can be categorized into two groups: CDTs, which are defined as TADs that contain at least one cancer driver PCG, and the remaining TADs, which are referred to as typical TADs. In the prostate cancer cell line LNCaP, the densities of cancer lncRNAs were higher in CDTs than in those of neighboring typical TADs or randomly selected typical TADs of similar length, whereas the densities of randomly selected lncRNAs did not show this tendency (Fig. 1C-D, Supplemental Fig. S2A-B, Supplemental Table S3, see Methods). We observed consistent results in the breast cancer cell line T47D (Fig. 1E-F, Supplemental Table S3). The known oncogenic lncRNA *PVT1* is adjacent to the oncogene *MYC* in the same TAD in the T47D cell line. A previous study demonstrated that lncRNA *PVT1* enhanced the stability of *MYC*, resulting in high protein levels and the promotion of cancer progression (Tseng et al. 2014). Moreover, lncRNA *CCAT1* could regulate *MYC* transcriptional expression by promoting loop interaction, and could also be found in the same TAD in the LNCaP cell line. However, lncRNA *PCAT1*, which overlapped with the TAD boundary in the LNCaP and T47D cell lines, was co-amplified with *MYC* and induced *MYC* expression through a post-transcriptional mechanism (Huarte 2015; Schmitt and Chang 2016) (Fig. 1G-H). These examples indicate a potent *cis*-regulatory mechanism of cancer lncRNAs regulating cancer driver PCGs in the CDT. In addition, we observed that almost half of the cancer driver lncRNAs identified in a recent publication (Esposito et al. 2023) were located in CDTs (Fig. 1I).

To explore the effect of CDTs, we then divided cancer lncRNAs into I-CRLs (inside CDT) and O-CRLs (outside CDT) for each CDT. We also defined that I-CRL pairs represent the pairs of I-CRL and cancer driver PCG, while O-CRL pairs represent the pairs of O-CRL and cancer driver PCG. We performed exact matching to compare the co-expression levels between I-CRL and O-CRL pairs (Supplemental Fig. S3A, see Methods), and observed that I-CRLs exhibited a higher degree of co-expression with cancer driver PCGs within the same TAD compared with O-CRLs (Fig. 1J). The distance between lncRNA-mRNA neighboring pairs could affect co-expression levels (Derrien et al. 2012). To control the effect of distance, we obtained I-CRL pairs and the closest O-CRL pairs for the same cancer driver PCGs (see Methods). However, the distances of I-CRL pairs were still closer than those of the O-CRL pairs (Supplemental Fig. S3B, D), and the co-expression levels of I-CRL pairs were also significantly higher than those of O-CRL pairs (Supplemental Fig. S3C, E). The location distribution of the cancer driver PCGs within the CDTs occurred in two cases (Supplemental Fig. S4A): 1) The PCG is distributed near the boundary of the CDT. 2) The cancer driver PCG is in the middle of the CDT. We found that cancer driver PCGs were mainly distributed in the middle of the CDTs (Supplemental Fig. S4B, see Methods). This phenomenon can explain why, even though we tried our best to choose the closest O-CRL pairs, the distances of O-CRL pairs were still much greater than those of I-CRL pairs (Supplemental Fig. S3B, D).

We then selected O-CRL pairs that have a minimum discrepancy of distance with I-CRL pairs to make their distance similar (Supplemental Table S4, see Methods). In this way, a lot of I-CRL pairs were excluded since some I-CRL pairs couldn’t have O-CRL pairs with similar distances, but we kept the distance similar and found the expression correlation levels of I-CRL pairs were significantly higher than those of O-CRL pairs (Supplemental Table S4).

Additionally, we matched lncRNAs inside and outside CDTs with similar distances to cancer driver PCGs (Supplemental Fig. S5A, see Methods), and found that lncRNAs in the CDTs had a slightly significant enrichment in cancer lncRNAs compared with the lncRNAs outside CDTs (Supplemental Fig. S5B-C). Considering all the lncRNAs located in the CDT, we found the enrichment degree to be higher (Supplemental Fig. S5D). These results indicated that after removing some lncRNAs that are very close to the cancer driver PCGs in CDTs, lncRNAs in the CDTs still remain slightly enriched in cancer lncRNAs.

Using a generalized linear regression model (GLM) to control the distance, we found that the feature CDT significantly affected co-expression levels between cancer lncRNAs and cancer driver PCGs (Supplemental Table S5-6, see Methods). The plus signs of CDT coefficients and minus signs of distance coefficients suit our expectations: cancer lncRNAs located in the CDTs positively contribute to co-expression levels with cancer driver PCGs, while the degree of expression correlation values decreases with distance. This means that when we control the distance between cancer driver PCGs and cancer lncRNAs, cancer lncRNAs located in CDTs have higher co-expression levels with cancer driver PCGs than those not located in CDTs. Furthermore, when the co-expression levels were shuffled, CDT and distance features did not significantly contribute to the degree of expression correlations. When the CDT and distance features were separately shuffled, only the corresponding feature did not significantly contribute to the co-expression levels. Moreover, the shuffling process decreased the R-squared value (Supplemental Table S5-6). Therefore, the co-expression levels are not only determined by distance, but CDT can also positively contribute to the response values.

To further validate CDT specificity, we calculated co-expression levels between cancer lncRNAs and non-cancer PCGs in typical TADs or CDTs. The typical TAD (TT) was defined in the previous analysis as the control group. We also named the non-cancer PCGs in CDT as typical genes (TGs) in CDT. Cancer lncRNAs and cancer driver PCGs in CDTs consistently showed higher co-expression levels than those of cancer lncRNAs and non-cancer PCGs in either typical TADs or CDTs (Supplemental Fig. S6A-B), reinforcing the specificity of the CDT effect.

lncRNAs can play an important role in biological networks (Zhang et al. 2018a). Here, we constructed a cancer-related lncRNA-mRNA network (Supplemental Fig. S7A-B, see Methods). Although both I-CRLs and O-CRLs exhibited a propensity to serve as hub-lncRNAs (Fig. 1K), the centrality score comparison and GSEA analysis revealed that the I-CRLs showed a significantly higher likelihood than the O-CRLs to function as hub-lncRNAs (Fig. 1K-L), highlighting the more important role of I-CRLs in cancer-related processes.

We also examined the effectiveness of cancer lncRNAs identified using CDT and expression correlation and found that a co-expression level threshold of 0.5 has a similar power to discover cancer lncRNAs with CDT (Supplemental Fig. S8A-B).

These results suggest that CDT is an important feature in discovering cancer lncRNAs. Moreover, cancer lncRNAs in the CDTs show higher location density near and higher co-expression levels with cancer driver PCGs, which may have the potential to regulate the expression of cancer driver PCGs.

### Cancer driver lncRNAs directly interact with cancer driver PCGs

LncRNAs can play pivotal roles in the regulation of gene expression by interacting with proteins and nucleic acids (Herman et al. 2022; Gil and Ulitsky 2020). Some studies have indicated that lncRNAs have the ability to interact with DNA, RNA, or protein to modulate many important cancer-related processes (Schmitt and Chang 2016). A pan-cancer analysis of lncRNA regulatory interactions suggested the dysregulation of hundreds of lncRNA targets and altered the expression of cancer genes and pathways in the tumor context (Chiu et al. 2018). Considering that gene interactions are more frequent within a TAD and that lncRNAs have the potential to modify gene expression, we are motivated to identify the core driver lncRNAs in CDTs, which are defined as lncRNAs that directly influence the expression of cancer driver PCGs. Chiu et al. constructed lncRNA regulatory networks in pan-cancer, and the networks included four types of lncRNA regulatory interactions (decoy, co-factor, guide, and switch) involving more than two types of molecules. To simplify the regulation mechanism, we considered only the most direct interaction between lncRNAs and DNA involving only two types of molecules to influence gene expression. Thus far, lncRNAs have been found to directly bind DNAs in two ways: triplex and R-loops, which may have different effects on gene expression (Niehrs and Luke 2020; Li et al. 2016a; Statello et al. 2021). Triplex is a triple-helical structure in which double-stranded DNA accommodates a single-stranded RNA in the major groove (Buske et al. 2012). Many studies have shown that lncRNAs can regulate target gene expression by forming a triplex (Leisegang et al. 2024; Mondal et al. 2015; Choudhury et al. 2021). We used Triplexator to predict the triplex-forming sites between lncRNA *MEG3* and cancer driver PCGs, and found that the predicted sites could match the *MEG3* binding sites (Fig. 2A, see Methods) (Buske et al. 2012; Mondal et al. 2015). The R-loop is a three-stranded nucleic acid structure that forms an RNA-DNA hybrid between a nascent guanine-rich RNA transcript segment and a DNA template while leaving the non-template DNA strand in a single-stranded conformation (Jenjaroenpun et al. 2015). lncRNAs can also regulate target genes through the R-loop (Arab et al. 2019; Ariel et al. 2020). Here, the R-loops between lncRNA *HOTTIP* and cancer driver PCGs were predicted using QmRLFS-finder (Jenjaroenpun et al. 2015), and we found that the predicted sites could match the R-loop formation sites (Fig. 2B). We then used the Triplexator and QmRLFS-finder to predict triplex and R-loops separately between the lncRNAs and the corresponding PCGs. Compared with randomly selected PCGs or lncRNAs, cancer lncRNAs have more potential triplex binding sites and R-loops with cancer driver PCGs (Fig. 2C-D, see Methods). The results suggested that cancer lncRNAs have a higher affinity with cancer driver PCGs through RNA-DNA interaction.

**Fig. 2.**
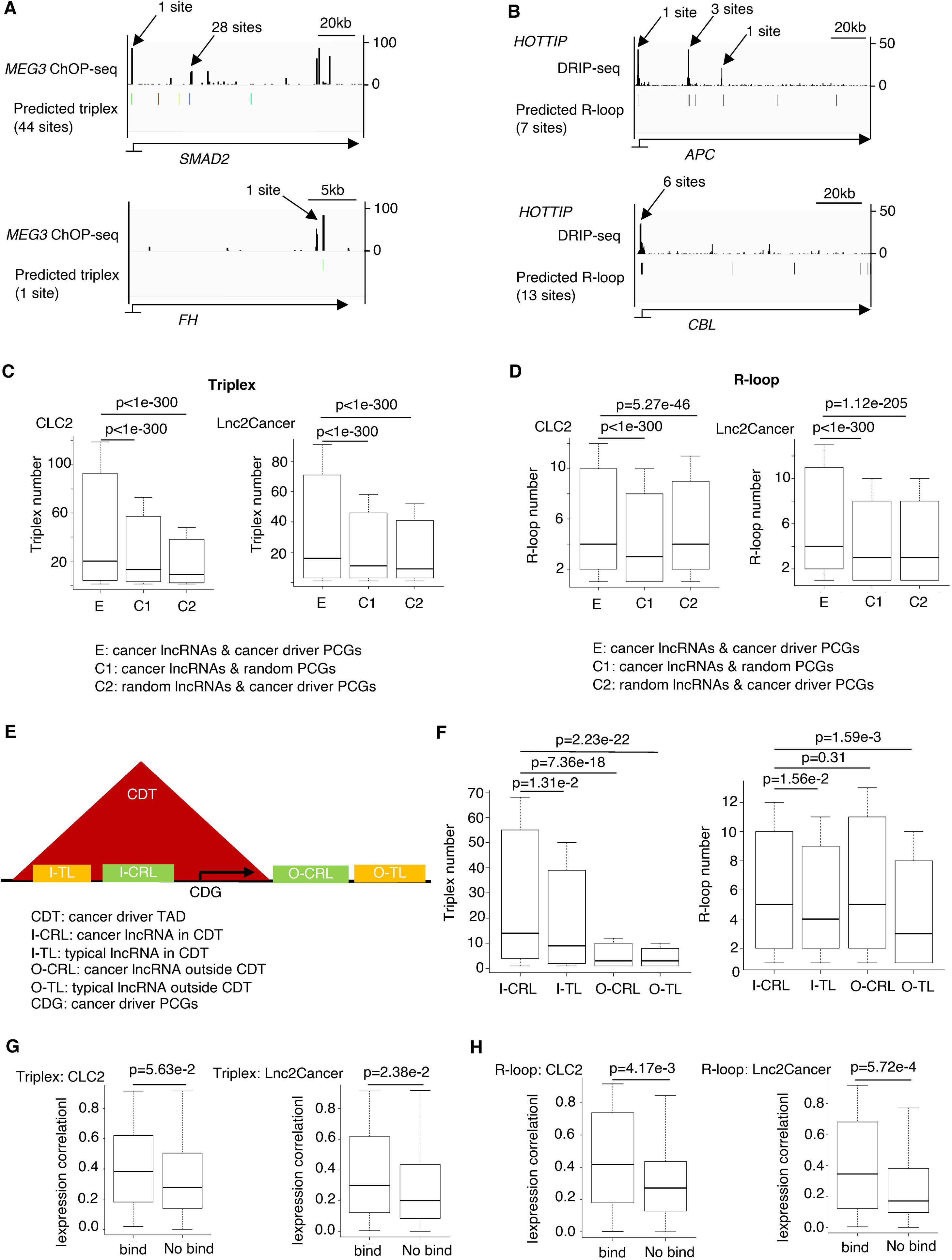
Cancer lncRNAs have a higher binding degree with cancer driver PCGs in the same TAD. **A.** lncRNA *MEG3* ChOP-seq signal and predicted triplex binding sites. **B.** R-loop DRIP-seq signal and predicted R-loop formation sites of lncRNA *HOTTIP*. **C.** Boxplot to show the difference of triplex number between the experimental pair and two control pairs. P values are determined by one tail Wilcoxon test. **D.** Boxplot to show the difference of R-loop number between the experimental pair and two control pairs. P values are determined by one tail Wilcoxon test. **E.** Cartoon pattern to show the definition of the 4 lncRNA types. **F.** Boxplot to show the difference of triplex and R-loop number among I-CRL, I-TL, O-CRL, and O-TL. I-CRL: cancer lncRNAs in CDT. I-TL: typical lncRNAs in CDT. O-CRL: cancer lncRNAs outside CDT. O-TL: typical lncRNAs outside CDT. P values determined by one tail Wilcoxon test. **G.** Boxplot to show the absolute Spearman’s expression correlation between predicted binding pairs and no binding pairs in the same CDT about triplex. P values are determined by one tail Wilcoxon test. **H.** Boxplot to show the absolute Spearman’s expression correlation between predicted binding pairs and no binding pairs in the same CDT about R-loop. P values are determined by one tail Wilcoxon test.

As mentioned previously, the expression levels of the I-CRLs were highly correlated with cancer driver PCGs. Here, we investigated whether the I-CRLs also exhibited a higher affinity for binding to the DNA sequences of cancer driver PCGs. Our results showed that the I-CRLs contained obviously more binding sites in the context of triplex and R-loops structures compared with other groups (Fig. 2E-F). However, there was some difference in O-CRLs between the two structures: the triplex binding sites of the O-CRLs were significantly less than those of the I-CRLs, while the R-loop number of the O-CRLs was almost the same as that of the I-CRLs. We infer that the triplex primarily functions as a *cis*-regulatory element, whereas the R-loops have the capacity to regulate gene expression in both *cis* and *trans* contexts.

To demonstrate the potential of these two RNA-DNA interaction mechanisms to affect gene expression, we next categorized the I-CRLs into two groups: the “bind” group, which comprises I-CRLs that were predicted to bind to cancer driver PCGs in the same CDT, and the “no bind” group, which comprises the remaining I-CRLs that were not predicted to bind to cancer driver PCGs in the same CDT. Whether for triplex or R-loop, the potential binding pairs had higher co-expression tendencies than the no binding pairs (Fig. 2G-H). These outcomes indicate that the two RNA-DNA interaction mechanisms have the potential to function in cancer driver PCG expression regulation by I-CRLs.

### Discovery of pan-cancer driver lncRNAs by constructing CADTAD pipeline

Given that I-CRLs play important roles in cancers, we next identified cancer driver lncRNAs in pan-cancer through the abovementioned characteristics. We collected 139 TAD data from GEO and performed principal component analysis (PCA). We noticed a difference in TAD among normal solid tissue samples, solid tumors, normal blood samples, and hematological malignancies. PC1 distinguished solid tissues and blood samples, while PC2 showed a difference between normal and cancer samples (Fig. 3A, see Methods). Taking into consideration the heterogeneity between solid tumors and hematological malignancies, and given the high prevalence of solid tumors (Chizuka et al. 2006), we exclusively utilized TAD data of solid tumors to identify pan-cancer driver lncRNAs. To accomplish this, we developed an integrated pipeline, CADTAD, to predict the cancer driver lncRNAs potentially involved in regulating the expression of cancer driver PCGs within CDTs. As a result, we identified 256 oncogenic lncRNAs, 177 tumor suppressive lncRNAs, and 75 dual-function lncRNAs (Fig. 3B, Supplemental Data S5).

**Fig. 3.**
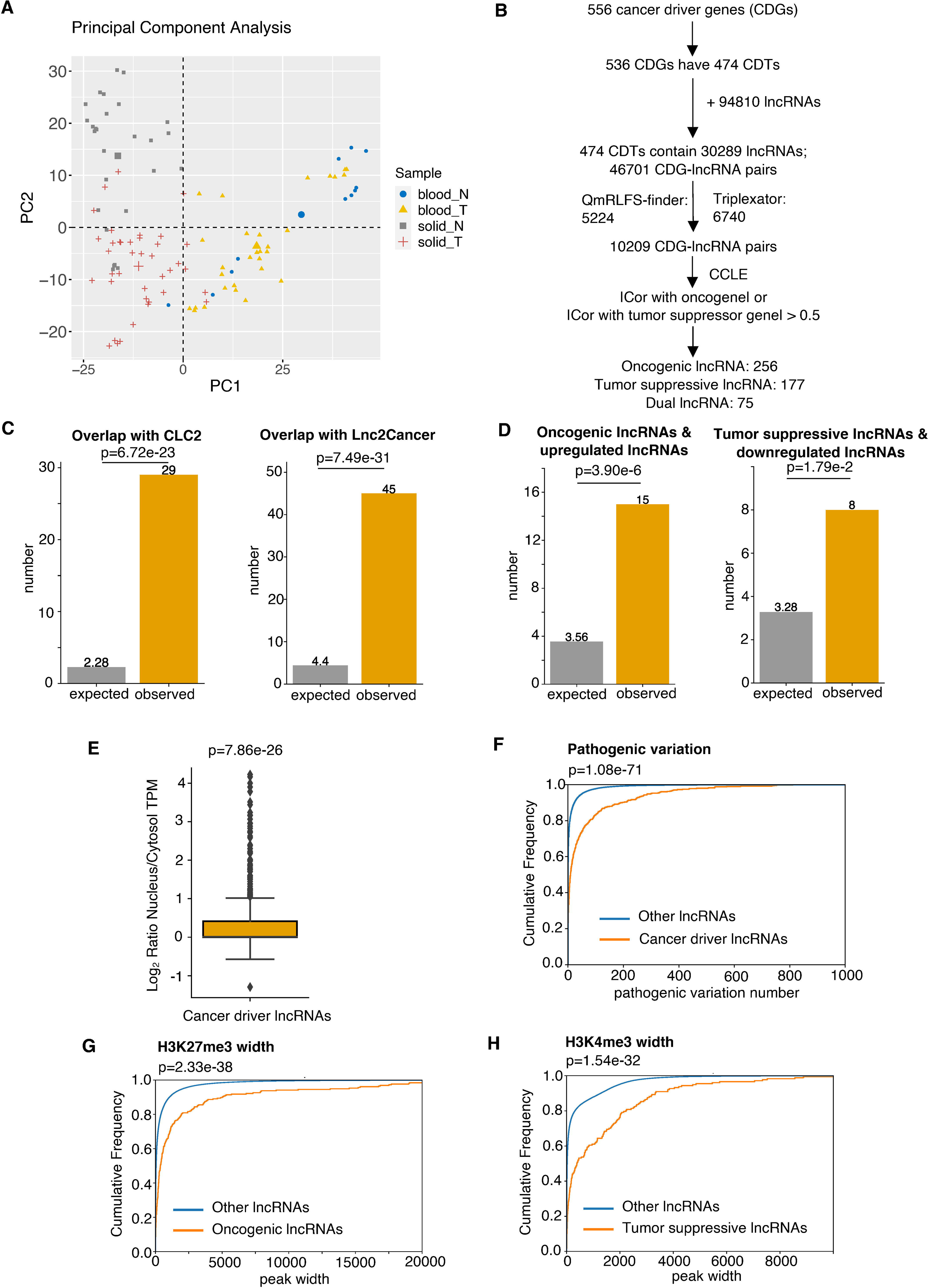
Discovery of potential pan-cancer driver lncRNAs using CADTAD and validation of their cancer-related characteristics using genomics and epigenomics data. **A.** PCA plot to show the difference of TAD among normal blood samples, blood cancers, normal solid samples, and solid tumors. The blue dots represent normal blood samples. The yellow triangles represent blood cancers. The gray rectangles represent normal solid samples. The red crosses represent solid tumors. **B.** The flowchart of identifying potential pan-cancer driver lncRNAs by CADTAD. **C.** Bar plot to show the enrichment of pan-cancer driver lncRNA candidates in CLC2 and Lnc2Cancer datasets. P values are determined by one tail Fisher’s exact test. **D.** Bar plot to show the enrichment of the potential oncogenic lncRNAs in upregulated lncRNAs in cancer, and potential tumor suppressive lncRNAs in downregulated lncRNAs in cancer. P values are determined by one tail Fisher’s exact test. **E.** Boxplot showing the subcellular location tendency of pan-cancer driver lncRNA candidates. P value is determined by one tail *t*-test. **F.** Empirical cumulative distribution plot to show the difference of pathogenic variations between putative driver lncRNAs and the rest of other lncRNAs. P value is determined by one tail Wilcoxon test. **G.** Empirical cumulative distribution plot to show the difference of H3K27me3 width between oncogenic lncRNA candidates and the rest other lncRNAs. P value is determined by one tail Wilcoxon test. **H.** Empirical cumulative distribution plot to show the difference of H3K4me3 width between tumor suppressive lncRNA candidates and the rest other lncRNAs. P value is determined by one tail Wilcoxon test.

To ascertain the tumor-related attributes of the identified cancer driver lncRNAs, we first examined the overlap between potential cancer driver lncRNAs and the cancer lncRNA datasets CLC2 and Lnc2Cancer. The analysis revealed a significant enrichment of candidates in both datasets, indicating that the identified cancer driver lncRNAs retained certain characteristics shared with cancer lncRNAs, thus potentially shedding light on their deeper cancer-related functions (Fig. 3C). We then investigated differentially expressed lncRNAs in pan-cancer (Supplemental Fig. S9A, see Methods), and found potential oncogenic lncRNAs exhibited enrichment in the upregulated lncRNAs, while candidate tumor suppressive lncRNAs were enriched among the downregulated lncRNAs in cancer (Fig. 3D). As expected, we observed no enrichment of potential oncogenic lncRNAs among downregulated lncRNAs, nor did we found an enrichment of candidate tumor suppressive lncRNAs within upregulated lncRNAs (Supplemental Fig. S9B-C). However, potential dual-function lncRNAs were not enriched in either of the differentially expressed lncRNA groups (Supplemental Fig. S9D). Subcellular localization can provide valuable clues about the function of lncRNAs (Bridges et al. 2021), and subcellular localization data analysis revealed that these candidates displayed a higher ratio of nucleus to cytosol expression (Fig. 3E). These findings underscore the predominant role of identified cancer driver lncRNAs within the nucleus.

Traditionally, researchers often rely on mutation data for the discovery of cancer driver PCGs. In our study, we sought to determine whether candidate driver lncRNAs shared similar characteristics. These candidates exhibited a higher number of SNPs classified as pathogenic variation effects annotated from COSMIC compared with the remaining lncRNAs (Fig. 3F), suggesting the pathogenic role of these candidate driver lncRNAs in cancer. We also found that these driver lncRNAs have higher single nucleotide variant (SNV) numbers than the remaining lncRNAs (Supplemental Fig. S9E). The putative oncogenic lncRNAs have a significantly higher copy number variation (CNV) occupation than the non-driver lncRNAs, the tumor suppressive lncRNAs have the contrast phenomenon, and the dual-function lncRNAs have little difference from the non-driver lncRNAs about CNV occupation (Supplemental Fig. S9F). Beyond mutation analysis, various other approaches are used to identify cancer genes through epigenetic markers. For example, broad H3K4me3 marks are associated with tumor-suppressor genes (Chen et al. 2015), while broad H3K27me3 marks are indicative of oncogenes (Zhao et al. 2020). Furthermore, our findings demonstrated that the identified oncogenic lncRNAs possessed wider H3K27me3 marks, whereas candidate tumor suppressive lncRNAs exhibited broader H3K4me3 marks (Fig. 3G-H). Collectively, these results provide compelling evidence of the potential roles in cancer of these identified cancer driver lncRNAs.

### CADTAD analysis for the discovery of putative cancer driver lncRNAs in individual cancers

Given the heterogeneity inherent in tumors, CADTAD was applied to individual cancer types (Supplemental Fig. S10A). To decide which specific cancer to focus on, we considered the distribution of the collected TAD sample numbers across various tumors. Prostate cancer, which ranked at the top, was chosen to identify cancer driver lncRNAs in an individual cancer context (Fig. 4A). We gathered cancer driver PCGs specific to prostate cancer from TUSON (Davoli et al. 2013) and analyzed their distribution (Fig. 4B). We then applied the pipeline to prostate cancer and identified 149 candidate oncogenic lncRNAs, 85 tumor suppressive lncRNAs, and seven dual-function lncRNAs in prostate cancer (Fig. 4C, Supplemental Data S6).

**Fig. 4.**
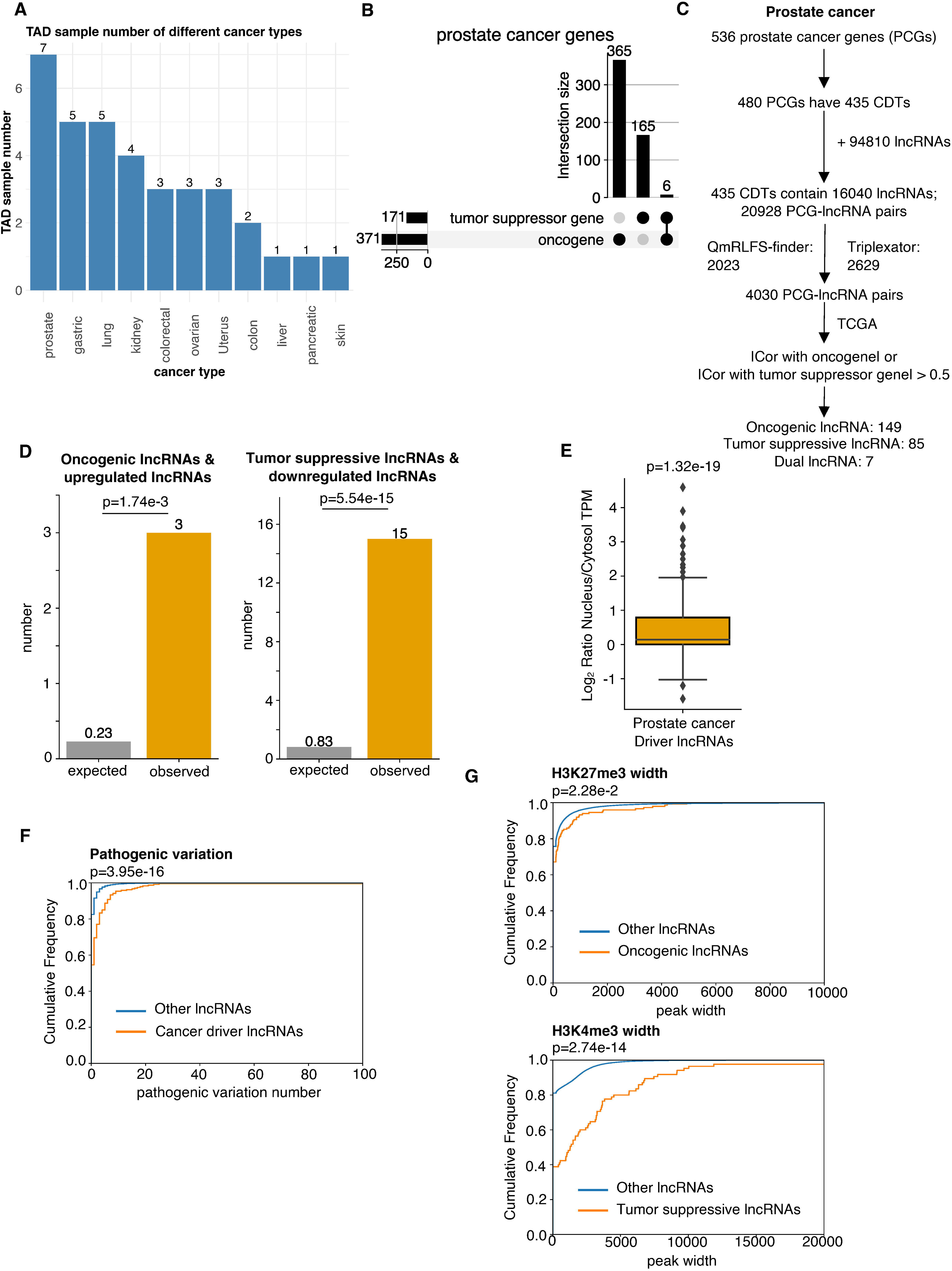
Discovery of potential cancer driver lncRNAs in prostate cancer and validation of their cancer characteristics using genomics and epigenomics data. **A.** Bar plot to show the statistics of sample numbers about TAD data in individual cancers. **B.** Upset plot to show the statistics of tumor suppressor genes and oncogenes in prostate cancer. **C.** The flowchart of identifying potential cancer driver lncRNAs in prostate cancer. **D.** Bar plot to show the enrichment of the putative oncogenic lncRNAs in upregulated lncRNAs in prostate cancer, and putative tumor suppressive lncRNAs in downregulated lncRNAs in prostate cancer. P values are determined by one tail Fisher’s exact test. **E.** Boxplot showing subcellular location tendency of the driver lncRNA candidates about prostate cancer. The P value is determined by one tail *t*-test. **F.** Empirical cumulative distribution plot to show the difference of pathogenic variations between putative cancer driver lncRNAs and the rest of other lncRNAs. P value is determined by one tail Wilcoxon test. **G.** Empirical cumulative distribution plot to show the difference of H3K27me3 width between oncogenic lncRNA candidates and the rest other lncRNAs, and H3K4me3 width between tumor tumor suppressive lncRNA candidates and the rest other lncRNAs. P values are determined by one tail Wilcoxon test.

Leveraging functional data relevant to prostate cancer, like pan-cancer driver lncRNAs, we found that the identified oncogenic lncRNAs in prostate cancer exhibited pronounced enrichment in upregulated lncRNAs, while candidate tumor suppressive lncRNAs were enriched in downregulated lncRNAs (Fig. 4D, Supplemental Fig. S11A). The potential cancer driver lncRNAs in prostate cancer displayed a higher likelihood of being localized in the nucleus (Fig. 4E) and featured a greater number of SNPs classified as pathogenic variations (Fig. 4F). In addition, broader H3K27me3 and H3K4me3 marks were detected in putative oncogenic lncRNAs and tumor suppressive lncRNAs, respectively, in prostate cancer (Fig. 4G).

To further validate the tumor-specific characteristics of potential driver lncRNAs, we randomly selected several candidates in prostate cancer for experimental evaluation. Six prospective oncogenic lncRNAs and four tumor suppressive lncRNAs (Supplemental Fig. S12-13) were disrupted using two pairs of CRISPR-Cas9 guide RNAs. The cutting efficiencies of each pair of guide RNAs were validated through T7 Endonuclease I assay (Supplemental Fig. S14). We also detected the expression level of each candidate lncRNA in edited cells via RT-qPCR. Knocking down five out of six oncogenic candidates significantly inhibited the proliferation of LNCaP cells, as expected (Fig. 5A). LNCaP with decreased HSALNG0082024 showed a tendency toward growth inhibition, although the difference was not significant. Conversely, downreglation of two out of four candidate tumor suppressive lncRNAs significantly promoted cell growth, and the knockdown of HSALNG0124011 had a similar tendency (Fig. 5B). Notably, we observed that the expressions of potential regulated driver PCGs were influenced after the candidate lncRNA knockdown: five oncogenic lncRNA downregulation led to a significant decrease in the expression levels of the associated cancer driver PCGs (Supplemental Fig. S15A), while the knockdown of HSALNG0116774 resulted in an increased mRNA level of its related cancer driver PCG *KIF18B*, as expected (Supplemental Fig. S15B). We also assessed whether these candidates influence cell motility. The downregulation of oncogenic candidates slowed the wound closure in PC-3 prostate cancer cells (Fig. 6A, Supplemental Fig. S16A). However, the cells with the decreased levels of two out of four tumor suppressive lncRNAs closed the wound faster than the control groups (Fig. 6B, Supplemental Fig. S16B).

**Fig. 5.**
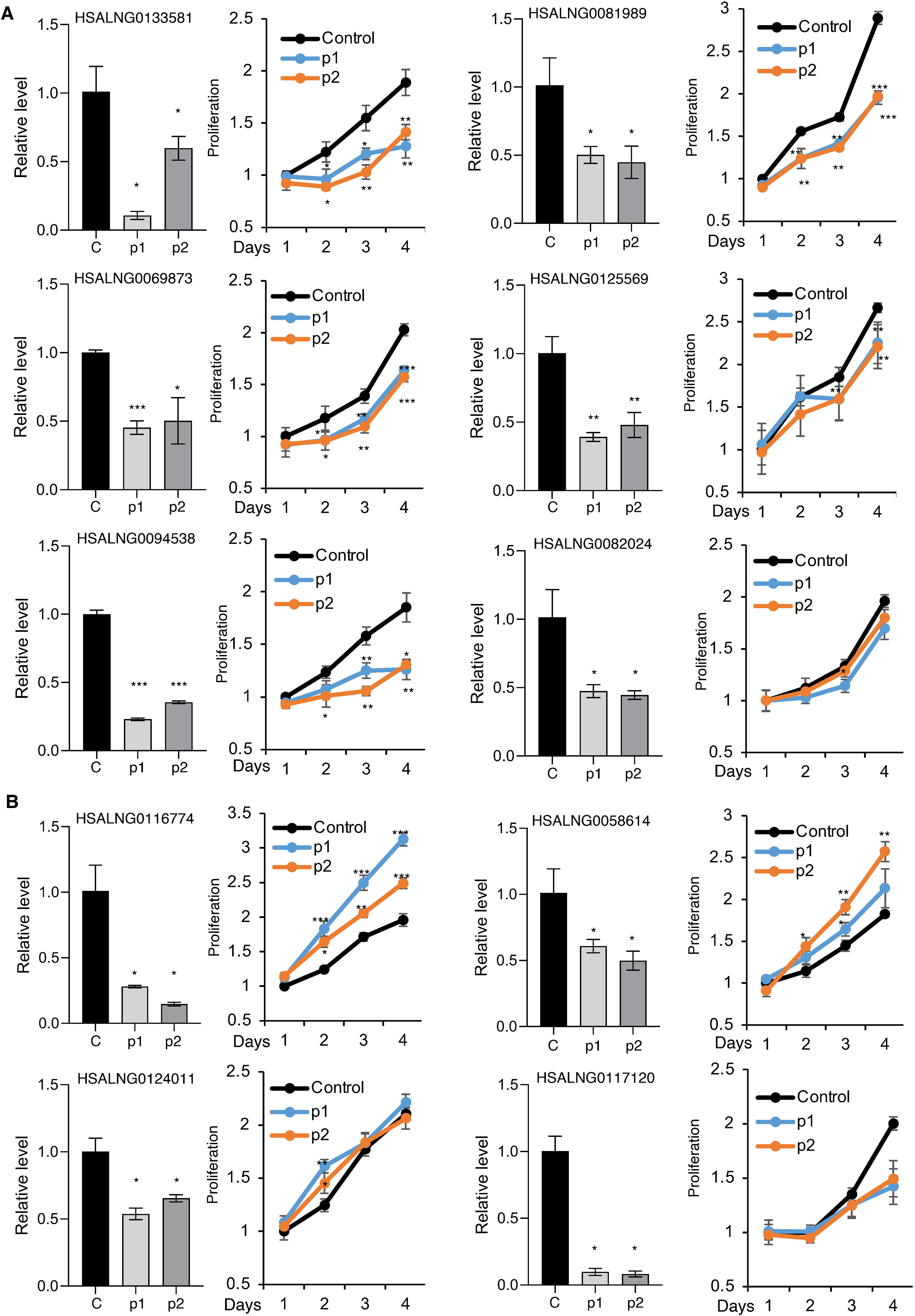
Depletion of identified putative cancer driver lncRNAs impairs or promotes the proliferation of prostate cancer cells. LncRNA expression level and cell proliferation of prostate cancer cell line LNCaP with each putative oncogenic lncRNA (**A**) or tumor suppressive lncRNA (**B**) disrupted by CRISPR-Cas9 guide RNA g1, g2, or under control condition. Bar plot to show lncRNA expression level and line plot to show cell proliferation.

**Fig. 6.**
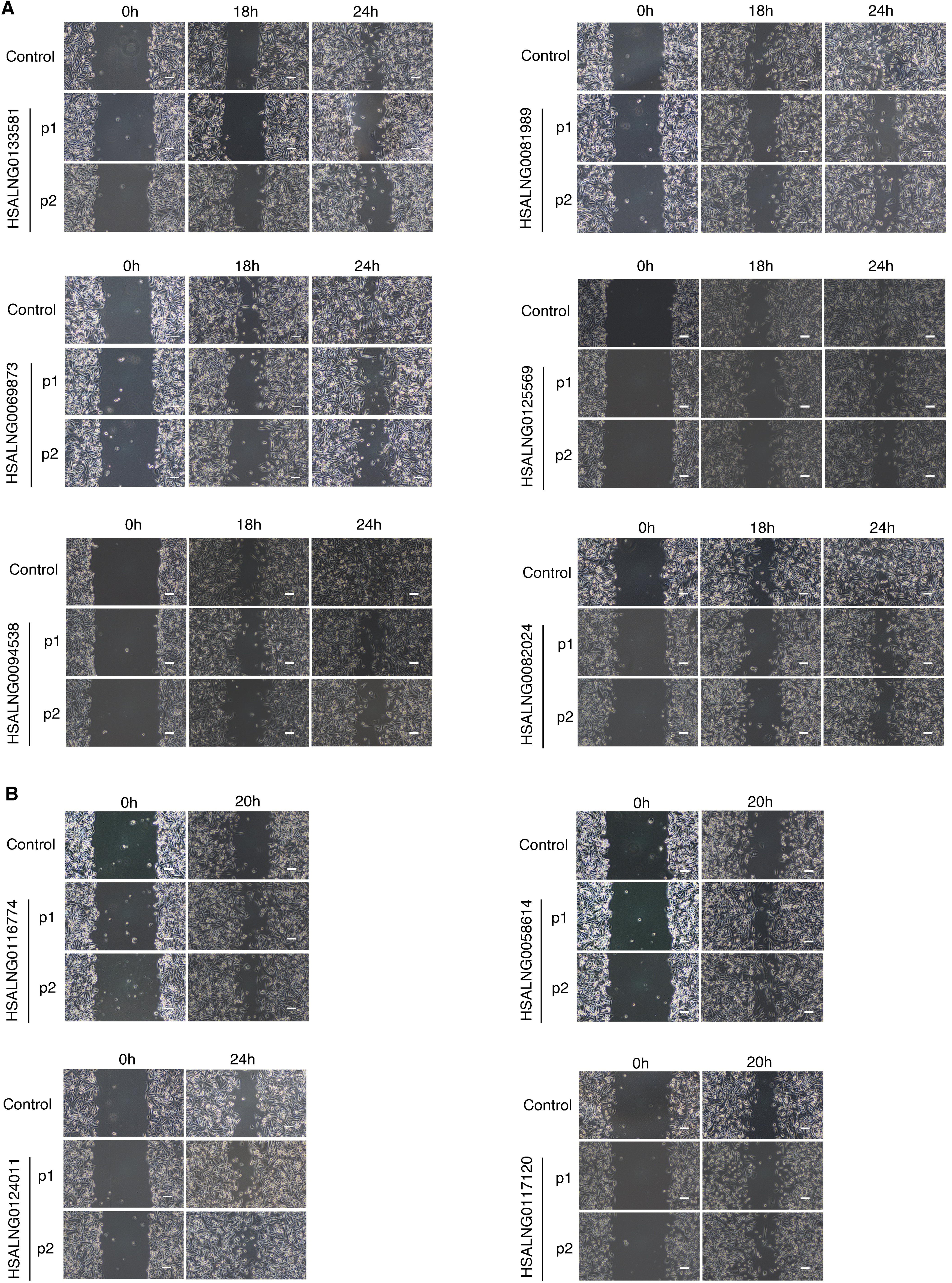
Depletion of the identified putative cancer driver lncRNAs slowed down or hastened the wound closure of prostate cancer cells. Wound healing assay of PC-3 prostate cancer cells with each putative oncogenic lncRNA (**A**) or tumor suppressive lncRNA (**B**) disrupted by CRISPR-Cas9 guide RNA g1, g2, or under control condition.

To exclude the effect caused by CRISPR induced mutations in regulatory elements, we used siRNA pools to target three candidates. The results showed that the downregulation of each lncRNA significantly inhibited the proliferation and motility of prostate cancer cells, consistent with the results using CRISPR (Supplemental Fig. S17A-C). We also found that CRISPR guide RNAs used to target the promoters of HSALNG0082024, HSALNG0094538, HSALNG0116774, and HSALNG0058614 overlapped with the genomic regions of *GATD1*, *MLEC*, *ADAM11*, and *RABGEF1*, respectively. We employed the siRNA pools to target these overlapped genes and figured out whether the alterations of these gene expressions would affect cell proliferation. The downregulation of *MLEC* and *AMAD11* did not significantly affect the proliferation of prostate cancer cells (Supplemental Fig. S18), confirming that the inhibition and promotion of cell growth and motility are specifically due to the knockdown of HSALNG0094538 and HSALNG0116774 by CRISPR, rather than the editing of other genomic regions. The downregulation of *RABGEF1* by siRNA pools significantly inhibited cell proliferation, which was opposite to the effect of HSALNG0058614 knockdown using CRISPR (Supplemental Fig. S18). It further indicated that HSALNG0058614 functions as a tumor suppressive lncRNA in prostate cancer cells. For *GATD1*, its downregulation by siRNA pools obviously inhibited cell proliferation, whereas the knocking down of HSALNG0082024 had little effect on cell proliferation using CRISPR (Fig. 5A, Supplemental Fig. S18).

Survival analysis in prostate adenocarcinoma (PRAD) revealed that higher expression levels of five candidate oncogenic lncRNAs correlated with poorer survival (Supplemental Fig. S19A), while two candidate tumor suppressive lncRNAs exhibited the opposite effect (Supplemental Fig. S19B). Thus, these survival outcomes also highlight the potential of these candidate driver lncRNAs as prognostic markers in prostate cancer.

In summary, these results suggest that CADTAD can effectively identify cancer driver lncRNAs, although its ability to recognize tumor suppressive lncRNAs is not as robust as its detection of oncogenic lncRNAs.

Furthermore, we selected gastric cancer and lung cancer, which ranked at the top based on the distribution of TAD sample numbers, to identify their respective candidate driver lncRNAs (Fig. 4A). Through the CADTAD approach (Supplemental Fig. S10A), we successfully identified 165 candidate oncogenic lncRNAs, 180 tumor suppressive lncRNAs, and 13 dual-function lncRNAs for gastric cancer (Supplemental Fig. S20A-B, Supplemental Data S7). We also identified 101 oncogenic lncRNAs, 278 tumor suppressive lncRNAs, and 19 dual-function lncRNAs for lung cancer (Supplemental Fig. S21A-B, Supplemental Data S8). The cancer driver lncRNAs identified in these two cancer types exhibited consistent cancer-related features, including enrichment of differentially expressed lncRNAs in cancer, nuclear localization, a higher proportion of SNPs classified as pathogenic, and the presence of histone markers associated with cancer genes (Supplemental Fig. S11B-C, Fig. S20C-F, Fig. S21C-F).

These findings underscore the significance of the CDTs combined with expression and interaction as an effective signature for the discovery of cancer driver lncRNAs.

## DISCUSSION

LncRNAs can create an additional regulatory dimension superimposed over the genomic and epigenomic programs by interacting with DNAs, RNAs, or proteins to orchestra gene expression (Herman et al. 2022), defying the conventional central dogma of molecular biology (Chang and Qi 2023). In our investigation into the relationship between cancer lncRNAs and cancer driver PCGs, we are the pioneer to show that cancer lncRNAs are more concentrated in CDTs, representing a novel feature for identifying cancer lncRNAs. Furthermore, cancer lncRNAs in CDTs demonstrated notably higher co-expression levels with cancer driver PCGs in the same TAD, indicating a potential regulatory connection. The co-expression levels are not only determined by distance; after applying three ways to control distance, the CDT feature still significantly affects the levels of expression correlation. In addition, a comprehensive comparison of lncRNA-DNA interactions further accentuated the likelihood that I-CRLs play a role in regulating the expression of cancer driver PCGs. By establishing a direct link between lncRNAs and cancer driver PCGs, we constructed a pipeline CADTAD and identified potential core driver lncRNAs in pan-cancer and across three specific cancer types. These candidates were validated through various bioinformatics function analysis and CRISPR-Cas9 experiments to unveil their potential function in cancer. Notably, CADTAD enables the functional categorization of potential driver lncRNAs into oncogenic, tumor suppressive, and dual-function lncRNAs, representing the first in-silico approach to achieving such classification.

Previous methods for driver lncRNAs discovery primarily relied on a mutation perspective. For example, oncodriveFML (Mularoni et al. 2016) could identify driver mutations in lncRNAs, which is achieved by estimating the cumulative functional impact bias of somatic mutations within tumors. ExInAtor1/2 (Lanzós et al. 2017; Esposito et al. 2023) considered the mutational burden and functional impact to identify cancer driver lncRNAs. However, the identification of driver lncRNAs based on mutation characteristics has yielded relatively low numbers (Mularoni et al. 2016; Lanzós et al. 2017; Esposito et al. 2023). To address this, we advocate approaching cancer driver lncRNAs from a distinct perspective, one that is different from the paradigm of cancer driver PCGs. In this context, we define cancer driver lncRNAs as those that directly regulate the expression of cancer driver PCGs, with the objective of uncovering the lncRNAs with the most significant potential effect on cancer. Although our CADTAD approach does not consider alterations within the cancer driver lncRNAs themselves, such as mutations or epigenetic modifications, it focuses on the crucial task of discerning the driver lncRNAs that govern the core drivers of cancer.

Other studies have also explored lncRNA function in tumors by focusing on lncRNA-mediated regulation. For example, they may identify dysregulated lncRNA-TF-gene triplets in glioblastoma (Li et al. 2016b), or develop methods, such as lncMAP (Li et al. 2018) to reveal distinct types of lncRNA regulatory molecules. DriverLncNet (Zhang et al. 2019b) offers a comprehensive strategy that combines both genomic and regulatory features. This method identifies cancer driver lncRNAs through the analysis of copy number alterations and the construction of regulatory networks. These networks aim to capture key functional effectors, thereby dissecting the functional role of driver lncRNAs. However, cancer lncRNAs are typically classified into two categories based on their functions: oncogenic lncRNAs and tumor suppressive lncRNAs. The current methods for discovering cancer lncRNAs have primarily remained at the association level, lacking the specificity to differentiate their precise roles in either promoting or inhibiting cancer progression. Our pipeline CADTAD can facilitate the distinction of lncRNA-specific functions into either promoting or inhibiting cancer progress.

Nonetheless, our exploration of cancer driver lncRNAs was limited to only three cancer types because of insufficient Hi-C data. In the future, the expansion of Hi-C data across various cancer types will enable the comprehensive discovery of driver lncRNAs in individual cancers, thereby enlarging the cancer driver lncRNA datasets. While the stoichiometric concern regarding lncRNAs and their function is a key debate in the field, the true functional capacity of lncRNAs may depend more on localization in the cell, tissue context, structure, and interaction specificity rather than just their quantity. Here, we used co-expression with cancer driver PCGs to filter cancer driver lncRNAs. This method could exclude lncRNAs that could not be detected in any sample. Therefore, we can confirm that the remaining lncRNAs have the potential to express and bind to DNA. Furthermore, it is essential to emphasize that the current driver lncRNA dataset exclusively focuses on lncRNAs that directly regulate the expression of cancer driver PCGs through lncRNA-DNA interaction prediction. On the one hand, many novel experimental techniques, such as GRID-seq, and RADICL-seq (Mattick et al. 2023), have been developed to detect RNA-chromatin interactions. These high throughput sequencing can be integrated to enhance the accuracy of lncRNA-DNA interactions. On the other hand, lncRNAs have the capacity to influence coding gene expression through various mechanisms, including liquid–liquid phase separation (Adnane et al. 2022) and miRNA sponging (Du et al. 2016). This driver lncRNA dataset represents the core driver dataset within the broader spectrum of all cancer driver lncRNAs capable of directly modulating cancer driver PCG expression. While acknowledging the intricacies of lncRNA regulatory mechanisms, there is an inherent need for a more comprehensive approach to mining driver lncRNAs that encompasses all conceivable lncRNA regulatory pathways. This comprehensive approach will enable us to explore the multifaceted landscape of cancer driver lncRNAs more comprehensively.

In summary, this study finds that CDT is an important feature in identifying cancer lncRNAs and constructs a pipeline CADTAD to identify potential cancer driver lncRNAs based on 3D genome interactions and co-expression features. Notably, it offers a new perspective to discover potential cancer driver lncRNAs. The pipeline CADTAD also holds the potential for broader applications in the study of other diseases.

## METHODS

### Source of datasets analyzed in this project

The lncRNA gene and protein-coding gene (PCG) reference were from LncExpDB (Li et al. 2021) and converted into hg19 using liftOver, which was downloaded from the UCSC Genome Browser website (https://hgdownload.soe.ucsc.edu/downloads.html#utilities_downloads). Thus, we got 94810 lncRNAs and 19840 PCGs. The expression of lncRNAs and PCGs also came from LncExpDB, with combined CCLE and HPA TPM data. FASTA file hg19.fa was from the UCSC (https://hgdownload.soe.ucsc.edu/goldenPath/hg19/bigZips/hg19.fa.gz) and used for later Hi-C mapping and lncRNA-DNA interaction prediction. Cancer lncRNA datasets were collected from CLC2 (Vancura et al. 2021) and Lnc2Cancer3.0 (Gao et al. 2021) and converted into LncBook lncRNA ID according to the LncExpDB reference. We obtained 426 and 822 cancer lncRNAs, respectively, to perform further analysis. Cancer driver PCGs were from the CGC database v.94 (Sondka et al. 2018) and were classified into tumor suppressor genes or oncogenes according to the “Role in Cancer” column containing “TSG” or “oncogene”. If one gene’s “Role in Cancer” column contained both “TSG” and “oncogene”, it would be classified as a dual-function gene. We obtained 556 cancer driver PCGs according to the LncExpDB reference. 1000 oncogenes and 1000 tumor suppressor genes were obtained from a publication (Davoli et al. 2013). A total of 531 genes from the PATHWAY IN CANCER (hsa05200) were downloaded from the Kyoto Encyclopedia of Genes and Genomes (KEGG) database (Kanehisa et al. 2023) (Supplemental Data S1).

Cancer driver PCGs in the individual cancer type were collected from a publication (Davoli et al. 2013). We combined the cancer driver PCG ranking of both individual cancer types and pan-cancer in this publication and took the average ranking. We then selected the cancer driver PCGs for individual cancer types with the top average ranking (<150) to obtain the tumor suppressor genes, oncogenes, and dual-function genes (Fig. 4B, Supplemental Fig. S20A, Fig. S21A, Supplemental Data S2). The statistics of individual cancer genes were calculated using the Python package UpSetPlot (v.0.8.0). RefLnc (Jiang et al. 2019) was used to supplement the TCGA (PRAD, STAD, LUSC, and LUAD) and GTEx projects with the combined FPKM data to obtain the expression of 26494 lncRNAs and 19490 PCGs in a single cancer type. The 17 driver lncRNAs identified by ExInAtor2 were obtained from another study (Esposito et al. 2023).

Hi-C raw data were collected from ENCODE and GEO, in which we obtained a total of 139 samples (Supplemental Data S3). For detailed Hi-C data processing, see Supplemental Methods. Our analyses mostly rely on conserved genomic features, gene annotations, or variant sites that remain largely unchanged between assemblies. Furthermore, any assembly-specific discrepancies would have a minor impact on the biological interpretations derived from our study. Therefore, our findings are not much impacted by the genome assembly used, whether it is GRCh38 or a more recent genome assembly.

### Definition of driver genes

Cancer driver PCGs are those PCGs containing driver mutations, and we collected these cancer driver PCGs from CGC (Sondka et al. 2018) and TUSON (Davoli et al. 2013). As there are some other methods defining cancer driver genes, such as epigenetics (Chen et al. 2015; Zhao et al. 2020), we used a custom method to define the lncRNA that directly regulates cancer driver PCG expression as cancer driver lncRNA.

### Expression correlation calculation

We calculated the absolute pairwise correlation (Spearman’s correlation) between cancer lncRNAs and cancer driver PCGs using the expression values (TPM) from CCLE collected from LncExpDB.

To control the identical expression levels between cancer lncRNAs (CLC2/Lnc2Cancer) and random lncRNAs or between cancer driver PCGs and random PCGs, we stratified the mean expression levels across samples and randomly sampled the same number of no cancer lncRNAs and no cancer PCGs (Supplemental Fig. S1A-C). No cancer lncRNAs were collected from all lncRNAs, excluding CLC2 and Lnc2Cancer, and no cancer PCGs were collected from the PCGs located in typical TADs. All cancer/random lncRNAs and all cancer driver/random PCGs were paired one by one to calculate the expression correlation, so that n lncRNAs and m PCGs had n×m pairs.

### Distance analysis

We took the TSS of PCG as the center, with 40kb as the bin size, and calculated the number of lncRNAs completely falling within this range every time after expanding them using the pybedtools (v. 0.9.0) intersect function with parameter f=1 (Supplemental Fig. S1D). Considering that the lncRNA farthest from the driver PCG in both datasets (CLC2 and Lnc2Cancer) is HSALNG0051162, with a distance of 27663752 bp, we only counted the number within 28 Mb (i.e., we expanded the regions 700 times). We used randomly selected lncRNAs and PCGs in the expression correlation analysis. To analyze in detail the distribution difference between cancer lncRNAs and randomly selected lncRNAs around cancer driver PCGs, we calculated the difference number between the two types of lncRNAs located in this extended area after each enlargement of the region and compared the numbers in these bins using a one-tail paired *t*-test. To obtain the closest lncRNAs for each driver PCGs or randomly selected PCGs, we used all lncRNAs and the closest function in pybedtools (v. 0.9.0). We then overlapped this dataset with cancer lncRNAs or randomly selected lncRNAs to obtain enrichment results using the one-tail Fisher’s exact test.

### Method for defining CDTs and calculating the lncRNA ratio

Using the intersect function in BEDTools (Quinlan and Hall 2010) (v.2.30.0) with the parameter F = 1, we identified TADs as cancer driver TADs (CDTs) if they entirely included at least one cancer driver PCG. The remaining TADs that completely contained non-cancer PCGs, excluding COSMIC genes, top1000 TUSON OGs/TSGs, and the PATHWAY IN CANCER genes, were defined as typical TADs. We randomly selected the same number of typical TADs as CDTs with almost the same length and determined the ratio by dividing the total number of cancer lncRNAs by the summed length of the TAD in each TAD type (Supplemental Fig. S2A). To compare with the neighboring TADs around the CDTs, we also calculated the ratio of up-/down-stream 5 non-CDTs. Once there was one CDT in the up-/down-stream 5 TADs, we skipped it until we found 5 non-CDTs. If the up-/down-stream of CDTs did not have 5 non-CDTs, we assigned a ratio of 0 in the non-TAD region (Supplemental Fig. S2B). We compared the ratios of cancer lncRNAs in CDTs with that in up-/down-stream TADs, typical TADs, and the ratios of random lncRNAs in CDTs using proportion test (Supplemental Table S3).

### Comparison between I-CRL pairs and O-CRL pairs

To compare the levels of co-expression between I-CRL pairs and O-CRL pairs (Fig. 1J), we performed exact matching, which means that each cancer driver PCG in a CDT would have both I-CRL and O-CRL pairs for comparison: we first obtained a pair of I-CRL and cancer driver PCG in the same CDT and then randomly selected an O-CRL after excluding I-CRLs in this CDT to replace the raw I-CRL of the pair (Supplemental Fig. S3A).

For the first try of controlling the distance by choosing the closest O-CRL pairs, we replaced the I-CRL with the O-CRL closest to this cancer driver PCG to obtain the control pairs of O-CRLs and cancer driver PCGs. To obtain the location distribution of the cancer driver PCGs within the CDTs, we scaled each CDT to 0-1 and obtained the relative distance of the cancer driver PCGs in the CDT to calculate the accumulated occurrence frequency in 0.0001 bin size.

For the second try to control the distance by excluding I-CRL pairs that have no O-CRL pairs with similar distances, we selected the O-CRL pairs that have minimum discrepancy of distance with I-CRL pairs and only retained the pairs under a specific distance difference (<50kb) to make the distance similar. In this way, a lot of I-CRL pairs were excluded since some I-CRL pairs couldn’t have O-CRL pairs with similar distances.

For each cancer driver PCG, we also attempted to select lncRNAs that are far away from the cancer driver PCG within the CDT and close to the cancer driver PCG outside the CDT as much as possible to maintain a similar distance to the cancer driver PCGs. After adopting this strategy, we discarded many lncRNAs that were close to the cancer driver PCGs in the CDT but could obtain lncRNAs within or outside CDTs with a similar distance to the cancer driver PCGs.

To assess how CDT and distance contribute to the levels of expression correlation, we fitted a generalized linear regression model (GLM). GLM can isolate the influence of the independent variable of interest (CDT) on the dependent variable (the levels of co-expression) by incorporating other variables into the model as covariates (distance), fixing the effects of other variables. We applied the Fisher transformation to normalize the expression correlation values and then squared them, ensuring they followed a non-central chi-square distribution and fitted a GLM to take the following form

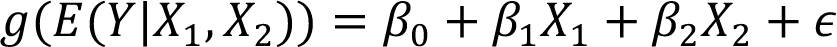

where g(·) is the link function, *Y* is the square of the Fisher transformation of the expression correlations between cancer driver PCGs and cancer lncRNAs, *X*_1_ measures whether the cancer lncRNA is in the CDT (1 indicates the lncRNA is located in the CDT, i.e., I-CRL pair, while 0 indicates that the lncRNA is located outside the CDT, i.e., O-CRL pair), and *X*_2_ quantifies the distance between cancer driver PCGs and cancer lncRNAs. β_0_ is the slope, and ϵ is a random error term. β_1_ is the marginal effect of *X*_1_ on *Y* when *X*_2_ is fixed, meaning that distance is controlled to observe how CDT affects the levels of co-expression. We fitted the model using statsmodels (v. 0.13.2)

### Construction of a cancer-related lncRNA-mRNA network and the definition of lncRNA centrality

We used the concept of weighted gene co-expression network analysis (WGCNA) to construct a cancer-related lncRNA-mRNA network (Zhang and Horvath 2005; Langfelder and Horvath 2008). We calculated the Pearson’s correlation between the expression of all lncRNAs and all mRNAs to obtain a similarity matrix and then transferred it to an adjacency matrix by raising the similarity to power. Next, we used the scale-free topology criterion for soft thresholding, and the minimum scale-free topology fitting index *R*^2^ was set to 0.9; thus, we obtained a soft threshold of 6 (Supplemental Fig. S7A-B). To identify cancer-related functions, we defined important genes as those containing cancer genes from the CGC database v. 94, the top 500 oncogenes and tumor suppressor genes from TUSON, and the PATHWAY IN CANCER genes from KEGG. We used power 6 to calculate the network between all lncRNAs and important genes and summed the values of one lncRNA with all important genes to define lncRNA centrality. As we only focused on the rank of I-CRLs and O-CRLs in all lncRNAs, we performed GSEA to compare them using the Python package GSEApy (Fang et al. 2023) (v. 1.0.5).

### lncRNA-DNA interaction analysis

We knew of two mechanisms through which lncRNA could interact with DNA: triplex and R-loop. We reviewed all relevant prediction techniques and chose the classic methods Triplextor (Buske et al. 2012) and QmRLFS-finder (Jenjaroenpun et al. 2015) to predict the lncRNA-DNA interaction. Before prediction, we obtained the sequences of lncRNAs and PCGs from hg19.fa using the getfasta function in BEDTools (v. 2.30.0) (Quinlan and Hall 2010). For the triplex prediction, we predicted the number between lncRNAs and PCGs using the default parameter. For the R-loop prediction, we first predicted the R-loop structure in each lncRNA and then counted all the matched numbers of paired PCGs whose sense or antisense strand had a RIZ sequence (Luo et al. 2022), as the RIZ sequence is very important for forming a R-loop. lncRNA *MEG3* ChOP-seq data was collected from a recent publication (Mondal et al. 2015). The DRIP-seq data on the lncRNA *HOTTIP* R-loop formation analysis was obtained from a study (Luo et al. 2022).

### Construction of a pipeline CADTAD to discover cancer driver lncRNAs

We constructed a pipeline CADTAD to discover cancer driver lncRNAs in pan-cancer and individual cancer types (Fig. 3B, Fig. 4C, Supplemental Fig. S10, S20B, Fig. S21B). First, we obtained all TADs completely covering cancer driver PCGs, and then we merged all intersected TADs to define the CDT using the intersect and merge function in pybedtools (v. 0.9.0) (Quinlan and Hall 2010; Dale et al. 2011). We then discovered the lncRNAs located in these CDTs with a reference of all lncRNAs and matched these lncRNAs with the cancer driver PCGs in the same CDT. Subsequently, we predicted the potential lncRNA-DNA interaction pairs of matched lncRNA-cancer driver PCG pairs using the Triplexator and QmRLF-finder as soon as one of the methods predicted. Finally, we used tumor samples’ RNA-seq expression data to calculate the expression correlation of the remaining lncRNA-cancer driver PCG pairs and obtained lncRNAs with absolute correlation values greater than 0.5 as cancer driver lncRNAs. We considered driver lncRNAs whose expression correlation values were only positive with oncogenes and/or negative with tumor suppressor genes as oncogenic lncRNAs, only positive with tumor suppressor genes and/or negative with oncogenes as tumor suppressive lncRNAs, and that had the above two cases as dual-function lncRNAs. We then applied CADTAD to pan-cancer and individual cancers to identify cancer driver lncRNAs (Supplemental Data S5-8).

### Bioinformatics methods related to cancer characteristics verification

To obtain upregulated and downregulated lncRNAs in cancer, we compared cancer samples with normal samples using the Wilcoxon test and calculated the fold change. lncRNAs whose p values were less than 0.05 and |logFC| were more than 1 were considered as differential expressions. We calculated the enrichment p-values using the one-tail Fisher’s exact test and the expected values by multiplying the total number of two datasets, which was then divided by the total number of background gene sets. The nucleus-and cytosol-featured lncRNAs were taken from LncExpDB (Li et al. 2021). The data were used to calculate the ratio of the nucleus to cytosol expression for each lncRNA to explore the subcellular location of the lncRNAs. Variation data were extracted from LncBook2.0 (Li et al. 2022). For comparison, we used only the COSMIC Variant Effect, which was pathogenic. The SNV and CNV data were collected from PCAWG (The ICGC/TCGA Pan-Cancer Analysis of Whole Genomes Consortium 2020). The H3K4me3 and H3K27me3 peak files were downloaded from ENCODE (Supplemental Data S4). We calculated each lncRNA’s peak width by adding the lengths of the intersected peaks. Survival data were taken from UCSC Xena (Caicedo et al. 2020), and survival analyses were conducted using the Python package lifelines (v.0.27.7).

### CRISPR gRNA and lentiviral vector design

For each candidate lncRNA, two pairs of guide RNAs were designed to delete about 4 kb of the promoter region, defined as from 5 kb upstream to 1 kb downstream of transcription site (TSS). A pair of two non-targeting sgRNAs was used as the control. The sequences of each guide RNAs were listed in Supplemental Table S7.

The oligos for each target site were annealed by heating to 95°C for 5 minutes and cooling to 85 °C at 2 °C per second and then further down to 25 °C at 0.1 °C per second. The annealed oligos were cloned into lentiCRISPR v2 plasmid and sequenced to confirm successful ligation.

### Cell culture and lentiviral transduction

LNCaP, PC-3, and HEK293T were purchased from ATCC. All cells were maintained in DMEM medium supplemented with 10% FBS and cultured in 5% CO2. These cell lines were mycoplasma negative during routine tests.

Lentiviral was packaged through the co-transfection of the second packaging vectors pMD2.G and psPAX2 into HEK293T cells. After 48 hours of transfection, the medium was harvested and centrifuged at 800 × g for 10 minutes to remove cell debris. The supernatant was used for infection when LNCaP or PC-3 cells were grown to 70% confluence. The medium was changed 24 hours after viral transduction, and the infected cells were incubated for another 24 hours before puromycin selection. The positive cells were collected for the cell proliferation assay, wound heal assay, and total RNA and genomic DNA extraction.

### PCR and RT-qPCR verification of CRISPR-Cas9-induced deletion at the candidate lncRNAs promoter

Mutagenesis of candidate lncRNAs was verified through PCR using primers flanking the deletion regions. In control cells, the PCR fragments are the full length between the pair of primers. But in the edited cells, this region was edited by the pair of CRISPR guide RNA, resulting in the shorter PCR fragments. RT-qPCR was used to detect the RNA levels of the lncRNA candidates after CRISPR-Cas9 editing. The total RNA of edited cells was extracted and converted into cDNA. Then, qPCR was performed with ChamQ Universal SYBR qPCR Master Mix (Vazyme, Nanjing, China). The primers used for PCR and RT-qPCR of candidate lncRNAs were listed in Supplemental Table S8. The primers used for RT-qPCR of associated driver PCGs were listed in Supplemental Table S9. The primers used for RT-qPCR of overlapped PCGs were listed in Supplemental Table S10.

### Cell proliferation and wound healing assay

The LNCaP cells were transduced by virus containing the pair of guide RNAs and then selected by 1 µg/mL puromycin. After selection, the LNCaP cells were seeded in 96-well plates at a density of 6000 cells per well and allowed to attach for 24 hours. For every time point, the cells were added with CCK8 reagent at a ratio of 1:10. After 1.5 hours of incubation at 37°C, the absorbance of 450 nm wavelength was measured to evaluate the cell proliferation rate. The CRISPR-Cas9 guide RNA edited PC-3 cells were inoculated into a 6-well plate and the wound was created until the cell confluence arrived at 90%. The photos were taken at the indicated time by using microscope. The wound closure rate was quantified as percentages by using Image J. LNCaP and PC-3 cells transfected with siRNAs targeting each indicated lncRNA were evaluated in terms of their proliferation and migration abilities.

### Statistical analysis

The one-tail Wilcoxon test was used to compare the feature values of different lncRNA categories. Functional dataset enrichment analyses were performed by using the one-tail Fisher’s exact test.

### Software availability

The CADTAD pipeline is available at GitHub (https://github.com/starrzy/CADTAD) and as Supplemental Code.

## COMPETING INTEREST STATEMENT

The authors declare that they have no competing interests.

## ACKNOWLEDGMENTS

We are grateful to Lina Ma, and Zhao Li for their helpful support of lncRNA data. This work was supported by High-performance Computing Platform of Peking University.

## Author Contributions

Dongyu Zhao conceived the project. Dongyu Zhao and Ziyan Rao designed the bioinformatics analyses and interpreted the results. Ziyan Rao performed the bioinformatics analyses, with assistance from Shaodong Huang, Chenyang Wu, and Yuheng Zhou. Min Zhang designed and performed the experiments, and interpreted the results, with assistance from Weijie Zhang, and Xia Lin. Dongyu Zhao, Min Zhang, and Ziyan Rao wrote the manuscript.

## Funding

This work was supported by the National Key R&D Program of China (2021YFF1201100), the National Natural Science Foundation of China (NSFC, 32270603) and the Fundamental Research Funds for the Central Universities (BMU2021YJ057).

## Notes

https://genome.cshlp.org/content/early/2025/05/27/gr.280235.124

### Competing Interest Statement

The authors have declared no competing interest.

### Summary of Updates

Enhanced Persuasiveness of Analyses: To make the comparison results of co-expression and distance analysis more convincing, we corrected some artifacts by randomly selecting expression-matched no-cancer PCGs and lncRNAs with the same number of cancer driver ones, ensuring a more robust and unbiased validation of our conclusions. Refinement of CDT validation: To control the influence of distance on the CDT, we implemented advanced sampling methods and generalized linear regression to validate the effectiveness of the CDT. Validation of associated cancer driver PCGs: To validate the relationship between the identified cancer driver lncRNAs and their related cancer driver PCG, we conducted additional experiments to evaluate the expression levels of the associated cancer driver PCGs after knockdown of the candidate lncRNAs. siRNA-based functional assays further confirmed the function of these lncRNA candidates in cancer cell line. Improved Clarity and Writing: We revised the manuscript extensively to refine the conclusions, provide more detailed description of the methods, and polish the whole manuscript with the help of a native speaker.

## REFERENCES

1. Adnane S, Marino A, Leucci E. 2022. LncRNAs in human cancers: signal from noise. Trends Cell Biol 32: 565–573. 10.1016/j.tcb.2022.01.006.

2. Arab K, Karaulanov E, Musheev M, Trnka P, Schäfer A, Grummt I, Niehrs C. 2019. GADD45A binds R-loops and recruits TET1 to CpG island promoters. Nat Genet 51: 217–223. 10.1038/s41588-018-0306-6.

3. Ariel F, Lucero L, Christ A, Mammarella MF, Jegu T, Veluchamy A, Mariappan K, Latrasse D, Blein T, Liu C, et al. 2020. R-Loop Mediated trans Action of the APOLO Long Noncoding RNA. Mol Cell 77: 1055–1065.e4. 10.1016/j.molcel.2019.12.015.

4. Bhan A, Soleimani M, Mandal SS. 2017. Long noncoding RNA and cancer: A new paradigm. Cancer Res 77: 3965–3981.

5. Bridges MC, Daulagala AC, Kourtidis A. 2021. LNCcation: lncRNA localization and function. J Cell Biol 220: 1–17.

6. Buske FA, Bauer DC, Mattick JS, Bailey TL. 2012. Triplexator: Detecting nucleic acid triple helices in genomic and transcriptomic data. Genome Res 22: 1372–1381.

7. Caicedo HH, Hashimoto DA, Caicedo JC, Pentland A, Pisano GP. 2020. Visualizing and interpreting cancer genomics data via the Xena platform. Nat Biotechnol 38: 669–673.

8. Chang HY, Qi LS. 2023. Reversing the Central Dogma: RNA-guided control of DNA in epigenetics and genome editing. Mol Cell 83: 442–451. 10.1016/j.molcel.2023.01.010.

9. Chen K, Chen Z, Wu D, Zhang L, Lin X, Su J, Rodriguez B, Xi Y, Xia Z, Chen X, et al. 2015. Broad H3K4me3 is associated with increased transcription elongation and enhancer activity at tumor-suppressor genes. Nat Genet 47: 1149–1157.

10. Chiu HS, Somvanshi S, Patel E, Chen TW, Singh VP, Zorman B, Patil SL, Pan Y, Chatterjee SS, Caesar-Johnson SJ, et al. 2018. Pan-Cancer Analysis of lncRNA Regulation Supports Their Targeting of Cancer Genes in Each Tumor Context. Cell Rep 23: 297–312.e12.

11. Chizuka A, Suda M, Shibata T, Kusumi E, Hori A, Hamaki T, Kodama Y, Horigome K, Kishi Y, Kobayashi K, et al. 2006. Difference between hematological malignancy and Solid tumor research articles published in four major medical journals. Leukemia 20: 1655–1657.

12. Choudhury SR, Dutta S, Bhaduri U, Rao MRS. 2021. LncRNA Hmrhl regulates expression of cancer related genes in chronic myelogenous leukemia through chromatin association. NAR Cancer 3: 1–26.

13. Dale RK, Pedersen BS, Quinlan AR. 2011. Pybedtools : a flexible Python library for manipulating genomic datasets and annotations. 27: 3423–3424.

14. Davoli T, Xu AW, Mengwasser KE, Sack LM, Yoon JC, Park PJ, Elledge SJ. 2013. Cumulative haploinsufficiency and triplosensitivity drive aneuploidy patterns and shape the cancer genome. Cell 155: 948. 10.1016/j.cell.2013.10.011.

15. Derrien T, Johnson R, Bussotti G, Tanzer A, Djebali S, Tilgner H, Guernec G, Martin D, Merkel A, Knowles DG, et al. 2012. The GENCODE v7 catalog of human long noncoding RNAs: Analysis of their gene structure, evolution, and expression. Genome Res 22: 1775–1789.

16. Du Z, Sun T, Hacisuleyman E, Fei T, Wang X, Brown M, Rinn JL, Lee MGS, Chen Y, Kantoff PW, et al. 2016. Integrative analyses reveal a long noncoding RNA-mediated sponge regulatory network in prostate cancer. Nat Commun 7.

17. Esposito R, Lanzós A, Uroda T, Ramnarayanan S, Büchi I, Polidori T, Guillen-Ramirez H, Mihaljevic A, Merlin BM, Mela L, et al. 2023. Tumour mutations in long noncoding RNAs enhance cell fitness. Nat Commun 14: 3342. https://www.nature.com/articles/s41467-023-39160-7.

18. Fang Z, Liu X, Peltz G. 2023. GSEApy: a comprehensive package for performing gene set enrichment analysis in Python. Bioinformatics 39: 1–3.

19. Gao Y, Shang S, Guo S, Li X, Zhou H, Liu H, Sun Y, Wang J, Wang P, Zhi H, et al. 2021. Lnc2Cancer 3.0: An updated resource for experimentally supported lncRNA/circRNA cancer associations and web tools based on RNA-seq and scRNA-seq data. Nucleic Acids Res 49: D1251–D1258.

20. Gil N, Ulitsky I. 2020. Regulation of gene expression by cis-acting long non-coding RNAs. Nat Rev Genet 21: 102–117. 10.1038/s41576-019-0184-5.

21. Gupta RA, Shah N, Wang KC, Kim J, Horlings HM, Wong DJ, Tsai MC, Hung T, Argani P, Rinn JL, et al. 2010. Long non-coding RNA HOTAIR reprograms chromatin state to promote cancer metastasis. Nature 464: 1071–1076.

22. Herman AB, Tsitsipatis D, Gorospe M. 2022. Integrated lncRNA function upon genomic and epigenomic regulation. Mol Cell 82: 2252–2266. 10.1016/j.molcel.2022.05.027.

23. Huarte M. 2015. The emerging role of lncRNAs in cancer. Nat Med 21: 1253–1261.

24. Jenjaroenpun P, Wongsurawat T, Yenamandra SP, Kuznetsov VA. 2015. QmRLFS-finder: A model, web server and stand-alone tool for prediction and analysis of R-loop forming sequences. Nucleic Acids Res 43: W527–W534.

25. Jiang S, Cheng SJ, Ren LC, Wang Q, Kang YJ, Ding Y, Hou M, Yang XX, Lin Y, Liang N, et al. 2019. An expanded landscape of human long noncoding RNA. Nucleic Acids Res 47: 7842–7856.

26. Kanehisa M, Furumichi M, Sato Y, Kawashima M, Ishiguro-Watanabe M. 2023. KEGG for taxonomy-based analysis of pathways and genomes. Nucleic Acids Res 51: D587–D592.

27. Kotake Y, Nakagawa T, Kitagawa K, Suzuki S, Liu N, Kitagawa M, Xiong Y. 2011. Long non-coding RNA ANRIL is required for the PRC2 recruitment to and silencing of p15 INK4B tumor suppressor gene. Oncogene 30: 1956–1962.

28. Langfelder P, Horvath S. 2008. WGCNA: An R package for weighted correlation network analysis. BMC Bioinformatics 9.

29. Lanzós A, Carlevaro-Fita J, Mularoni L, Reverter F, Palumbo E, Guigó R, Johnson R. 2017. Discovery of Cancer Driver Long Noncoding RNAs across 1112 Tumour Genomes: New Candidates and Distinguishing Features. Sci Rep 7: 1–16. 10.1038/srep41544.

30. Leisegang MS, Warwick T, Stötzel J, Brandes RP. 2024. RNA-DNA triplexes: molecular mechanisms and functional relevance. Trends Biochem Sci **xx**: 1–13. 10.1016/j.tibs.2024.03.009.

31. Li Y, Li L, Wang Z, Pan T, Sahni N, Jin X, Wang G, Li J, Zheng X, Zhang Y, et al. 2018. LncMAP: Pan-cancer Atlas of long noncoding RNA-mediated transcriptional network perturbations. Nucleic Acids Res 46: 1113–1123.

32. Li Y, Syed J, Sugiyama H. 2016a. RNA-DNA Triplex Formation by Long Noncoding RNAs. Cell Chem Biol 23: 1325–1333. 10.1016/j.chembiol.2016.09.011.

33. Li Y, Wang Z, Wang Y, Zhao Z, Zhang J, Lu J, Xu J, Li X. 2016b. Identification and characterization of lncRNA mediated transcriptional dysregulation dictates lncRNA roles in glioblastoma. Oncotarget 7: 45027–45041.

34. Li Z, Liu L, Feng C, Qin Y, Xiao J, Zhang Z, Ma L. 2022. LncBook 2.0: integrating human long non-coding RNAs with multi-omics annotations. Nucleic Acids Res 51: 186–191.

35. Li Z, Liu L, Jiang S, Li Q, Feng C, Du Q, Zou D, Xiao J, Zhang Z, Ma L. 2021. LncExpDB: An expression database of human long non-coding RNAs. Nucleic Acids Res 49: D962–D968.

36. Liu Z, Zhang Y, Han X, Li C, Yang X, Gao J, Xie G, Du N. 2020. Identifying Cancer-Related lncRNAs Based on a Convolutional Neural Network. Front Cell Dev Biol 8: 1–7.

37. Luo H, Zhu G, Eshelman MA, Fung TK, Lai Q, Wang F, Zeisig BB, Lesperance J, Ma X, Chen S, et al. 2022. HOTTIP-dependent R-loop formation regulates CTCF boundary activity and TAD integrity in leukemia. Mol Cell 82: 833–851.e11. 10.1016/j.molcel.2022.01.014.

38. Martínez-Jiménez F, Muiños F, Sentís I, Deu-Pons J, Reyes-Salazar I, Arnedo-Pac C, Mularoni L, Pich O, Bonet J, Kranas H, et al. 2020. A compendium of mutational cancer driver genes. Nat Rev Cancer 20: 555–572. 10.1038/s41568-020-0290-x.

39. Mattick JS, Amaral PP, Carninci P, Carpenter S, Chang HY, Chen LL, Chen R, Dean C, Dinger ME, Fitzgerald KA, et al. 2023. Long non-coding RNAs: definitions, functions, challenges and recommendations. Nat Rev Mol Cell Biol.

40. Melé M, Rinn JL. 2016. “Cat’s Cradling” the 3D Genome by the Act of LncRNA Transcription. Mol Cell 62: 657–664.

41. Mondal T, Subhash S, Vaid R, Enroth S, Uday S, Reinius B, Mitra S, Mohammed A, James AR, Hoberg E, et al. 2015. MEG3 long noncoding RNA regulates the TGF-β pathway genes through formation of RNA-DNA triplex structures. Nat Commun 6.

42. Mularoni L, Sabarinathan R, Deu-Pons J, Gonzalez-Perez A, López-Bigas N. 2016. OncodriveFML: A general framework to identify coding and non-coding regions with cancer driver mutations. Genome Biol 17: 1–13. 10.1186/s13059-016-0994-0.

43. Niehrs C, Luke B. 2020. Regulatory R-loops as facilitators of gene expression and genome stability. Nat Rev Mol Cell Biol 21: 167–178. 10.1038/s41580-019-0206-3.

44. Nora EP, Lajoie BR, Schulz EG, Giorgetti L, Okamoto I, Servant N, Piolot T, Van Berkum NL, Meisig J, Sedat J, et al. 2012. Spatial partitioning of the regulatory landscape of the X-inactivation centre. Nature 485: 381–385.

45. Poliseno L, Salmena L, Zhang J, Carver B, Haveman WJ, Pandolfi PP. 2010. A coding-independent function of gene and pseudogene mRNAs regulates tumour biology. Nature 465: 1033–1038. 10.1038/nature09144.

46. Quinlan AR, Hall IM. 2010. BEDTools : a flexible suite of utilities for comparing genomic features. 26: 841–842.

47. Quinodoz SA, Jachowicz JW, Bhat P, Ollikainen N, Banerjee AK, Goronzy IN, Blanco MR, Chovanec P, Chow A, Markaki Y, et al. 2021. RNA promotes the formation of spatial compartments in the nucleus. Cell 184: 5775–5790.e30. 10.1016/j.cell.2021.10.014.

48. Rinn JL, Kertesz M, Wang JK, Squazzo SL, Xu X, Brugmann SA, Goodnough LH, Helms JA, Farnham PJ, Segal E, et al. 2007. Functional Demarcation of Active and Silent Chromatin Domains in Human HOX Loci by Noncoding RNAs. Cell 129: 1311–1323.

49. Schmitt AM, Chang HY. 2016. Long Noncoding RNAs in Cancer Pathways. Cancer Cell 29: 452–463. 10.1016/j.ccell.2016.03.010.

50. Sondka Z, Bamford S, Cole CG, Ward SA, Dunham I, Forbes SA. 2018. The COSMIC Cancer Gene Census: describing genetic dysfunction across all human cancers. Nat Rev Cancer 18: 696–705. 10.1038/s41568-018-0060-1.

51. Statello L, Guo CJ, Chen LL, Huarte M. 2021. Gene regulation by long non-coding RNAs and its biological functions. Nat Rev Mol Cell Biol 22: 96–118. 10.1038/s41580-020-00315-9.

52. The ICGC/TCGA Pan-Cancer Analysis of Whole Genomes Consortium. 2020. Pan-cancer analysis of whole genomes. Nature 578: 82–93.

53. Tseng YY, Moriarity BS, Gong W, Akiyama R, Tiwari A, Kawakami H, Ronning P, Reuland B, Guenther K, Beadnell TC, et al. 2014. PVT1 dependence in cancer with MYC copy-number increase. Nature 512: 82–86.

54. Valton AL, Dekker J. 2016. TAD disruption as oncogenic driver. Curr Opin Genet Dev 36: 34–40. 10.1016/j.gde.2016.03.008.

55. Vancura A, Lanzós A, Bosch-Guiteras N, Esteban MT, Gutierrez AH, Haefliger S, Johnson R. 2021. Cancer LncRNA Census 2 (CLC2): an enhanced resource reveals clinical features of cancer lncRNAs. NAR Cancer 3: 1–15.

56. Werner MS, Sullivan MA, Shah RN, Nadadur RD, Grzybowski AT, Galat V, Moskowitz IP, Ruthenburg AJ. 2017. Chromatin-enriched lncRNAs can act as cell-type specific activators of proximal gene transcription. Nat Struct Mol Biol 24: 596–603.

57. Yu W, Gius D, Onyango P, Muldoon-Jacobs K, Karp J, Feinberg AP, Cui H. 2008. Epigenetic silencing of tumour suppressor gene p15 by its antisense RNA. Nature 451: 202–206.

58. Yuan L, Zhao J, Sun T, Shen Z. 2021. A machine learning framework that integrates multi-omics data predicts cancer-related LncRNAs. BMC Bioinformatics 22: 1–18. 10.1186/s12859-021-04256-8.

59. Zhang B, Horvath S. 2005. A general framework for weighted gene co-expression network analysis. Stat Appl Genet Mol Biol 4.

60. Zhang J, Gao Y, Wang P, Zhi H, Zhang Y, Guo M, Yue M, Li X, Zhou D, Wang Y, et al. 2020. CLING: Candidate Cancer-Related lncRNA Prioritization via Integrating Multiple Biological Networks. Front Bioeng Biotechnol 8: 1–12.

61. Zhang J, Le TD, Liu L, Li J. 2018a. Inferring and analyzing module-specific lncRNA-mRNA causal regulatory networks in human cancer. Brief Bioinform 20: 1403–1419.

62. Zhang X, Li T, Wang J, Li J, Chen L, Liu C. 2019a. Identification of cancer-related long non-coding RNAs using XGboost with high accuracy. Front Genet 10: 1–14.

63. Zhang X, Wang J, Li J, Chen W, Liu C. 2018b. CRlncRC: A machine learning-based method for cancer-related long noncoding RNA identification using integrated features. BMC Med Genomics 11.

64. Zhang Y, Liao G, Bai J, Zhang X, Xu L, Deng C, Yan M, Xie A, Luo T, Long Z, et al. 2019b. Identifying Cancer Driver lncRNAs Bridged by Functional Effectors through Integrating Multi-omics Data in Human Cancers. Mol Ther - Nucleic Acids 17: 362–373. 10.1016/j.omtn.2019.05.030.

65. Zhao D, Zhang L, Zhang M, Xia B, Lv J, Gao X, Wang G, Meng Q, Yi Y, Zhu S, et al. 2020. Broad genic repression domains signify enhanced silencing of oncogenes. Nat Commun 11. 10.1038/s41467-020-18913-8.

66. Zhao T, Xu J, Liu L, Bai J, Xu C, Xiao Y, Li X, Zhang L. 2015. Identification of cancer-related lncRNAs through integrating genome, regulome and transcriptome features. Mol Biosyst 11: 126–136.

