## Supplementary Materials for "Cancer Driver Topologically Associated Domains identify oncogenic and tumor suppressive lncRNAs"

### Supplemental Materials for
### Cancer Driver Topologically Associated Domains identify oncogenic and tumor suppressive lncRNAs

Ziyan Rao *et al*

This PDF file includes:

- Supplemental Methods
- Supplemental Tables S1-11
- Supplemental Figs. S1-21

Other Supplemental Materials for this manuscript include the following:

Supplemental Data S1. The list of cancer lncRNA datasets CLC2, Lnc2Cancer, CGC genes, Pathway In Cancer (KEGG) genes, TUSON top 1000 oncogenes and tumor suppressors (in a separate Excel file).

Supplemental Data S2. The list of cancer driver PCGs in individual cancer types (prostate cancer, gastric cancer, and lung cancer) from TUSON (in a separate Excel file).

Supplemental Data S3. Detailed 139 Hi-C raw data information collected from ENCODE and GEO (in a separate Excel file).

Supplemental Data S4. The information about H3K4me3 and H3K27me3 peak files from ENCODE (in a separate Excel file).

Supplemental Data S5. The identified potential cancer driver lncRNAs through CADTAD pipeline in pan-cancer (in a separate Excel file).

Supplemental Data S6. The identified potential cancer driver lncRNAs through CADTAD pipeline in prostate cancer (in a separate Excel file).

Supplemental Data S7. The identified potential cancer driver lncRNAs through CADTAD pipeline in gastric cancer (in a separate Excel file).

Supplemental Data S8. The identified potential cancer driver lncRNAs through CADTAD pipeline in lung cancer (in a separate Excel file).

Supplemental Code. The code of CADTAD pipeline (in a separate zip file).

#### Supplemental methods

##### Bioinformatics analysis of Hi-C data.

Hi-C raw data were mapped to hg19 using bwa-mem2 (v.2.0) with the parameters E=50, L=0, and M (Ramírez et al. 2018), and then transferred to BAM files using SAMtools (Li et al. 2009) (v. 1.6). The downstream analysis was performed using HiCExplorer (Ramírez et al. 2018) (v. 3.7.2). Each enzyme's specific restriction sites were located using hicFindRestSite. To obtain a Hi-C matrix h5 file, we then utilized hicBuildMatrix with the following parameters: binSize = 1000, inputBufferSize = 200000, corresponding enzyme's restriction sites, and dangling sequence. Subsequently, we merged each h5 file under the parameters nb = 40 to create an h5 file with a bin size of 40kb, normalized them solely using hicNormalize with the parameter normalize = smallest, and corrected these files using hicCorrectMatrix with default parameters. Finally, we normalized them all using hicNormalize, with the parameter normalize = smallest.

We then utilized the normalized matrix to calculate the topologically associated domain (TAD) using hicFindTADs with the parameter correctForMultipleTesting = FDR. Domains.bed files were used for downstream TAD analysis. To distinguish the differences in TADs among normal and tumor samples, we converted h5 files into cool files using hicConvertFormat and chose the file with the largest TAD number as the basis for calculating the interval scores using chromosight (Matthey-Doret et al. 2020) (v.1.6.1). We analyzed these interval scores to plot the Principal Component Analysis (PCA) in R v.4.2.0 (R Core Team 2022). pyGenomeTracks was used to plot the TAD.

##### Random selection of non-cancer genes and non-cancer lncRNAs.

For the TAD ratio calculation (Fig. 1C-F), we chose typical TAD as the control; this type of TAD should overlap with at least one non-cancer PCG. We defined non-cancer PCGs as all PCGs excluding COSMIC genes, top1000 TUSON OGs/TSGs, and PATHWAY IN CANCER genes in KEGG. To compare binding (Fig. 2C-D), non-cancer PCGs and non-cancer lncRNAs were chosen at random. We randomly selected non-cancer PCGs from typical PCGs located in typical TADs with the same number as cancer driver PCGs. Similarly, after excluding CLC2 and Lnc2Cancer, non-cancer lncRNAs were randomly selected with the same number as cancer lncRNAs. For comparison of four lncRNA types (Fig. 2F), we considered all binding interactions between each lncRNA type and cancer driver PCGs.

##### Effectiveness comparison of cancer lncRNA discovery.

We obtained 30289 potential cancer lncRNAs filtered by the CDT feature in pan-cancer. We also acquired cancer lncRNAs by calculating the expression correlation and choosing different thresholds. To assess the cancer lncRNA discovery effectiveness using the CDT and co-expression characteristics, we performed enrichment analysis and calculated the ratio between these two types of identified cancer lncRNAs and cancer-related lncRNA datasets.

**Candidate lncRNAs and other overlapped PCGs knockdown by siRNAs**

LNCaP cells were transfected with siRNA pools targeting selected lncRNAs using RNAiMAX reagent. Cells were collected 48 hours after transfection for total RNA extraction and cell function assay. The knockdown efficiency was determined by RT-qPCR. The primers used for RT-qPCR were listed before. The sequence of each siRNA duplex for siRNA pools was listed in Supplemental Table S11.

#### References


Li H, Handsaker B, Wysoker A, Fennell T, Ruan J, Homer N, Marth G, Abecasis G, Durbin R. 2009. The Sequence Alignment/Map format and SAMtools. *Bioinformatics* **25**: 2078–2079.

Matthey-Doret C, Baudry L, Breuer A, Montagne R, Guiglielmoni N, Scolari V, Jean E, Campeas A, Chanut PH, Oriol E, et al. 2020. Computer vision for pattern detection in chromosome contact maps. *Nat Commun* **11**. http://dx.doi.org/10.1038/s41467-020-19562-7.

R Core Team. 2022. R: A Language and Environment for Statistical Computing. https://www.r-project.org.

Ramírez F, Bhardwaj V, Arrigoni L, Lam KC, Grüning BA, Villaveces J, Habermann B, Akhtar A, Manke T. 2018. High-resolution TADs reveal DNA sequences underlying genome organization in flies. *Nat Commun* **9**. http://dx.doi.org/10.1038/s41467-017-02525-w.

**Supplemental Table S1. The distance to TSS of cancer driver/random PCGs when cancer/random lncRNAs reaches a certain quantile.**

| Type | quantile | 0.1 | 0.2 | 0.3 | 0.4 | 0.5 | 0.6 | 0.7 | 0.8 | 0.9 |
|---|---|---|---|---|---|---|---|---|---|---|
| CLC2 | E (kb) | 80 | 240 | 520 | 840 | 1320 | 1800 | 2520 | 3760 | 6240 |
| | C1 (kb) | 440 | 880 | 1520 | 2360 | 3600 | 5280 | 7240 | 9680 | 13080 |
| | C2 (kb) | 160 | 440 | 680 | 960 | 1360 | 2080 | 2920 | 4400 | 6440 |
| Lnc2Cancer | E (kb) | 80 | 360 | 640 | 1000 | 1440 | 1840 | 2680 | 4120 | 7000 |
| | C1 (kb) | 520 | 1160 | 1880 | 2800 | 3760 | 5600 | 7740 | 10760 | 14600 |
| | C2 (kb) | 160 | 400 | 600 | 1040 | 1520 | 2120 | 3000 | 4280 | 6840 |

E: cancer lncRNAs & cancer driver PCGs
C1: cancer lncRNAs & random PCGs
C2: random lncRNAs & cancer driver PCGs

**Supplemental Table S2. Statistic of cancer/random lncRNA distance to cancer driver/random PCG.**

| Type | Group | mean | median | std |
|---|---|---|---|---|
| CLC2 | E (Mb) | 2.444 | 1.178 | 3.744 |
| | C1 (Mb) | 5.689 | 3.512 | 6.674 |
| | C2 (Mb) | 2.612 | 1.301 | 3.454 |
| Lnc2Cancer | E (Mb) | 2.606 | 1.304 | 3.724 |
| | C1 (Mb) | 6.153 | 3.697 | 6.850 |
| | C2 (Mb) | 2.697 | 1.439 | 3.600 |

E: cancer lncRNAs & cancer driver PCGs
C1: cancer lncRNAs & random PCGs
C2: random lncRNAs & cancer driver PCGs

**Supplemental Table S3. The p values of the comparison between cancer lncRNA ratios in CDTs and in other TADs, or between cancer lncRNA ratios and random lncRNA ratios in CDT by proportion test.**

| | LNCaP | | T47D | |
|---|---|---|---|---|
| | CLC2 | Lnc2Cancer | CLC2 | Lnc2Cancer |
| Up1 | $2.08\times10^{-5}$ | $2.52\times10^{-10}$ | $1.70\times10^{-4}$ | $1.19\times10^{-6}$ |
| Up2 | $7.39\times10^{-5}$ | $1.21\times10^{-11}$ | $5.06\times10^{-6}$ | $9.03\times10^{-10}$ |
| Up3 | $3.63\times10^{-3}$ | $2.13\times10^{-3}$ | $1.47\times10^{-5}$ | $4.57\times10^{-3}$ |
| Up4 | $2.24\times10^{-2}$ | $7.01\times10^{-5}$ | $2.09\times10^{-2}$ | $3.14\times10^{-4}$ |
| Up5 | $1.13\times10^{-3}$ | $1.69\times10^{-5}$ | $2.43\times10^{-4}$ | $2.94\times10^{-4}$ |
| Down1 | $4.59\times10^{-3}$ | $2.52\times10^{-8}$ | $2.37\times10^{-6}$ | $4.50\times10^{-7}$ |
| Down2 | $3.67\times10^{-4}$ | $2.70\times10^{-10}$ | $2.67\times10^{-4}$ | $4.69\times10^{-6}$ |
| Down3 | $7.80\times10^{-8}$ | $6.00\times10^{-12}$ | $2.48\times10^{-8}$ | $4.94\times10^{-8}$ |
| Down4 | $4.00\times10^{-4}$ | $2.47\times10^{-6}$ | $9.93\times10^{-7}$ | $1.28\times10^{-5}$ |
| Down5 | $1.06\times10^{-5}$ | $1.56\times10^{-12}$ | $4.83\times10^{-4}$ | $3.43\times10^{-7}$ |
| Typical TAD | $7.18\times10^{-5}$ | $4.31\times10^{-13}$ | $9.59\times10^{-8}$ | $1.59\times10^{-9}$ |
| Random lncRNAs | $4.69\times10^{-2}$ | $1.00\times10^{-2}$ | $6.48\times10^{-4}$ | $7.21\times10^{-2}$ |

**Supplemental Table S4. The comparison of the distances and expression correlations between I-CRL pairs and O-CRL pairs with similar distances.**

| | p (dist) | p (cor) | Mean (cor, I-CRL) | Mean (cor, O-CRL) | Number of remained pairs | Number of total pairs |
|---|---|---|---|---|---|---|
| CLC2 | 0.746 | 0.038 | 0.60 | 0.49 | 10 | 122 |
| Lnc2Cancer | 0.875 | 0.039 | 0.45 | 0.37 | 15 | 224 |

dist: distance
cor: the absolute value of expression correlation
I-CRL: cancer lncRNA in the CDT
O-CRL: cancer lncRNA outside the CDT
p (dist) was determined by two-sided paired *t*-test.
p (cor) was determined by one-sided paired *t*-test.

**Supplemental Table S5. GLM results about lncRNAs in CLC2.**

| Model | Variable | Coefficient | Std.error | z | p-value |
|---|---|---|---|---|---|
| Raw model $R^2$= 0.6338 | Intercept | 2.9610 | 0.029 | 102.391 | <0.001 |
| | CDT | 2.4471 | 0.057 | 43.239 | <0.001 |
| | Distance (Mb) | -0.0039 | <0.001 | -8.804 | <0.001 |
| Expression correlations (shuffled) $R^2$ = 0.0012 | Intercept | 2.8742 | 0.043 | 67.619 | <0.001 |
| | CDT | -0.3875 | 0.329 | -1.178 | 0.239 |
| | Distance (Mb) | 0.0003 | 0.001 | 0.589 | 0.556 |
| CDT (shuffled) $R^2$ = 0.1033 | Intercept | 3.2453 | 0.036 | 90.973 | <0.001 |
| | CDT | -0.2256 | 0.262 | -0.860 | 0.390 |
| | Distance (Mb) | -0.0072 | 0.001 | -11.747 | <0.001 |
| Distance (shuffled) $R^2$ = 0.6117 | Intercept | 2.7494 | 0.029 | 94.319 | <0.001 |
| | CDT | 2.6515 | 0.054 | 49.327 | <0.001 |
| | Distance (Mb) | 0.0001 | <0.001 | 0.302 | 0.753 |

**Supplemental Table S6. GLM results about lncRNAs in Lnc2Cancer.**

| Model | Variable | Coefficient | Std.error | z | p-value |
|---|---|---|---|---|---|
| Raw model $R^2$= 0.4718 | Intercept | 2.8870 | 0.021 | 137.425 | <0.001 |
| | CDT | 2.3547 | 0.043 | 54.329 | <0.001 |
| | Distance (Mb) | -0.0033 | <0.001 | -10.846 | <0.001 |
| Expression correlations (shuffled) $R^2$ = 0.0003 | Intercept | 2.8276 | 0.030 | 94.511 | <0.001 |
| | CDT | -0.0694 | 0.198 | -0.351 | 0.726 |
| | Distance (Mb) | -0.0002 | <0.001 | -0.585 | 0.558 |
| CDT (shuffled) $R^2$ = 0.0634 | Intercept | 3.1310 | 0.026 | 122.397 | <0.001 |
| | CDT | 0.0779 | 0.167 | 0.465 | 0.642 |
| | Distance (Mb) | -0.0060 | <0.001 | -14.848 | <0.001 |
| Distance (shuffled) $R^2$ = 0.4510 | Intercept | 2.7106 | 0.021 | 128.213 | <0.001 |
| | CDT | 2.5384 | 0.041 | 61.703 | <0.001 |
| | Distance (Mb) | -0.0001 | <0.001 | -0.509 | 0.611 |

**Supplemental Table S7. The sequences of each guide RNAs for candidate lncRNAs.**

| Gene | pgRNA | gRNA1 sequence | gRNA2 sequence |
|---|---|---|---|
| NT | | CTGAAAAAGGAAGGAGTTGA | AAGATGAAAGGAAAGGCGTT |
| HSALNG0 082024 | p1 | AGGCTCCCTCTGAGCCGACA | GTGTGGCCACCGGAACACCA |
| | p2 | AGGCTCCCTCTGAGCCGACA | CGTCCCCCTCCTAGTCTCCG |
| HSALNG0 069873 | p1 | GCTCTGCTAGATAAATCTGG | AGGGCGCCGCTTAAAGTGTG |
| | p2 | GCTCTGCTAGATAAATCTGG | CAGGGTGTGGAGCCGCACTG |
| HSALNG0 081989 | p1 | GTCAGGACACAGATGACCGA | ACCTTGGCAAGTGAAGACGC |
| | p2 | GTCAGGACACAGATGACCGA | AAACATTCAGGCCCCCACCG |
| HSALNG0 125569 | p1 | GCTGTGTCAACCATGCCCCG | TTCCAACCAGAATAGGACCA |
| | p2 | GCTGTGTCAACCATGCCCCG | CAGAGCTCGTAATTGCTGCG |
| HSALNG0 094538 | p1 | GCTCTTCTGATACTGACCCG | TAGTAGGGCTCAAATCTCTG |
| | p2 | CTAAGATGTCCAAGTCCTAG | TAGTAGGGCTCAAATCTCTG |
| HSALNG0 133581 | p1 | GTCCACAGCAAGAGTGTCCG | CTTCCCTCCAAAAATCACCG |
| | p2 | TGTTCCTGGGTGGCGACGTG | AGGTTTCTAGACGTGACCCA |
| HSALNG0 116774 | p1 | GTTGCAGCCATCGTCCTGGG | TTTCGGCATGTGATCCGAGG |
| | p2 | GTTGCAGCCATCGTCCTGGG | GCGGGACTTCCAGAGCCCGA |
| HSALNG0 058614 | p1 | GGTGTGATAACTTTACTGAT | CTCGCCCGCTTCGCCCGGGT |
| | p2 | ATGTTATCAGCCGCACACGG | CTCGCCCGCTTCGCCCGGGT |
| HSALNG0 124011 | p1 | CACTTGGAAACAATTACTGT | AAGTCGCCGACAACTGCCTG |
| | p2 | TTTCTGATATTAATCAGGCG | AAGTCGCCGACAACTGCCTG |
| HSALNG0 117120 | p1 | GCTGAGGCTATTGTCCGGGG | AGGACACTGTTAGACCACCT |
| | p2 | TCACCCATCGACTACCCGGG | GTGAAACATTAACCCCCTCA |

**Supplemental Table S8. The primers used for PCR and RT-qPCR of candidate lncRNAs.**

| Gene | PCR primers | Real-time PCR primers |
|---|---|---|
| HSALNG 0082024 | F: TGAGACTACCTGCAGGACCA | F: CCTTCCTCAGCCATTGTCCA |
| | R: GGAAGTGACACTCCGTCCTC | R: TTGCTTCACAGTGCCCCTAA |
| HSALNG 0069873 | F: GTTCTGTCCGGCCATTCTGT | F: TGGGCCTTTGGTCAGTCTTT |
| | R: ATCATCTTCCATCTCCGCCG | R: TGCTATGTTTGGGGGTTGGG |
| HSALNG 0081989 | F: GCAGACAGAGGGGTCAACAG | F: TGGGGGCCATTCTCAATGTT |
| | R: CTCAGGTGCCTCTGGAACTC | R: AGAGAGCAACGCTCTGTGAG |
| HSALNG 0125569 | F: CCAGGGTGGAACCCAACTATT | F: CCAGCCATGTGGTGTAGACG |
| | R: ACATCCTGATGATGCCTCCA | R: AGATGCAATCGCAGATGTGG |
| HSALNG 0094538 | F: TGCGGGAGGAGAGACTACAT | F: GGCCCAGCGATGCTTCTTAT |
| | R: GGTTAGCTTCAGAGCTGCCA | R: TAGCTTCAGAGCTGCCAAGG |
| HSALNG 0133581 | F: GGGGACACCACTCCTCCATC | F: GAGAATGGAGCGGAGGAACC |
| | R: TGGCTCAGGAATTTGCTCCC | R: TGTGTCATGGCTGACATGCT |
| HSALNG 0116774 | F: CGGCTCTGACCTGAGCTATG | F: TGAGAGGCTACGCTGTATGG |
| | R: CCCCCTGCACTCTGAATAGG | R: GAGACATGGTACAGGGTGGG |
| HSALNG 0058614 | F: AGCCATGTGAAACAACACACA | F: ATCGGTTGATCCAAAAGCAG |
| | R: AGTTTCTGAAGCGTCCCTCG | R: ACCTCAGCCTCCCTGGTAGT |
| HSALNG 0124011 | F: CCCTACAAACCTAACCCCAACA | F: AGAGGCACGTGGTTCGAAAT |
| | R: CATTTTGGCTTCAGGGGCTG | R: AGTCATTGGCATGGAGTGGG |
| HSALNG 0117120 | F: AGCCTGGGAAGTTCGCTTAC | F: CAGCCCGGTTTGGGAGTAAA |
| | R: AAAGATTGAGGGTCGGGCTG | R: AGCCAGACAGAAACTACGCC |

**Supplemental Table S9. The primers used for RT-qPCR of associated driver PCGs.**

| Gene | Driver PCG | PCR primers |
|---|---|---|
| HSALNG0082024 | *RASSF7* | F: CTCTGAGTCCCATGCTGGTG |
| | | R: GTCTTGCAGCCCTCACTAGG |
| HSALNG0069873 | *GLDC* | F: CAGAAGGTCTCAAGCGAGCA |
| | | R: GTGCCATCCTCAAAAAGCCG |
| HSALNG0081989 | *RASSF7* | F: CTCTGAGTCCCATGCTGGTG |
| | | R: GTCTTGCAGCCCTCACTAGG |
| HSALNG0125569 | *HPN* | F: GACCCCAGGCCCAATCTG |
| | | R: CCATCCAGGCAGGCTGTAG |
| HSALNG0094538 | *MLXIP* | F: AGAGCCCCAGTCCTCAATCT |
| | | R: TGCTTCATCTGCCGGTTCTT |
| HSALNG0133581 | *ITGB2* | F: ACTGGTAGCAAAGCCCCCAC |
| | | R: GCACTCCTGAGAGAGGACGC |
| HSALNG0116774 | *KIF18B* | F: CGACGAAAGGTGTCACCACA |
| | | R: AGGGTTAAACACCAGCACCC |
| HSALNG0058614 | *KCTD7* | F: CACATGGTGGTGAGTAGCCC |
| | | R: GGATGCTGTTCTCACCTCGG |
| HSALNG0124011 | *KANK2* | F: CCCAGAAGTTCCTGCCGAAT |
| | | R: GTGCTCTTCTCAGGCTGTGT |
| HSALNG0117120 | *ZNF652* | F: GAGACACCGCAGAACTCACA |
| | | R: TCACCCGAAAGCACTTCTCC |

**Supplemental Table S10. The primers used for RT-qPCR of overlapped PCGs.**

| Overlapped PCG | PCR primers |
|---|---|
| *GATD1* | F: GTGACTGAGAGCAATGCACG |
| | R: TGGAAGTGCTGCAGGATACG |
| *MLEC* | F: CCCGAGAGCGTCATTTGGG |
| | R: AACGCAGGATTGGCAGTTTCA |
| *ADAM11* | F: GAAGGAAAAGGCAGGTCCGC |
| | R: AGGACGATGCGAGTGTTGAG |
| *RABGEF1* | F: TGTGGTTACTACGGCAACCC |
| | R: TCCCGCTGGAGTCGCT |

**Supplemental Table S11. The sequence of each siRNA duplex for siRNA pools.**

| Gene | siRNA duplex | |
|---|---|---|
| | sense (5'-3') | antisense (5'-3') |
| HSALNG 0094538 | GCAGCAACAGAACACUUGATT | UCAAGUGUUCUGUUGCUGCTT |
| | GGCUGCAGUCAGAGGUAUATT | UAUACCUCUGACUGCAGCCTT |
| | GCAGUGGACAUCCUAACUATT | UAGUUAGGAUGUCCACUGCTT |
| | GGCCUACUCUGGCAUAAUUTT | AAUUAUGCCAGAGUAGGCCTT |
| HSALNG 0125569 | GGACAGAGUCUUGUACCUATT | UAGGUACAAGACUCUGUCCTT |
| | GCACCCAGCCAAUAAUAAUTT | AUUAUUAUUGGCUGGGUGCTT |
| | GCCAUCCGCACAAUUGAAUTT | AUUCAAUUGUGCGGAUGGCTT |
| | GGCUAUGCACUCAAUAAGUTT | ACUUAUUGAGUGCAUAGCCTT |
| HSALNG 0069873 | GGCGGAGAUGGAAGAUGAUTT | AUCAUCUUCCAUCUCCGCCTT |
| | GGUUUCUGCCUGGUAUUGUTT | ACAAUACCAGGCAGAAACCTT |
| | GCUGCCCAAUUCAAGAAAUTT | AUUUCUUGAAUUGGGCAGCTT |
| | GGGACACCAAGAAUAAUCUTT | AGAUUAUUCUUGGUGUCCCTT |
| *GATD1* | GAUGUGACUGAGAGCAAUGTT | CAUUGCUCUCAGUCACAUCTT |
| | GUGUGUUCAUCUACCCAUGTT | CAUGGGUAGAUGAACACACTT |
| | GUGGAUGUGACUGAGAGCATT | UGCUCUCAGUCACAUCCACTT |
| *MLEC* | GUACGGUCUGAAGUUAUGUTT | ACAUAACUUCAGACCGUACTT |
| | GACUACGUGCUGGUCUUGATT | UCAAGACCAGCACGUAGUCTT |
| | GAGCGGUACAAUGAGGAGATT | UCUCCUCAUUGUACCGCUCTT |
| *ADAM11* | CAUCGAUUAAAGAUCACAUTT | AUGUGAUCUUUAAUCGAUGTT |
| | GACGAGUACAACCAGUUUCTT | GAAACUGGUUGUACUCGUCTT |
| | GGACGGUUACUACUGUGACTT | GUCACAGUAGUAACCGUCCTT |
| *RABGEF1* | GUUACGCCUCAGAUGCUGUTT | ACAGCAUCUGAGGCGUAACTT |
| | CUCCGUUAAGAACCACUGATT | UCAGUGGUUCUUAACGGAGTT |
| | GGUCGGAUCAAAGAAGGAATT | UUCCUUCUUUGAUCCGACCTT |

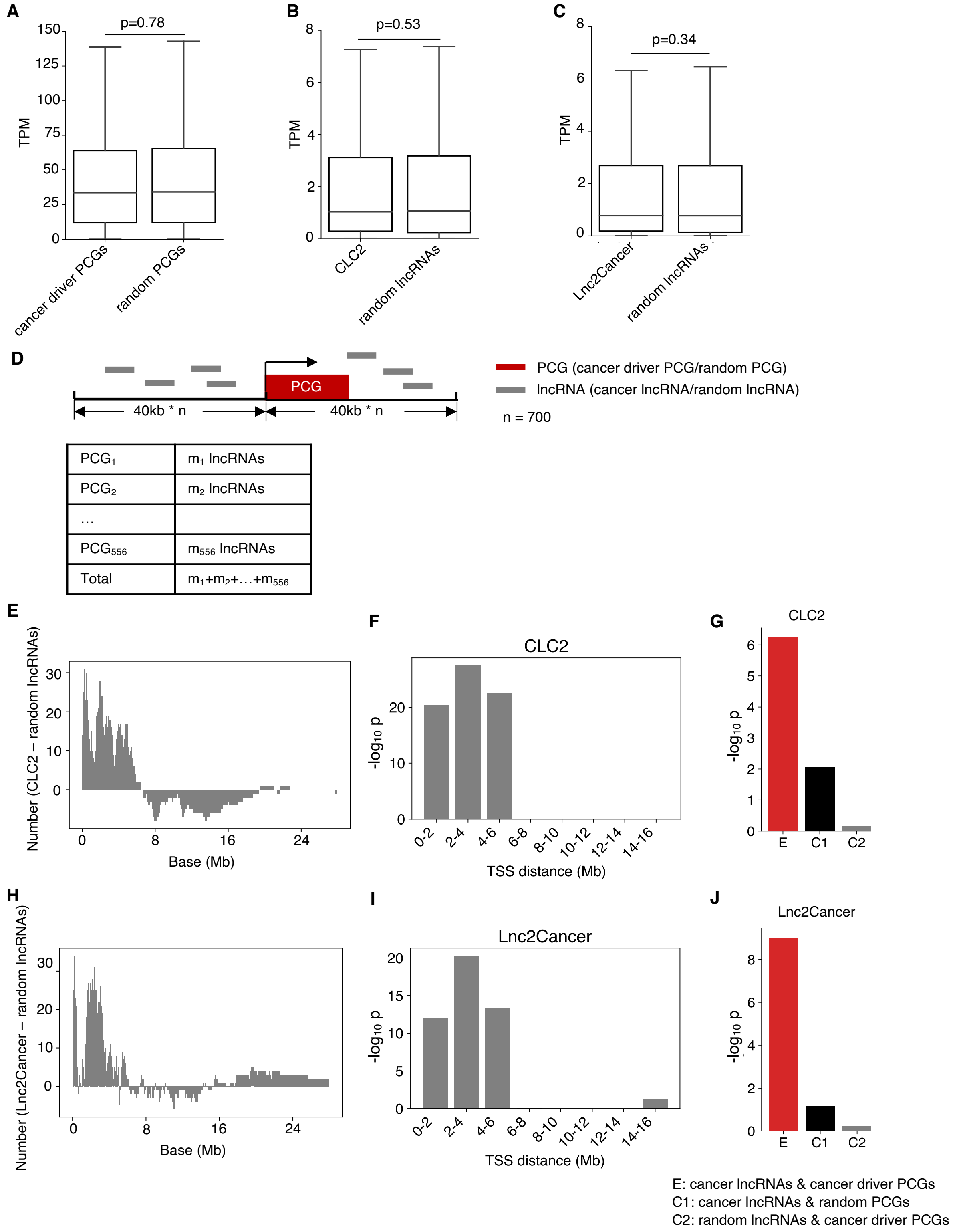


**Supplemental Fig. S1. Randomly select no-cancer PCGs and lncRNAs, and compare cancer/random lncRNA distribution near the cancer driver PCG.**

A. Boxplot to show the expression distribution of cancer driver PCGs and randomly selected PCGs with the same number. The P value is determined by two tail *t*-test.

B. Boxplot to show the expression distribution of cancer lncRNAs in CLC2 and randomly selected lncRNAs with the same number. The P value is determined by two tail *t*-test.

C. Boxplot to show the expression distribution of cancer lncRNAs in Lnc2Cancer and randomly selected lncRNAs with the same number. The P value is determined by two tail *t*-test.

D. Model of a statistic of nearby lncRNAs around cancer driver PCGs/random PCGs.

E. The quantity difference between cancer lncRNAs in CLC2 and random lncRNAs in a series of extended regions around the TSS of cancer driver PCGs.

F. The difference between the numbers of cancer lncRNAs in CLC2 and the numbers of random lncRNAs across a series of bins in large intervals. P values are determined by one tail paired *t*-test.

G. The enrichment of the closest lncRNAs near cancer driver/random PCGs in cancer/random lncRNAs (CLC2). The P values are determined by one tail Fisher's exact test.

H. The quantity difference between cancer lncRNAs in Lnc2Cancer and random lncRNAs in a series of extended regions around the TSS of cancer driver PCGs.

I. The difference between the numbers of cancer lncRNAs in Lnc2Cancer and the numbers of random lncRNAs across a series of bins in large intervals. P values are determined by one tail paired *t*-test.

J. The enrichment of the closest lncRNAs near cancer driver/random PCGs in cancer/random lncRNAs (Lnc2Cancer). The P values are determined by one tail Fisher's exact test.

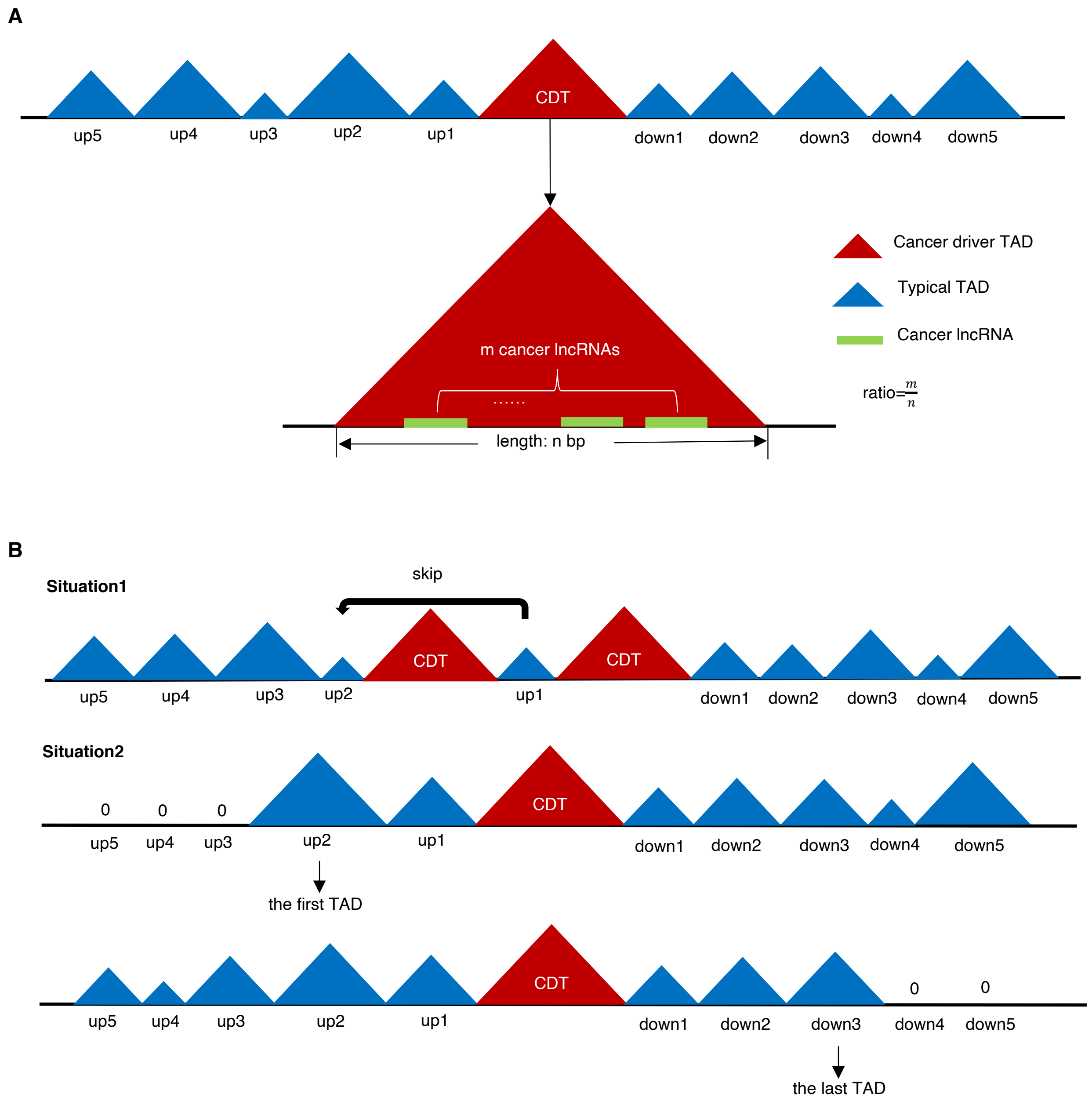


**Supplemental Fig. S2. The models for calculating the cancer lncRNA density in a TAD.**

A. Schematic illustrating cancer driver TAD (CDT) and the calculation of cancer lncRNA density.

B. Models of different situations to find up-stream and down-stream typical TADs.

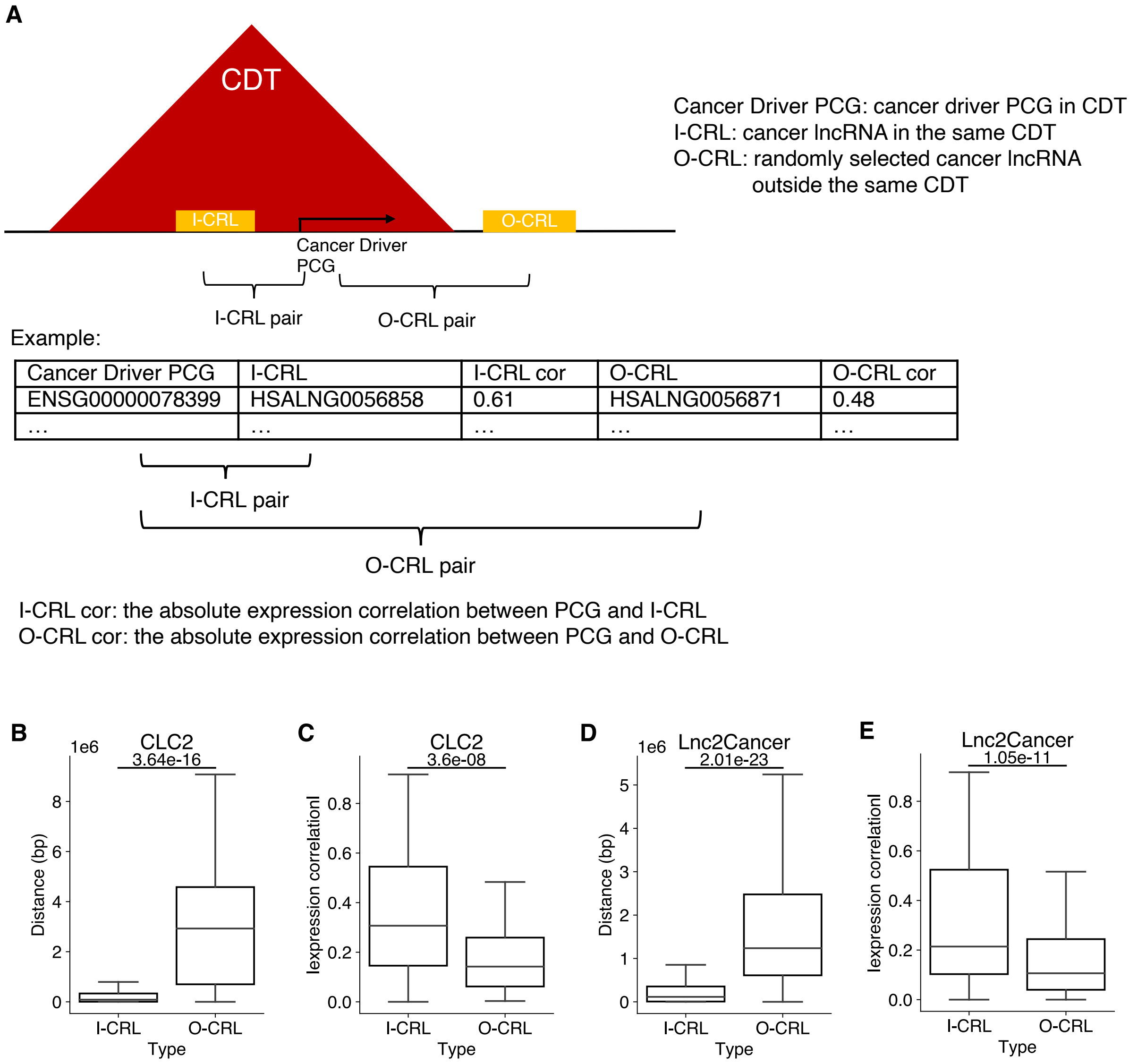


| Cancer Driver PCG | I-CRL | I-CRL cor | O-CRL | O-CRL cor |
|---|---|---|---|---|
| ENSG00000078399 | HSALNG0056858 | 0.61 | HSALNG0056871 | 0.48 |
| … | … | … | … | … |

**Supplemental Fig. S3. The schematic diagram of O-CRL selection and comparison of co-expression levels between I-CRL pairs and O-CRL pairs under distance control.**

(A) The schematic diagram of O-CRL selection for the comparison between I-CRL pairs and O-CRL pairs. Boxplot to show the distance between I-CRL pairs and O-CRL pairs in CLC2 (B) and Lnc2Cancer (D). P value is determined by one-tailed paired *t*-test.

Boxplot to show the difference in absolute Spearman's correlation between I-CRL pairs and O-CRL pairs in CLC2 (C) and Lnc2Cancer (E). P value is determined by one-tailed paired *t*-test.

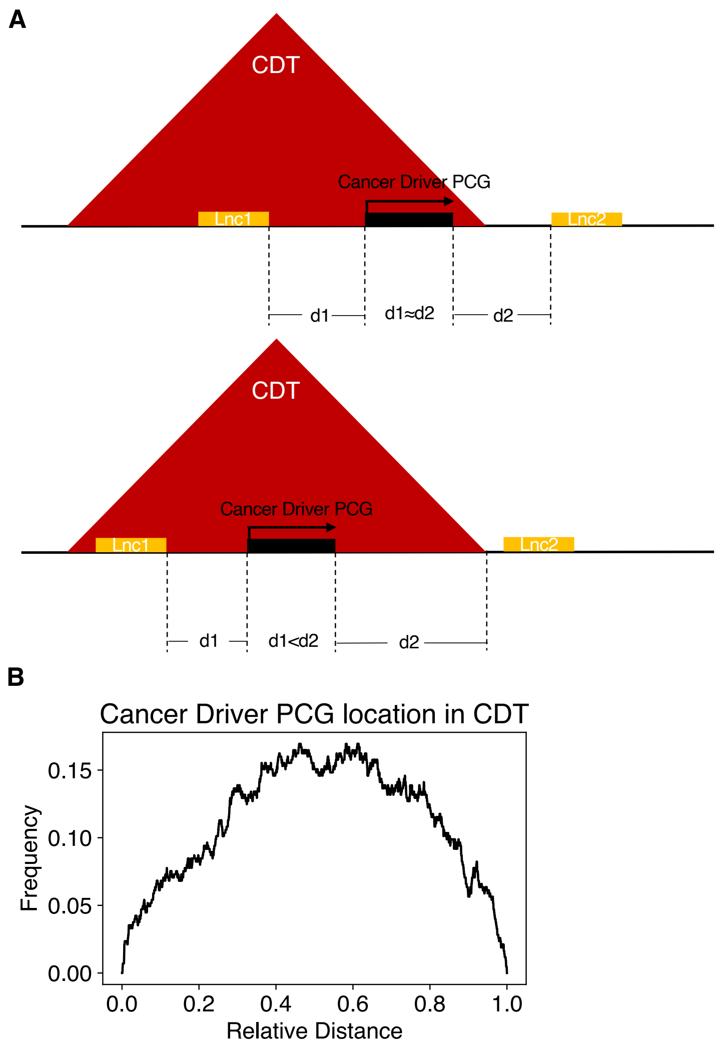


**Supplemental Fig. S4. The distribution of cancer driver PCGs location in CDTs.**

A. Schematic illustrating two types of cancer driver PCG location distribution in a CDT.

B. Frequency diagram of cancer driver PCGs relative position distribution in CDTs.

CDT: cancer driver TAD; Lnc: lncRNA gene.

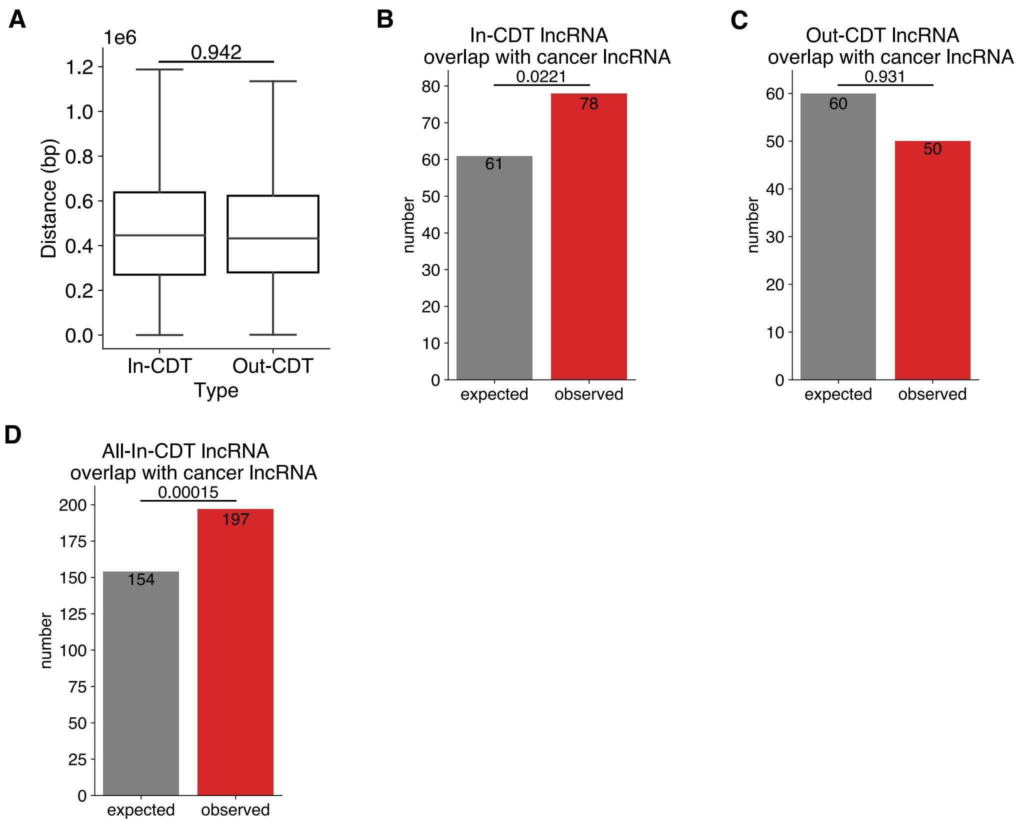


**Supplemental Fig. S5. Cancer lncRNA enrichment of lncRNAs within and outside CDTs.**

A. Boxplot to show the distance between cancer driver PCGs and selected lncRNAs within/outside CDTs. P value is determined by two-sided paired *t*-test.

B-D. Bar plot to show the enrichment of overlap between cancer lncRNAs and selected lncRNAs within CDTs (B)/selected lncRNAs outside CDTs (C)/all lncRNAs within CDTs (D). P values are determined by one tail Fisher's exact test.

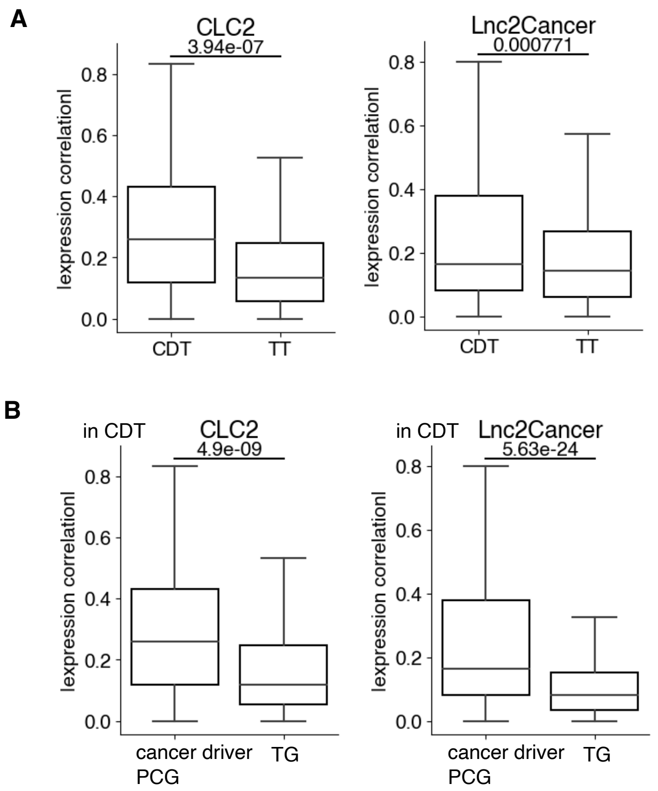


**Supplemental Fig. S6. Comparison between CDTs and Typical TADs, or cancer driver PCGs and non-cancer PCGs.**

A. Boxplot to show the difference of absolute Spearman's correlation between CDTs and TTs about CLC2 (left) and Lnc2Cancer (right). P value is determined by one-tailed paired *t*-test. TT: Typical TAD.

B. Boxplot to show the difference of absolute Spearman's correlation between cancer driver PCGs and non-cancer PCGs in CDT about CLC2 (left) and Lnc2Cancer (right). P value is determined by one-tailed paired *t*-test. TG: Typical genes (non-cancer PCGs).

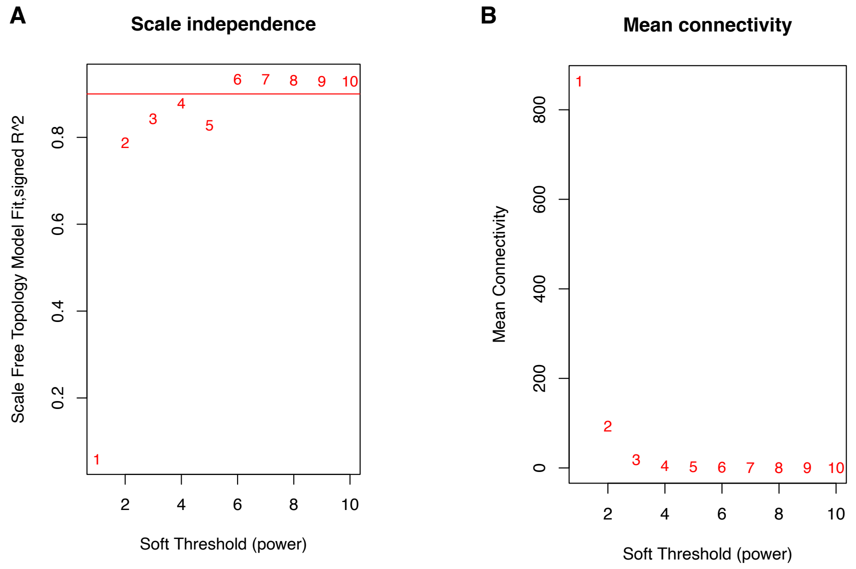


**Supplemental Fig. S7. The characteristics of constructing a lncRNA-mRNA co-expression network.**

A. Network properties plot to visualize the scale-free topology fitting index. Points are labeled by the corresponding power parameters. The red line indicates 0.9.

B. Network properties plot to visualize the mean connectivity. Points are labeled by the corresponding power parameters.

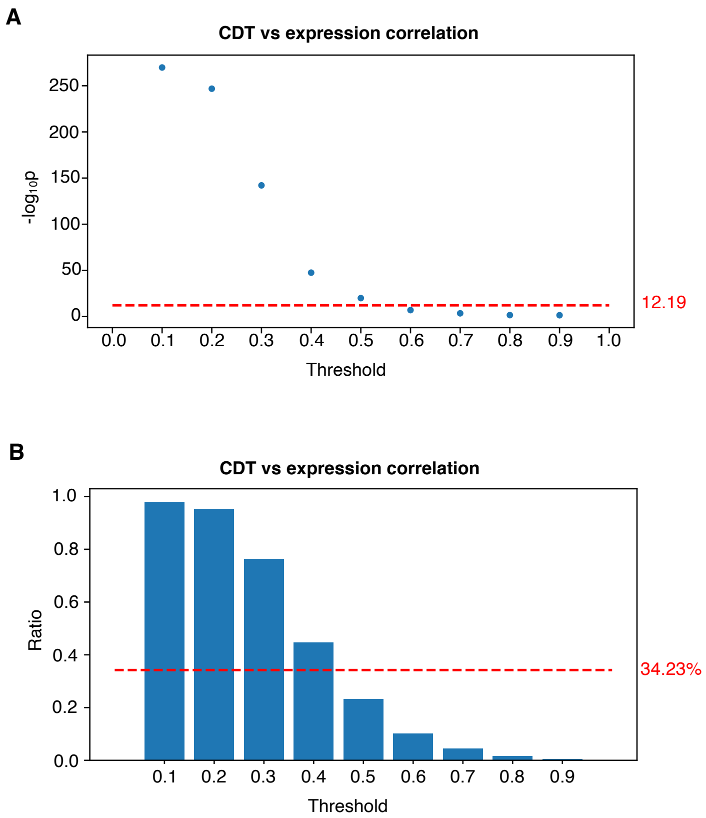


**Supplemental Fig. S8. The effectiveness comparison of identifying cancer lncRNAs using CDT or expression correlation features.**

A. Blue dots to show the enrichment degree in cancer lncRNA datasets of different thresholds of lncRNAs filtered by expression correlation features. Red line to show the enrichment degree of lncRNAs filtered by CDT. P value is determined by one tail Fisher's exact test.

B. Blue bars to show the ratio of cancer lncRNAs filtered under different expression correlation thresholds to all original cancer lncRNAs. Red line to show the ratio of cancer lncRNAs filtered by CDT to all original cancer lncRNAs.

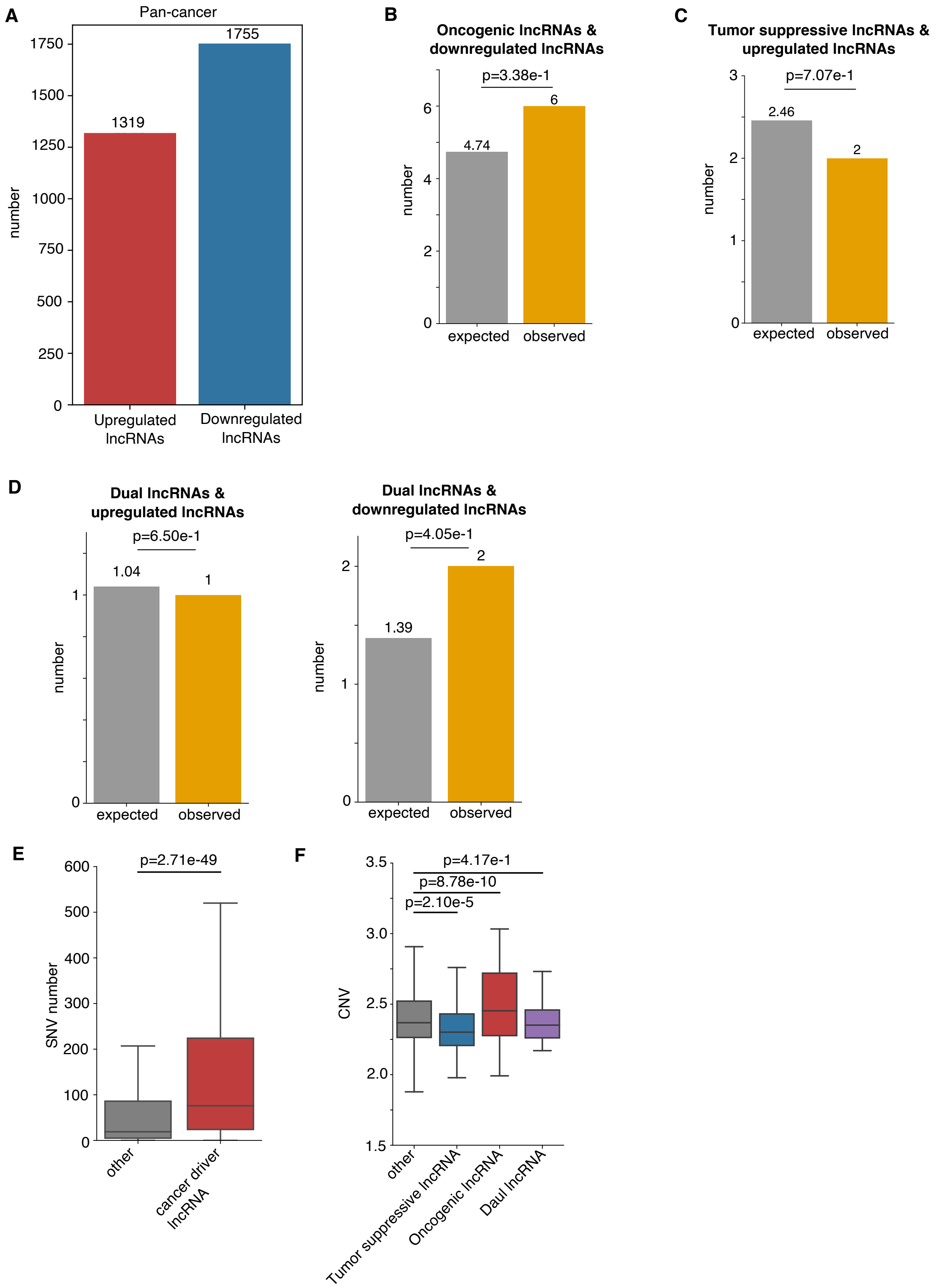
A
Pan-cancer
1319
1755
number
Upregulated lncRNAs
Downregulated lncRNAs
B
Oncogenic lncRNAs & downregulated lncRNAs
p=3.38e-1
4.74
6
expected
observed
C
Tumor suppressive lncRNAs & upregulated lncRNAs
p=7.07e-1
2.46
2
expected
observed
D
Dual lncRNAs & upregulated lncRNAs
p=6.50e-1
1.04
1
Dual lncRNAs & downregulated lncRNAs
p=4.05e-1
1.39
2
E
p=2.71e-49
SNV number
other
cancer driver lncRNA
F
p=4.17e-1
p=8.78e-10
p=2.10e-5
CNV
other
Tumor suppressive lncRNA
Oncogenic lncRNA
Daul lncRNA

**Supplemental Fig. S9. Cancer-related features of putative pan-cancer driver lncRNAs.**

A. Bar plot to show the number of upregulated and downregulated lncRNAs in cancer compared to normal samples.

B. Bar plot to show the enrichment of oncogenic lncRNA candidates in downregulated lncRNAs in cancer. P value is determined by one tail Fisher's exact test.

C. Bar plot to show the enrichment of tumor suppressive lncRNA candidates in upregulated lncRNAs in cancer. P value is determined by one tail Fisher's exact test.

D. Bar plot to show the enrichment of putative dual-function lncRNAs in upregulated lncRNAs and downregulated lncRNAs in cancer. P values are determined by one tail Fisher's exact test.

E. Boxplot to show the SNV number of putative cancer driver lncRNAs in pan-cancer and the remained lncRNAs. The P value is determined by one tail Wilcoxon test.

F. Boxplot to show the CNV occupation of predicted tumor suppressive lncRNAs, oncogenic lncRNAs, dual-function lncRNAs, and the remained lncRNAs. The P values are determined by two tail Wilcoxon test.

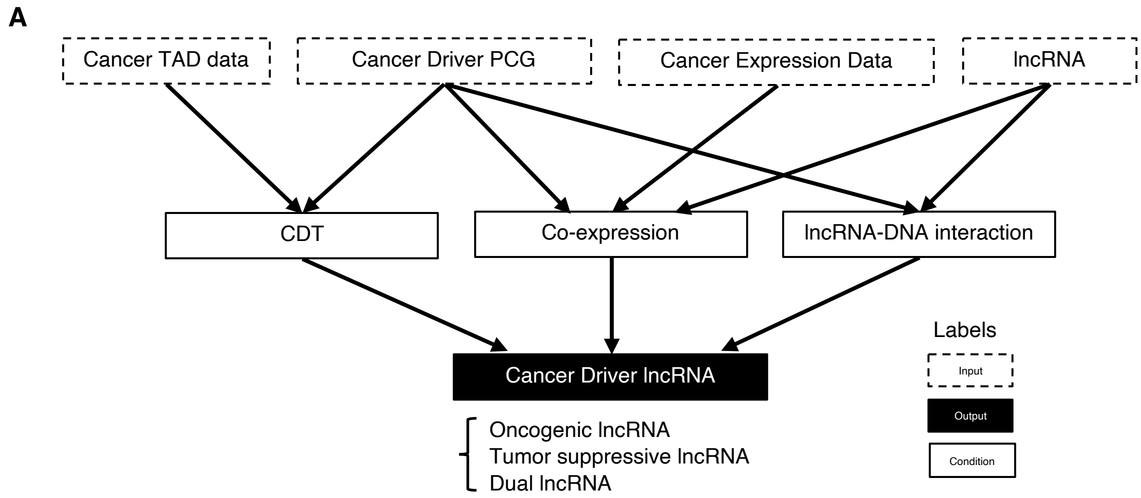


**Supplemental Fig. S10. The diagram of CADTAD.**

A. The flowchart of CADTAD about input, output, and conditions.

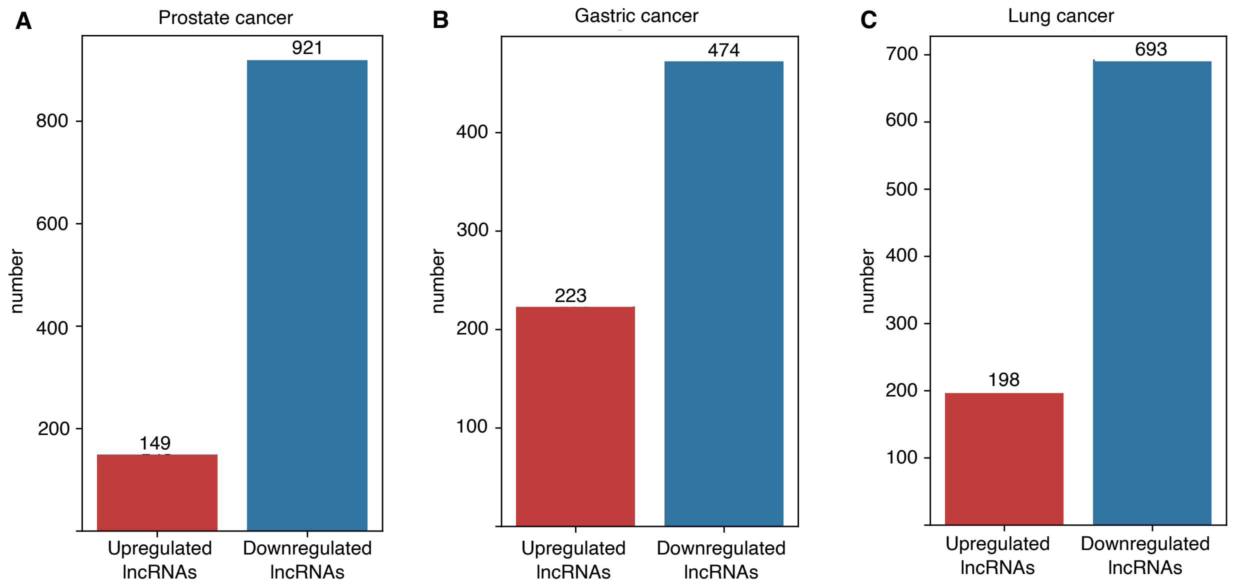


**Supplemental Fig. S11. The statistics of differentially expressed gene numbers in individual cancer.** Bar plot to show the number of upregulated and downregulated lncRNAs in cancer samples compared to the normal samples about prostate cancer (A), gastric cancer (B), and lung cancer (C).

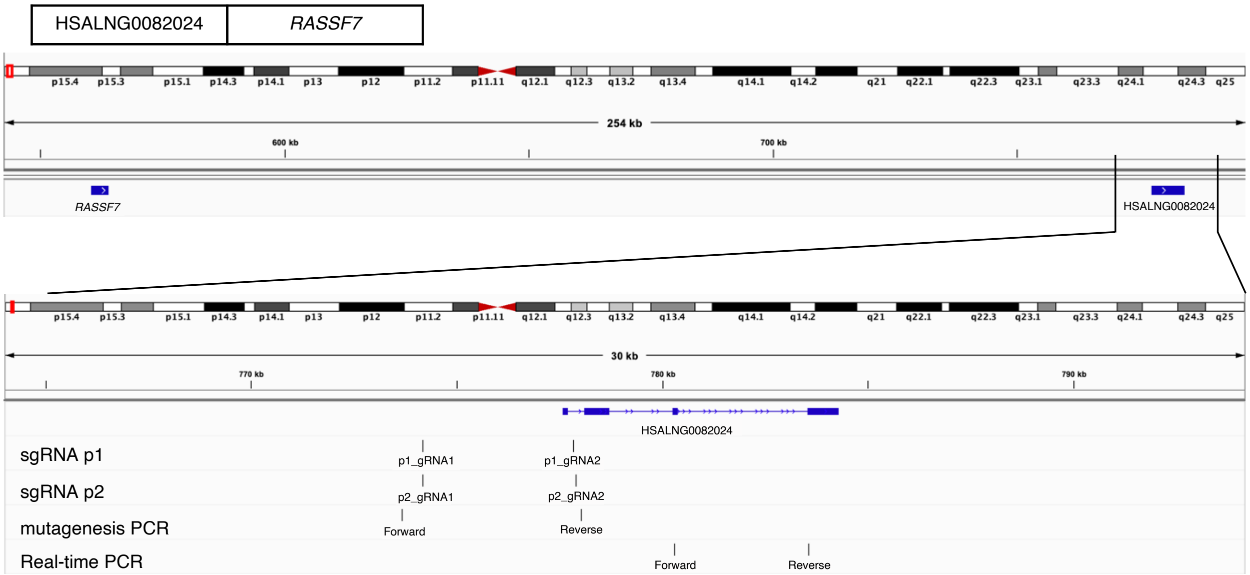
HSALNG0082024
RASSF7
254 kb
600 kb
700 kb
RASSF7
HSALNG0082024
30 kb
770 kb
780 kb
790 kb
HSALNG0082024
sgRNA p1
p1_gRNA1
p1_gRNA2
sgRNA p2
p2_gRNA1
p2_gRNA2
mutagenesis PCR
Forward
Reverse
Real-time PCR
Forward
Reverse

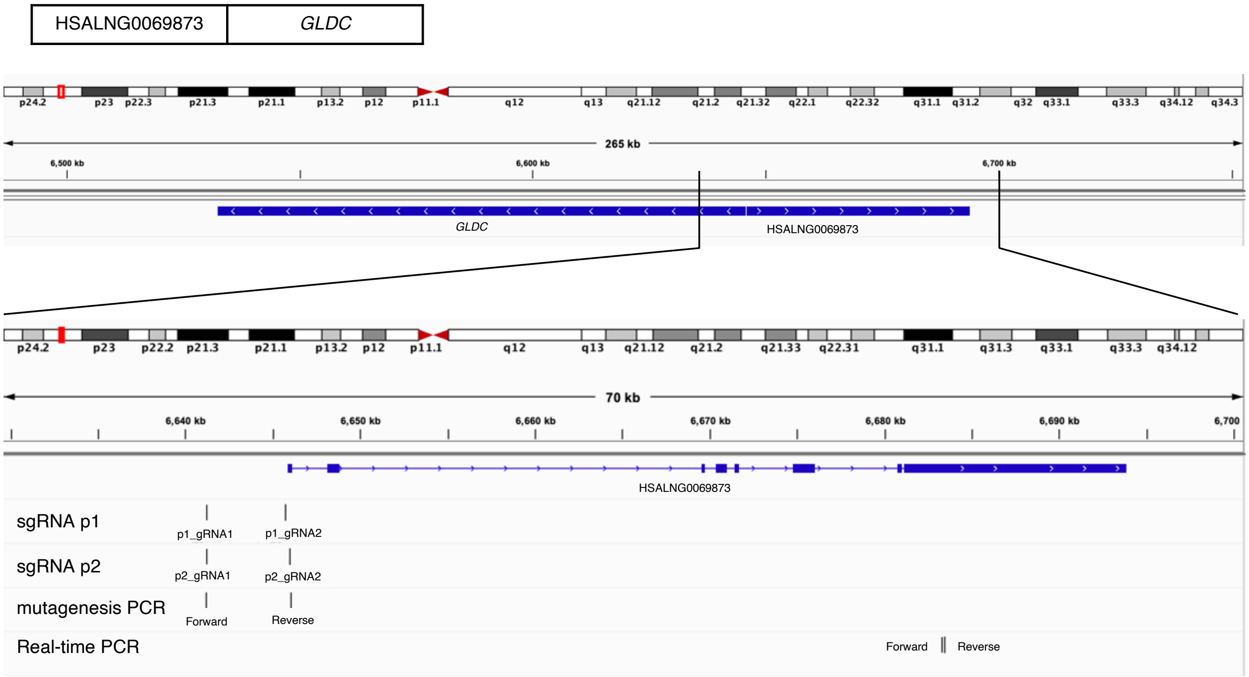
HSALNG0069873
GLDC
265 kb
6,500 kb
6,600 kb
6,700 kb
GLDC
HSALNG0069873
70 kb
6,640 kb
6,650 kb
6,660 kb
6,670 kb
6,680 kb
6,690 kb
6,700
HSALNG0069873
sgRNA p1
p1_gRNA1
p1_gRNA2
sgRNA p2
p2_gRNA1
p2_gRNA2
mutagenesis PCR
Forward
Reverse
Real-time PCR
Forward
Reverse

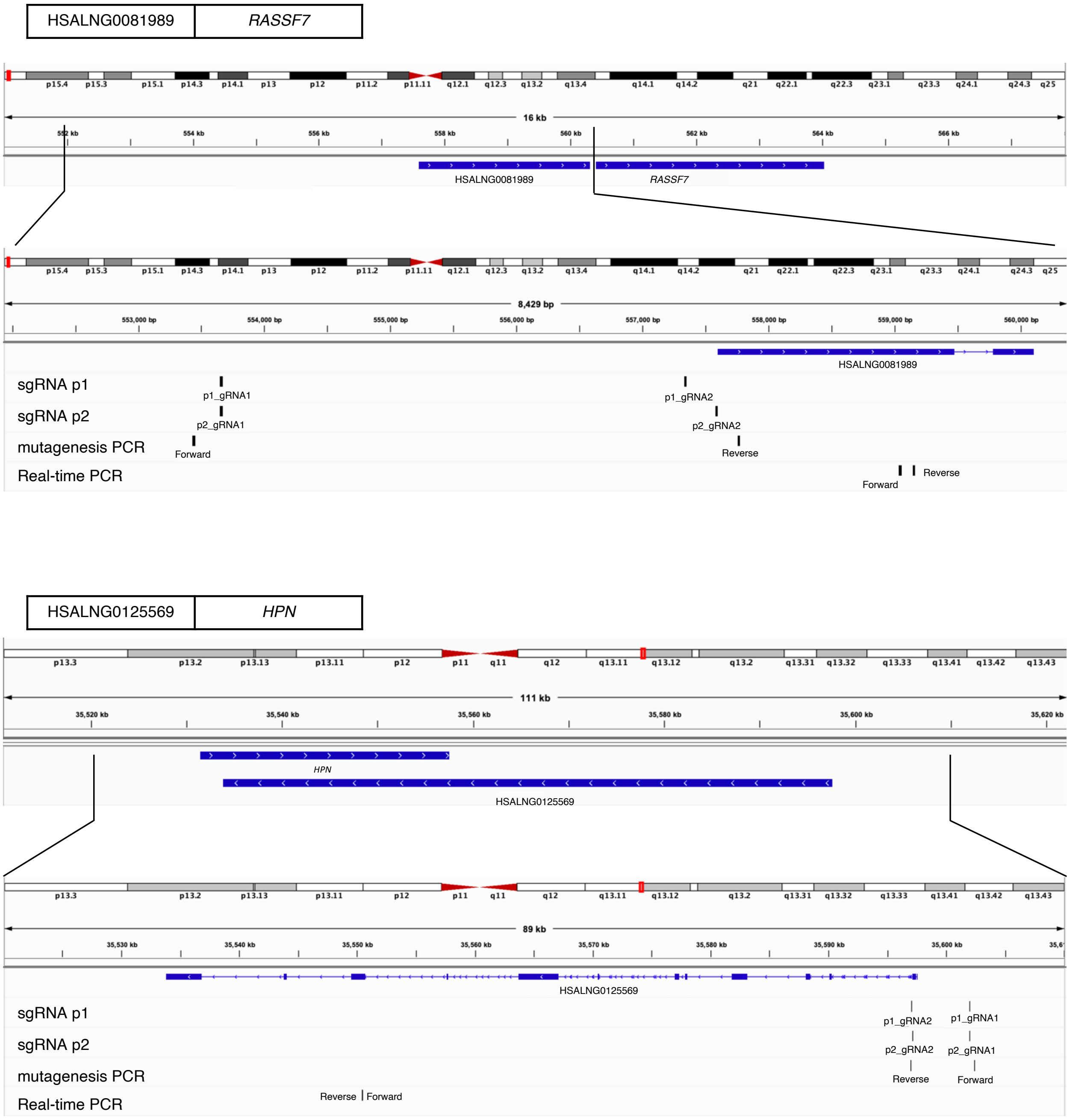
HSALNG0081989
RASSF7
16 kb
HSALNG0081989
RASSF7
8,429 bp
HSALNG0081989
sgRNA p1
p1_gRNA1
p1_gRNA2
sgRNA p2
p2_gRNA1
p2_gRNA2
mutagenesis PCR
Forward
Reverse
Real-time PCR
Forward
Reverse
HSALNG0125569
HPN
111 kb
HPN
HSALNG0125569
89 kb
HSALNG0125569
sgRNA p1
p1_gRNA2
p1_gRNA1
sgRNA p2
p2_gRNA2
p2_gRNA1
mutagenesis PCR
Reverse
Forward
Real-time PCR
Reverse
Forward

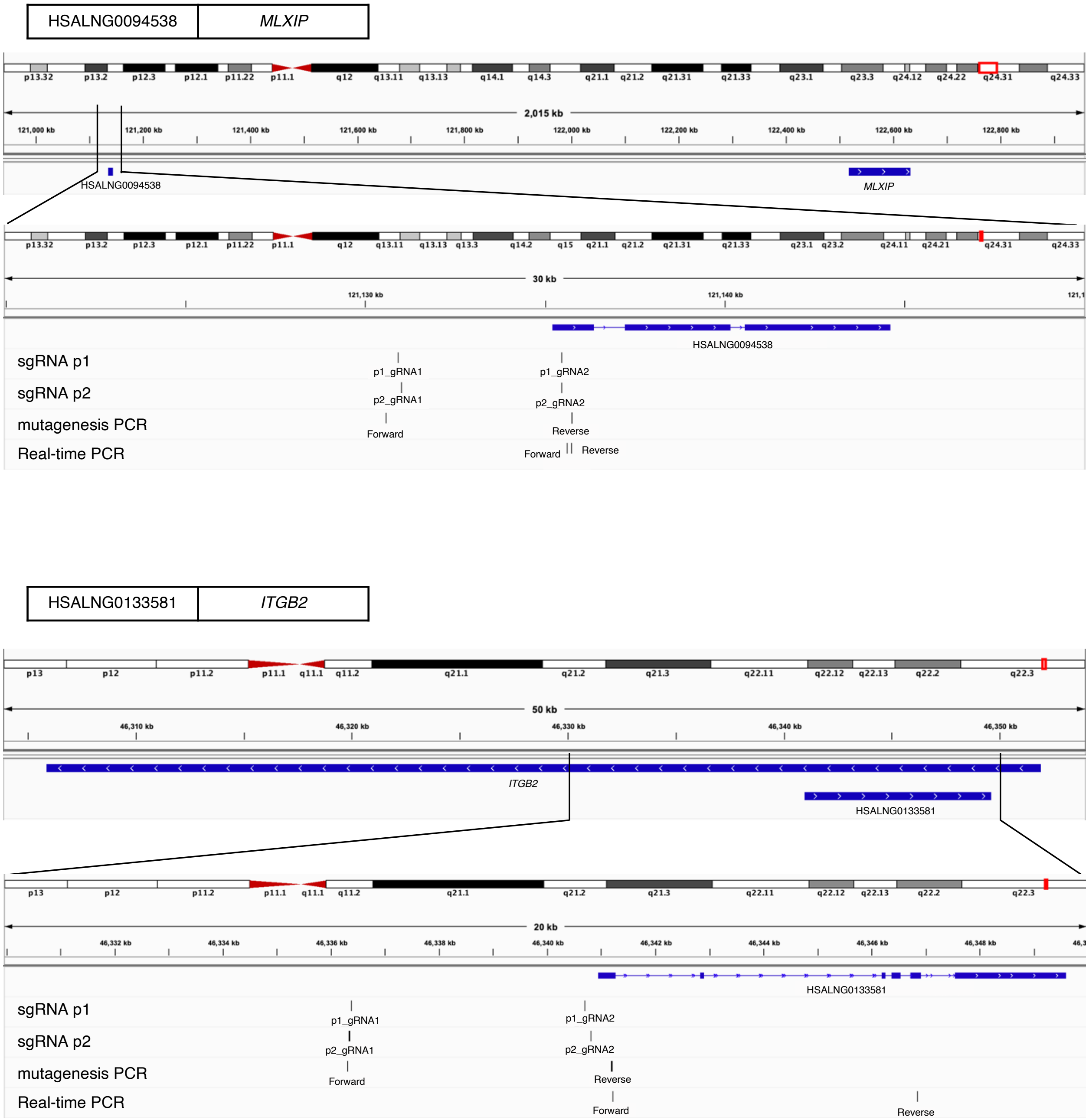


**Supplemental Fig. S12. The gene, sgRNA, and primer locations of putative oncogenic lncRNAs and the gene locations of the corresponding driver PCGs.**

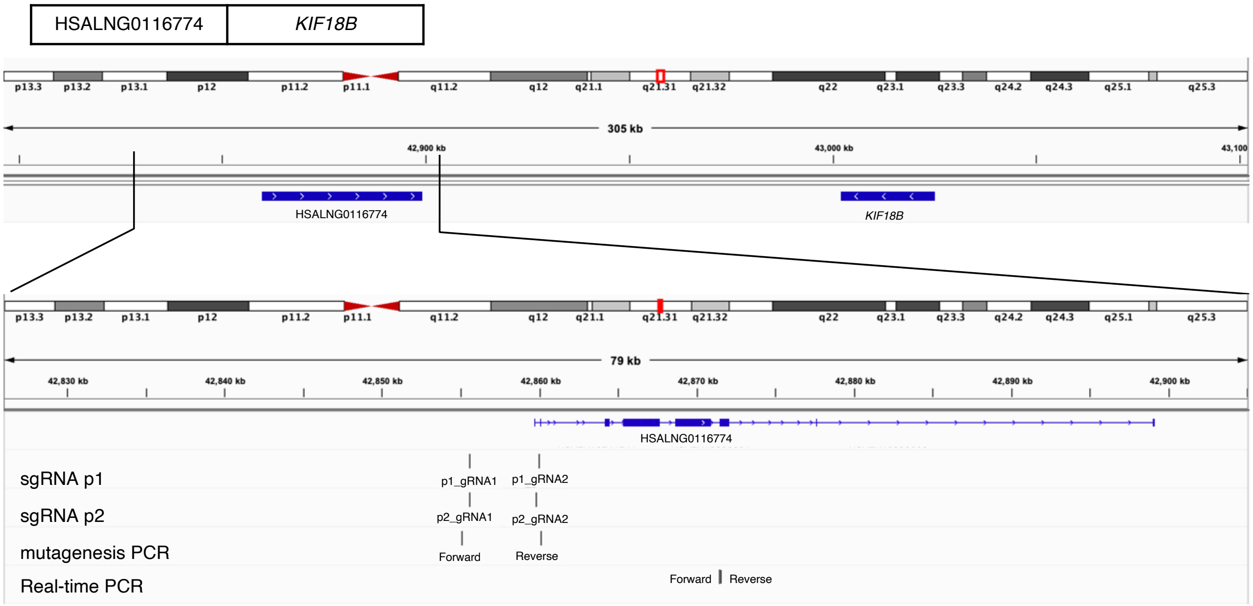
HSALNG0116774
KIF18B
p13.3 p13.2 p13.1 p12 p11.2 p11.1 q11.2 q12 q21.1 q21.31 q21.32 q22 q23.1 q23.3 q24.2 q24.3 q25.1 q25.3
305 kb
42,900 kb
43,000 kb
43,100
HSALNG0116774
KIF18B
79 kb
42,830 kb
42,840 kb
42,850 kb
42,860 kb
42,870 kb
42,880 kb
42,890 kb
42,900 kb
HSALNG0116774
sgRNA p1
p1_gRNA1
p1_gRNA2
sgRNA p2
p2_gRNA1
p2_gRNA2
mutagenesis PCR
Forward
Reverse
Real-time PCR
Forward
Reverse

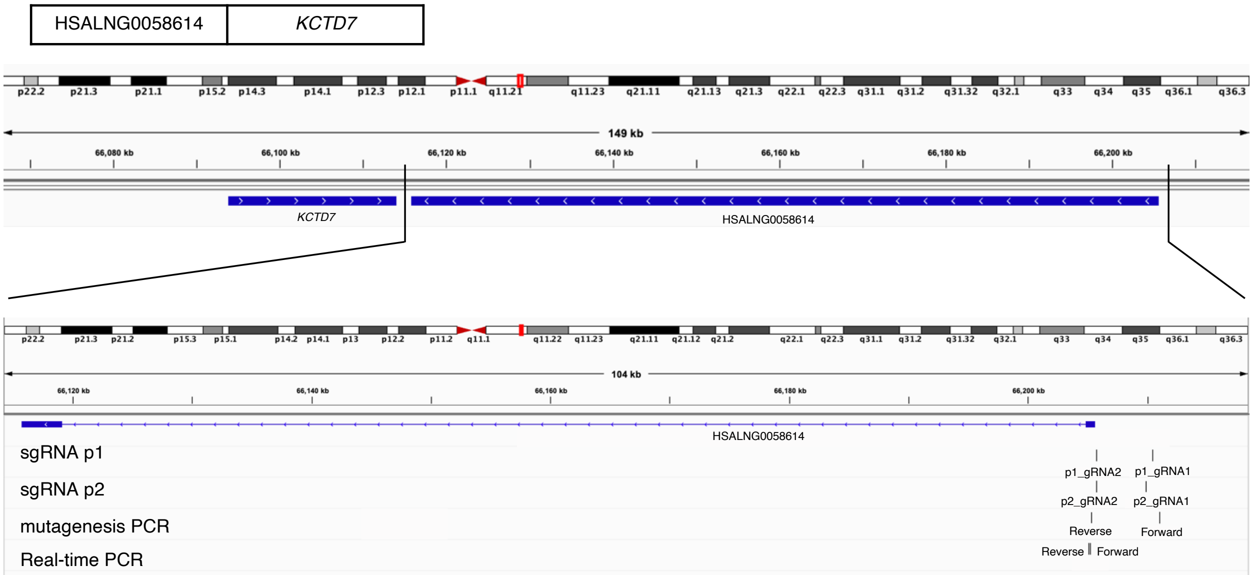
HSALNG0058614
KCTD7
p22.2 p21.3 p21.1 p15.2 p14.3 p14.1 p12.3 p12.1 p11.1 q11.21 q11.23 q21.11 q21.13 q21.3 q22.1 q22.3 q31.1 q31.2 q31.32 q32.1 q33 q34 q35 q36.1 q36.3
149 kb
66,080 kb
66,100 kb
66,120 kb
66,140 kb
66,160 kb
66,180 kb
66,200 kb
KCTD7
HSALNG0058614
104 kb
66,120 kb
66,140 kb
66,160 kb
66,180 kb
66,200 kb
HSALNG0058614
sgRNA p1
p1_gRNA2
p1_gRNA1
sgRNA p2
p2_gRNA2
p2_gRNA1
mutagenesis PCR
Reverse
Forward
Real-time PCR
Reverse
Forward

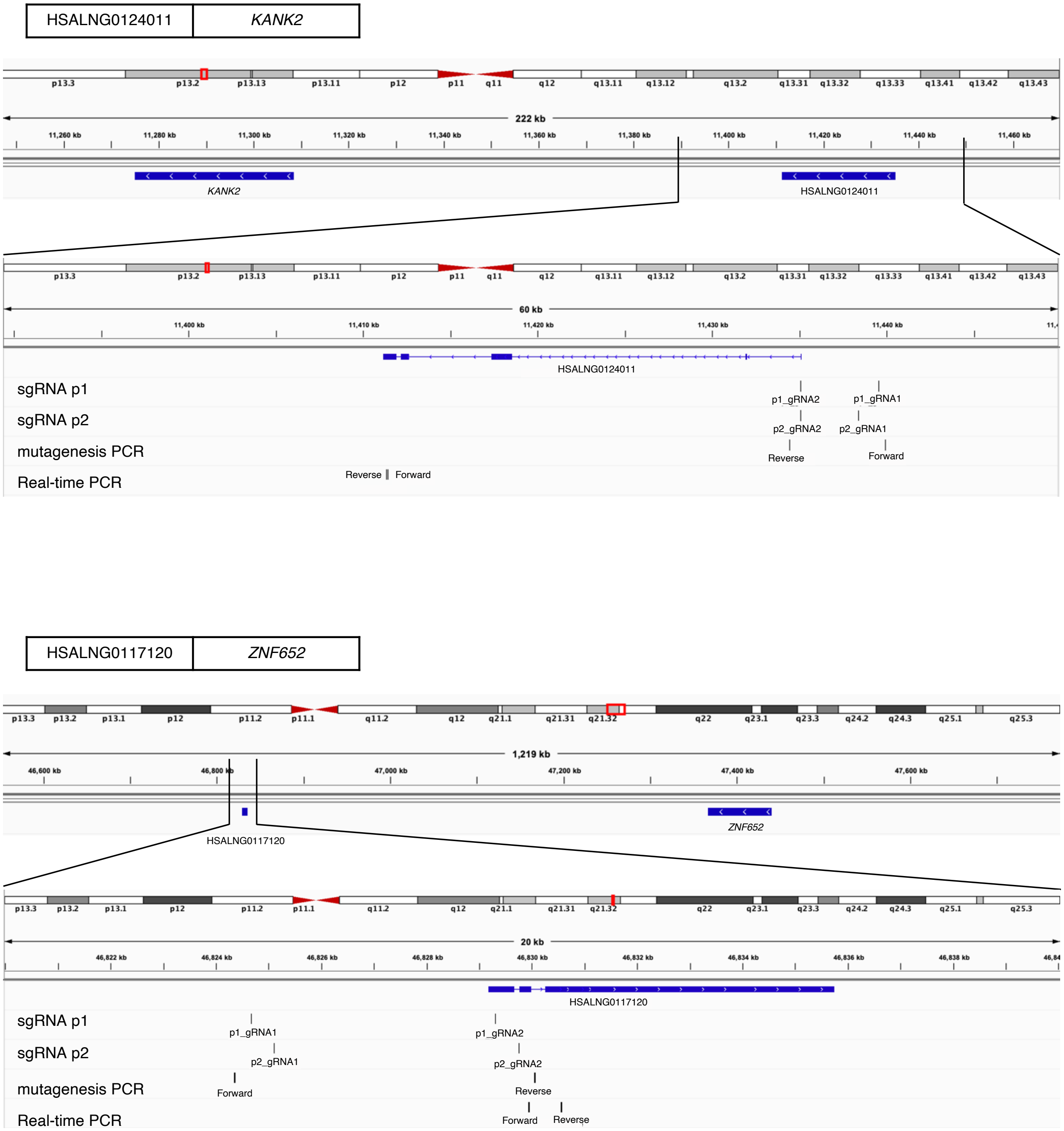


**Supplemental Fig. S13. The gene, sgRNA, and primer locations of putative tumor suppressive lncRNAs and the gene locations of the corresponding driver PCGs.**

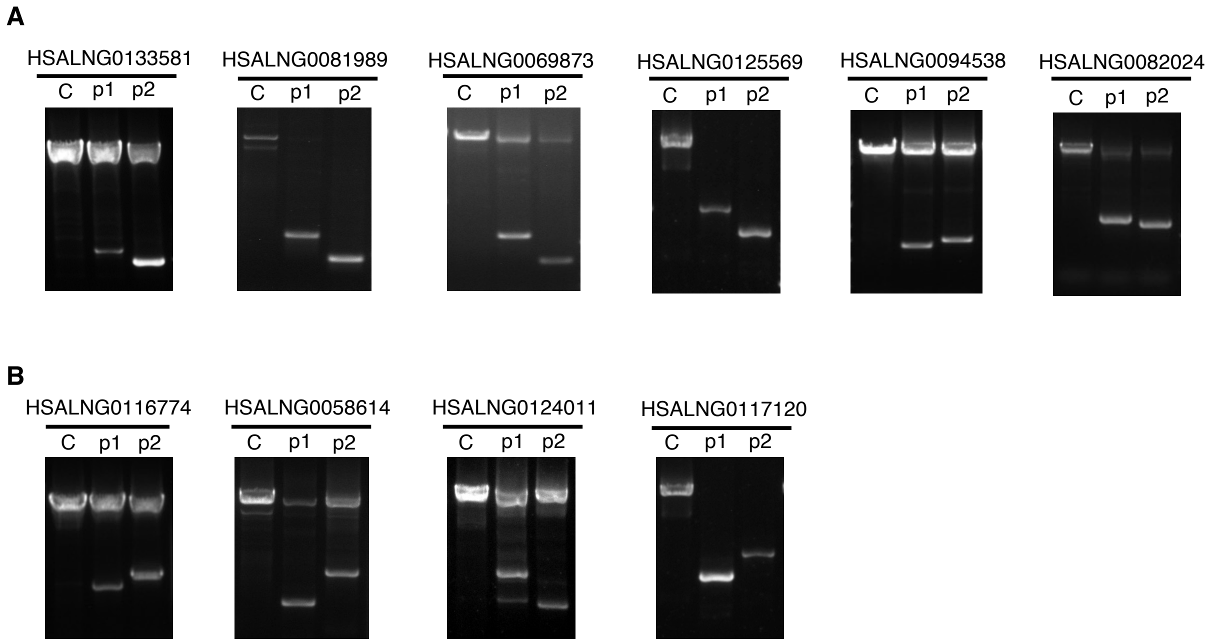


**Supplemental Fig. S14. Validation of the cutting efficiencies of guide RNAs for experiment-validated cancer driver lncRNAs in LNCaP.**

Knockout efficiency at DNA level was determined by PCR in prostate cancer cell LNCaP with each oncogenic lncRNA candidate (A) or tumor suppressive lncRNA candidate (B) disrupted by CRISPR-Cas9 guide RNA pairs p1, p2, or under control condition.

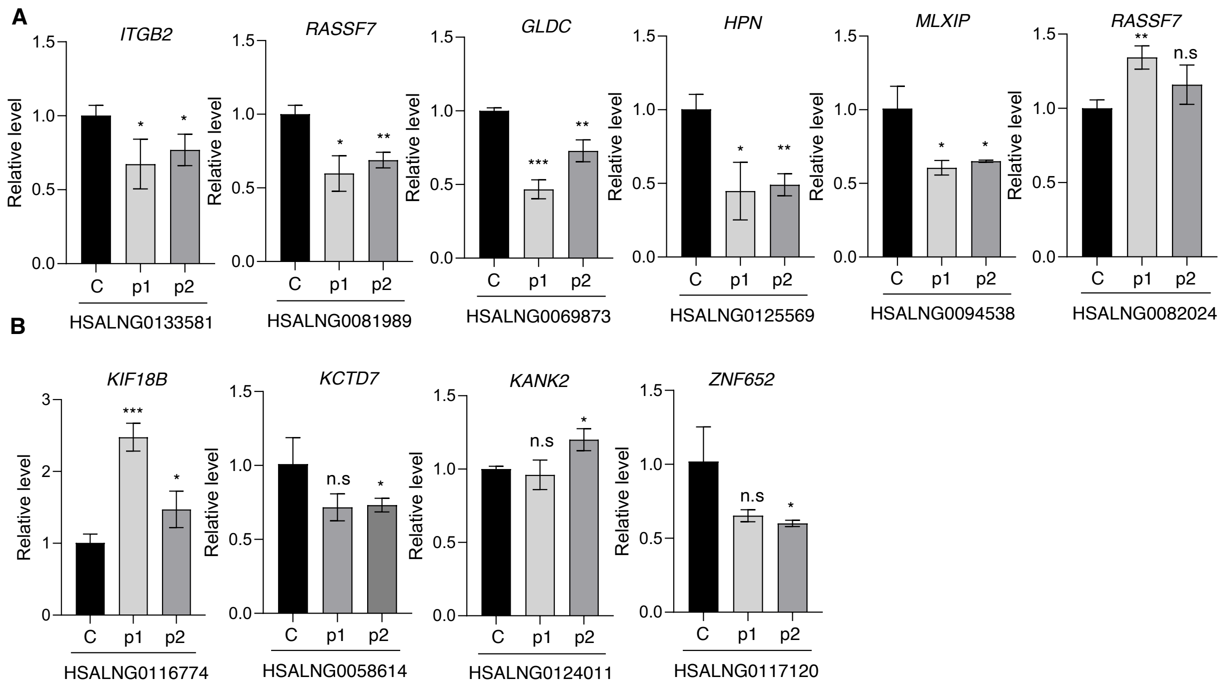


**Supplemental Fig. S15. Expression levels of the corresponding cancer driver PCGs in CRISPR-edited cells.**

The expression of the associated cancer driver PCGs expression level in prostate cancer cell line LNCaP with each putative oncogenic lncRNA (A) or tumor suppressive lncRNA (B) disrupted by CRISPR-Cas9 guide RNA g1, g2, or under control condition.

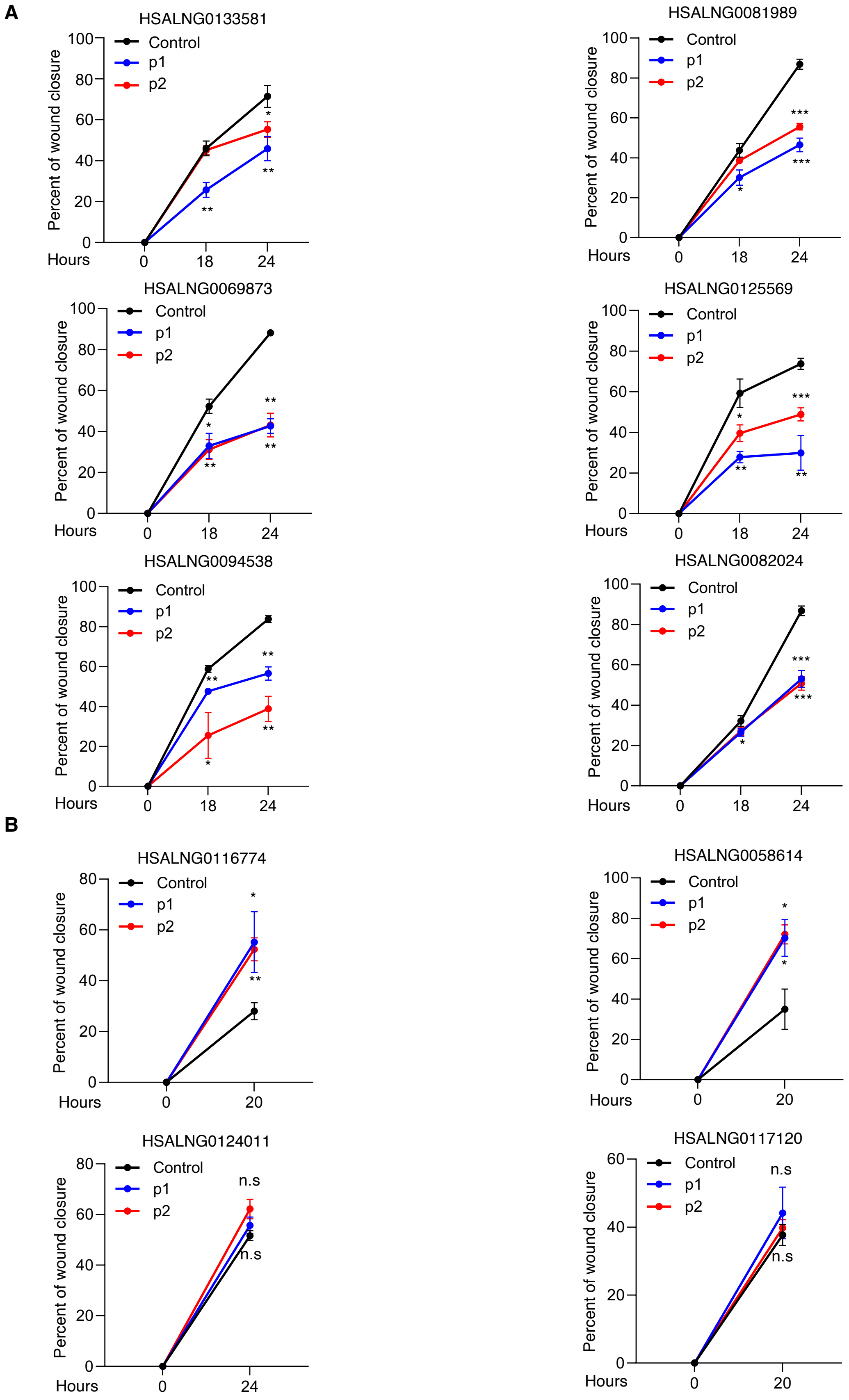
A
HSALNG0133581
HSALNG0081989
HSALNG0069873
HSALNG0125569
HSALNG0094538
HSALNG0082024
B
HSALNG0116774
HSALNG0058614
HSALNG0124011
HSALNG0117120
Control
p1
p2
Percent of wound closure
Hours
n.s

**Supplemental Fig. S16. Quantification of wound healing assay.**

The quantification of wound healing assay of PC-3 prostate cancer cells with each putative oncogenic lncRNA (A) or tumor suppressive lncRNA (B) disrupted by CRISPR-Cas9 guide RNA g1, g2, or under control condition.

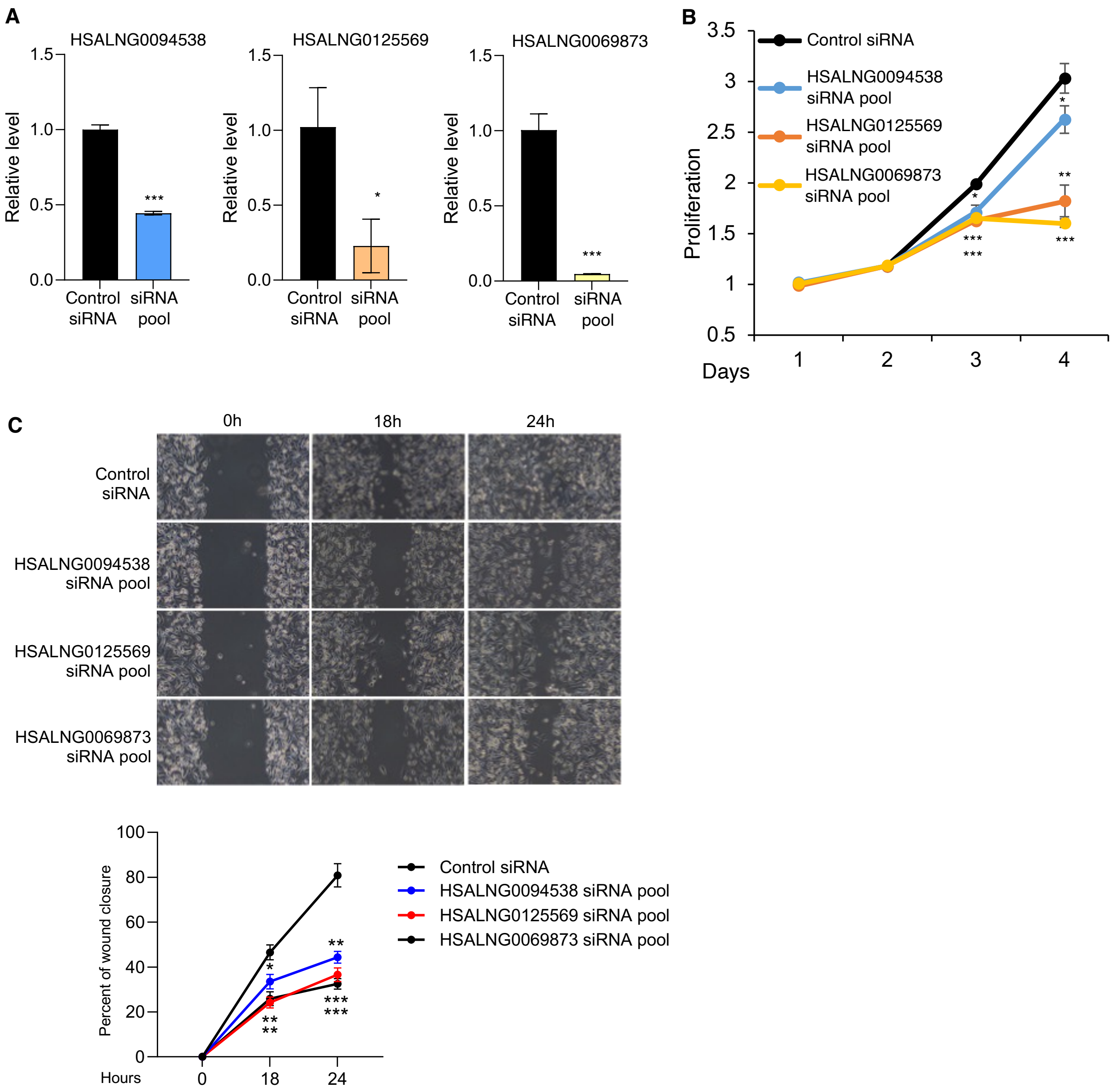


**Supplemental Fig. S17. The knockdown of candidate lncRNAs by siRNA pools decreased the cancer cell proliferation and motility.**

LncRNA expression level (A), cell proliferation (B), and wound healing assay (C) with each lncRNA candidate disrupted by siRNA pools.

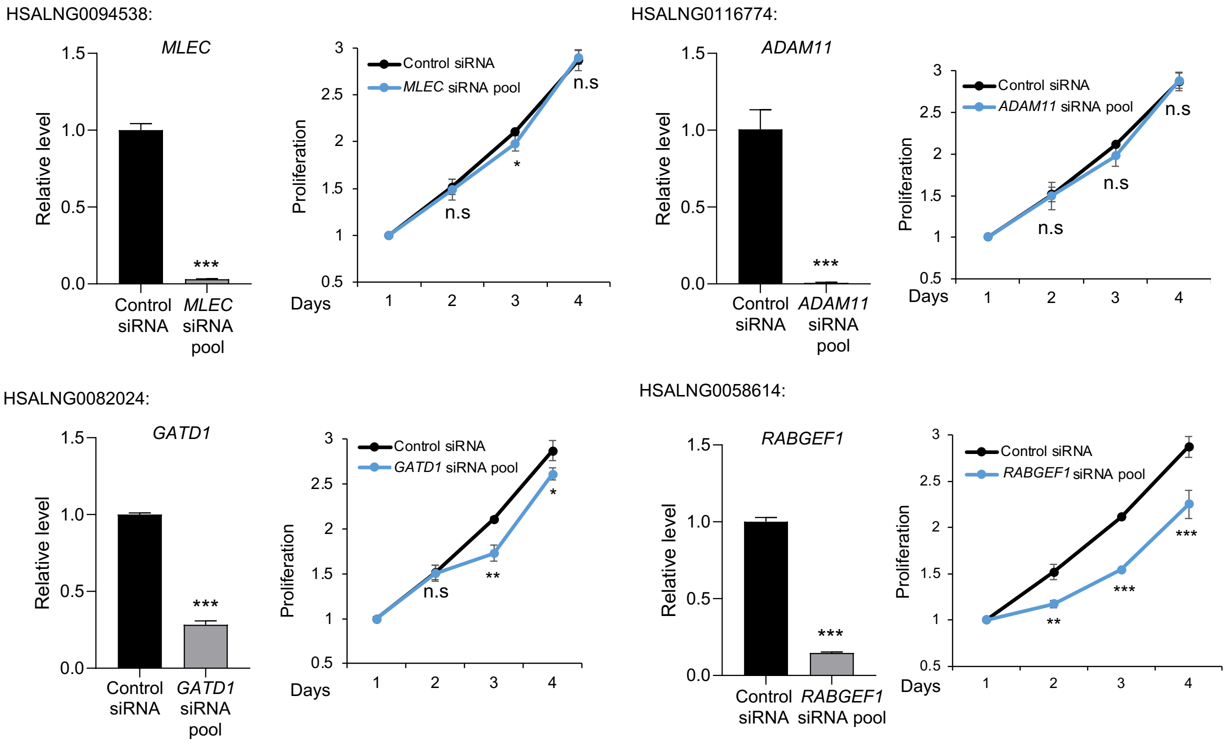


**Supplemental Fig. S18. The expression level of indicated genes and the cell proliferation in the LNCaP cells with knockdown of the overlapped protein-coding genes by siRNA pools.**

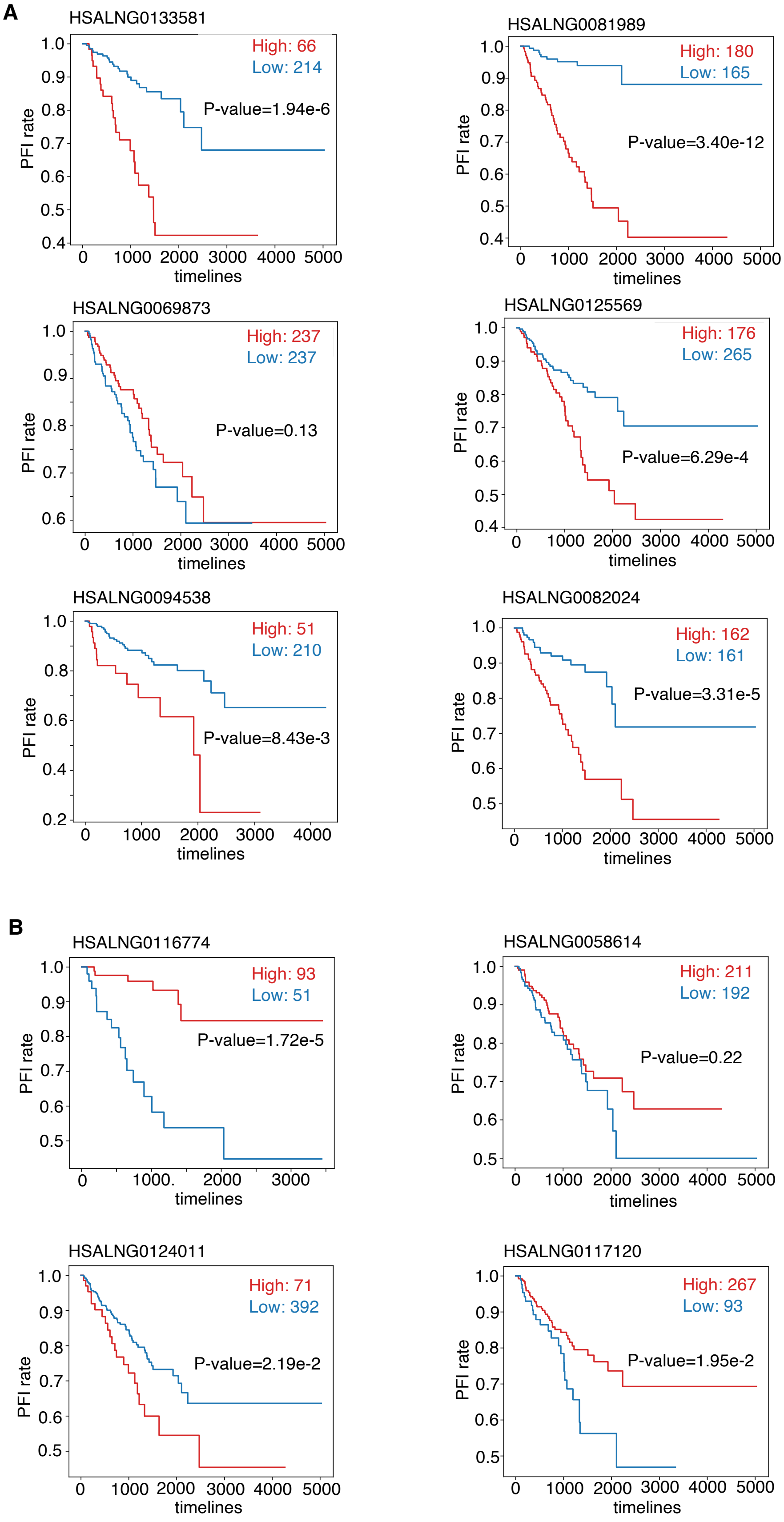
A
HSALNG0133581
High: 66
Low: 214
P-value=1.94e-6
PFI rate
timelines
HSALNG0081989
High: 180
Low: 165
P-value=3.40e-12
PFI rate
timelines
HSALNG0069873
High: 237
Low: 237
P-value=0.13
PFI rate
timelines
HSALNG0125569
High: 176
Low: 265
P-value=6.29e-4
PFI rate
timelines
HSALNG0094538
High: 51
Low: 210
P-value=8.43e-3
PFI rate
timelines
HSALNG0082024
High: 162
Low: 161
P-value=3.31e-5
PFI rate
timelines
B
HSALNG0116774
High: 93
Low: 51
P-value=1.72e-5
PFI rate
timelines
HSALNG0058614
High: 211
Low: 192
P-value=0.22
PFI rate
timelines
HSALNG0124011
High: 71
Low: 392
P-value=2.19e-2
PFI rate
timelines
HSALNG0117120
High: 267
Low: 93
P-value=1.95e-2
PFI rate
timelines

**Supplemental Fig. S19. Survival analysis of the experiment validated cancer driver lncRNAs in prostate cancer.**

K-M plot of high expressed samples and low expressed samples in PRAD for each putative oncogenic lncRNA (A) or tumor suppressive lncRNA (B). P values are determined by the Log-Rank test. PFI: Progression-Free Interval.

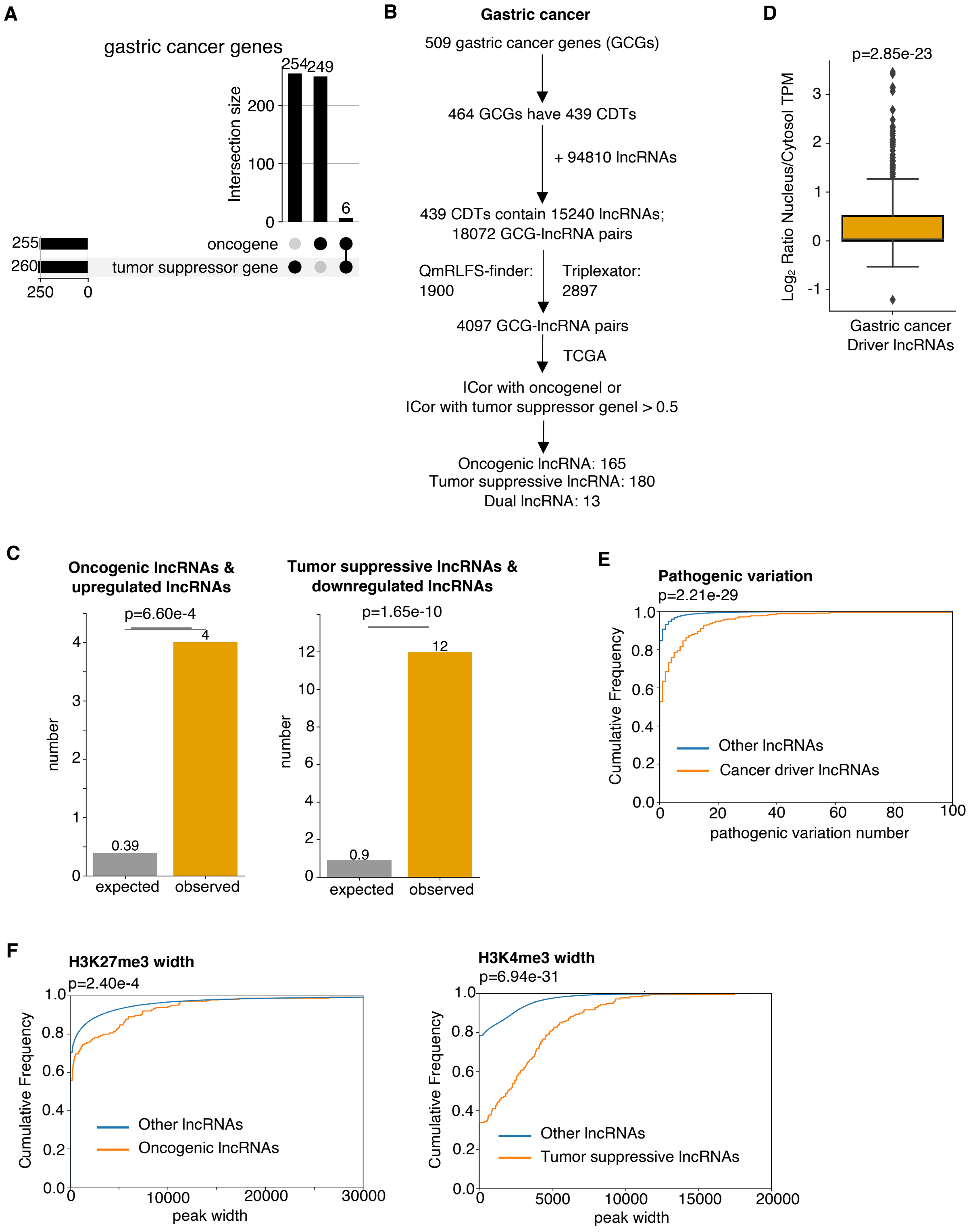


**Supplemental Fig. S20. Discovery of potential cancer driver lncRNAs in gastric cancer and validating their cancer characteristics using genomics and epigenomics data.**

A. Upset plot to show the statistics of tumor suppressor genes and oncogenes in gastric cancer.

B. The flowchart of identifying potential cancer driver lncRNAs in gastric cancer.

C. Bar plot to show the enrichment of oncogenic lncRNA candidates in upregulated lncRNAs in cancer and tumor suppressive lncRNA candidates in downregulated lncRNAs in cancer. P values are determined by one tail Fisher's exact test.

D. Boxplot showing subcellular location tendency of the putative gastric cancer driver lncRNAs. The P value is determined by one tail *t*-test.

E. Empirical cumulative distribution plot to show the difference of pathogenic variations between putative cancer driver lncRNAs and the rest other lncRNAs. P value is determined by one tail Wilcoxon test.

F. Empirical cumulative distribution plot to show the difference of H3K27me3 width between oncogenic lncRNA candidates and the rest other lncRNAs, and H3K4me3 width between tumor suppressive lncRNA candidates and the rest other lncRNAs. P values are determined by one tail Wilcoxon test.

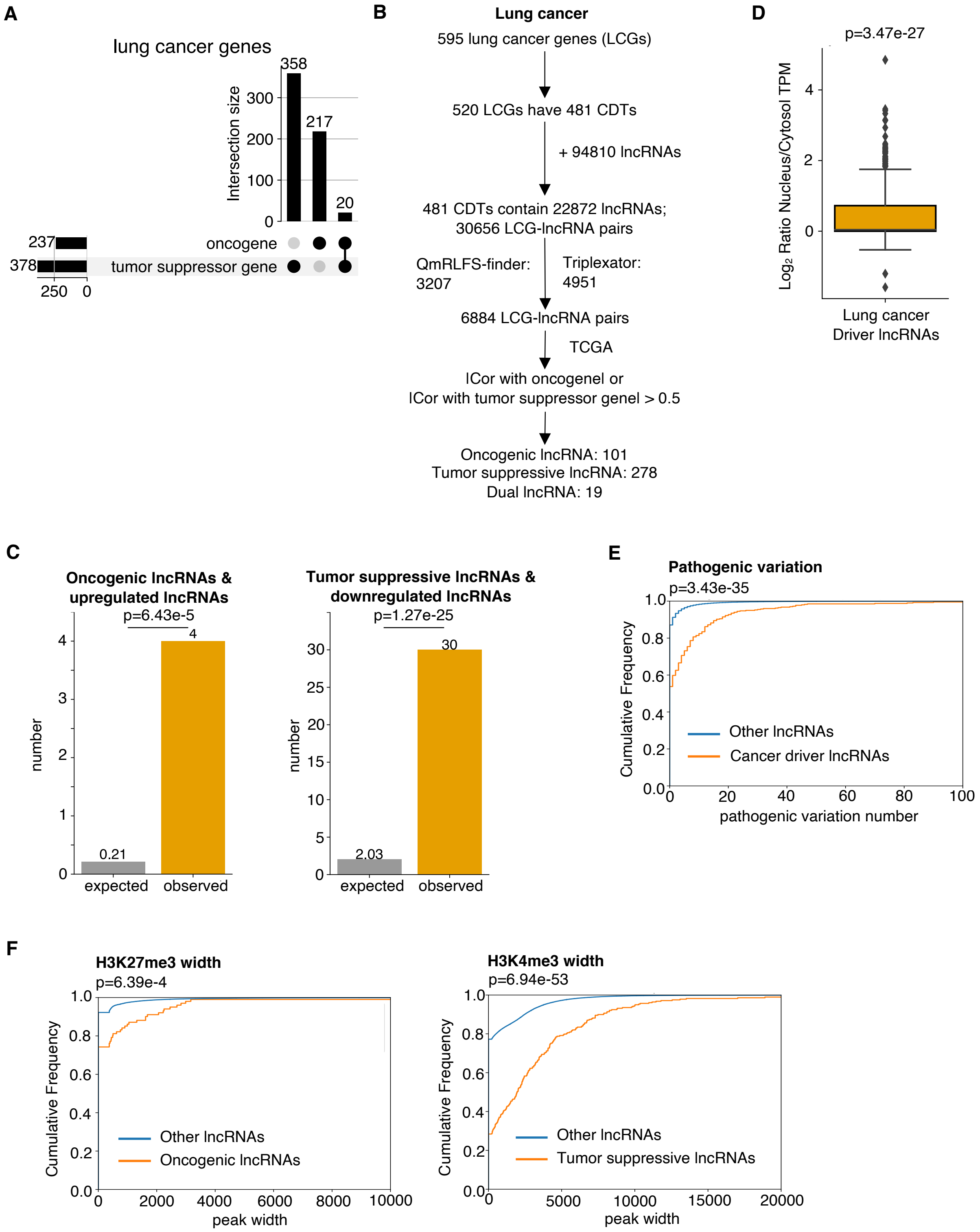


**Supplemental Fig. S21. Discovery of potential cancer driver lncRNAs in lung cancer and validating their cancer characteristics using genomics and epigenomics data.**

A. Upset plot to show the statistics of tumor suppressor genes and oncogenes in lung cancer.

B. The flowchart of identifying potential cancer driver lncRNAs in lung cancer.

C. Bar plot to show the enrichment of the oncogenic lncRNA candidates in upregulated lncRNAs in cancer and tumor suppressive lncRNA candidates in downregulated lncRNAs in cancer. P values are determined by one tail Fisher's exact test.

D. Boxplot showing subcellular location tendency of the putative lung cancer driver lncRNAs. The P value is determined by one *t*-test.

E. Empirical cumulative distribution plot to show the difference of pathogenic variations between putative cancer driver lncRNAs and the rest other lncRNAs. P value is determined by one tail Wilcoxon test.

F. Empirical cumulative distribution plot to show the difference of H3K27me3 width between oncogenic lncRNA candidates and the rest other lncRNAs, and H3K4me3 width between tumor suppressive lncRNA candidates and the rest other lncRNAs. P values are determined by one tail Wilcoxon test.
