## Supplemental Code for "Cancer Driver Topologically Associated Domains identify oncogenic and tumor suppressive lncRNAs": CADTAD_pipeline.ipynb

```
# %% [code]
import numpy as np
import pandas as pd
import os
from gtfparse import read_gtf
from Bio import SeqIO
import pybedtools
import warnings
warnings.filterwarnings('ignore')

# %% [code]
# download hg19.fa file: https://hgdownload.soe.ucsc.edu/goldenPath/hg19/bigZips/hg19.fa.gz
# download expression data file: ftp://download.big.ac.cn/lncexpdb/7-CancerCellLines/1-CancerCellLineEncyclopedia/expression_profiles_CCLE.tar.gz
# put them in ./data/raw_data/

# %% [code]
def process_gtf_to_bed(anno):
    """ 
    process gtf (1-base) to bed (0-base), 
    and key depend on gene type: lncRNA('gene_id'); protein_coding gene ('gene_name')
    """
    anno = anno[['seqname','start','end','gene_name','score','strand','gene_id']].copy()
    anno.start = anno.start - 1
    anno.score = '.'
    
    return anno

# %% [code]
def read_gene(file):
    """ 
    read gene list
    """ 
    with open(file) as f:
        gene = f.readlines()
    gene = [x.rstrip() for x in gene]
    
    return gene

# %% [code]
class CancerDriverLnc:
    """
    CADTAD: the pipeline of mining cancer driver lncRNAs
    """
    def __init__(self, gene, gene_type='gene_name'):
        self.gene = gene
        self.gene_type = gene_type
        
    def extract_gene_bed(self, gene_gtf):
        "1.extract gene bed"
        gtf = read_gtf(gene_gtf)
        if self.gene_type == 'gene_name':
            self.gene_gtf = gtf[gtf.gene_name.isin(self.gene)]
        elif self.gene_type == 'gene_id':
            self.gene_gtf = gtf[gtf.gene_id.isin(self.gene)]
        
        self.gene_bed = process_gtf_to_bed(self.gene_gtf)
        print("The number of genes:", self.gene_bed.shape[0])
    
    def gene_TAD(self, TAD_file_path):
        "2.gene_TAD"
        TAD_files = os.listdir(TAD_file_path)
        TAD_bed = [pybedtools.BedTool(''.join([TAD_file_path,i])) for i in TAD_files]
        gene_TAD = {}
        df = {'gene_name':[],'gene_id':[],'CDT':[]}
        
        for i in range(self.gene_bed.shape[0]):
            tmp_single_gene_bed = pybedtools.BedTool.from_dataframe(self.gene_bed.iloc[[i],])
            tmp_intersect = [j.intersect(tmp_single_gene_bed, wa=True, F=1) for j in TAD_bed]
            gene_TAD[list(self.gene_bed.gene_id)[i]] = tmp_intersect[0].cat(*tmp_intersect[1:])
            
            df['gene_name'].append(list(self.gene_bed.gene_name)[i])
            df['gene_id'].append(list(self.gene_bed.gene_id)[i])
            interval = tmp_intersect[0].cat(*tmp_intersect[1:]).to_dataframe()
            try:
                df['CDT'].append(interval.iloc[0,0]+':'+str(interval.iloc[0,1])+'-'+str(interval.iloc[0,2]))
            except:
                df['CDT'].append(None)
        
        self.gene_TAD = gene_TAD
        self.df = pd.DataFrame(df)
        
        print("The number of genes having TAD:", sum(-self.df.CDT.isnull()))
        print("The number of Cancer Driver TADs:", len(set(self.df.CDT)) - list(set(self.df.CDT)).count(None))
        
    
    def gene_TAD_lncRNA(self,lnc_gtf):
        "3.gene_TAD_lncRNA"
        lnc_gtf = read_gtf(lnc_gtf)
        print("total lncRNA:",lnc_gtf.shape[0])
        lnc_bed = process_gtf_to_bed(lnc_gtf)
        lnc_bed_py = pybedtools.BedTool.from_dataframe(lnc_bed)
        
        gene_TAD_lnc = []
        
        for i in self.gene_TAD.keys():
            tmp_TAD = self.gene_TAD[i]
            try:
                TAD_lnc = list(tmp_TAD.intersect(lnc_bed_py, F=1, wo=True).to_dataframe().iloc[:,9])
            except:
                TAD_lnc = None
            
            gene_TAD_lnc.append(TAD_lnc)
        
        self.df['lnc'] = gene_TAD_lnc
        self.df = self.df.dropna(axis=0)
        
        
        lnc_n = []
        for i in self.df.lnc:
            lnc_n = lnc_n + i
        
        self.lnc_bed = lnc_bed[lnc_bed.gene_id.isin(list(set(lnc_n)))]
        print("Cancer Driver TADs contain", len(set(lnc_n)), "lncRNAs")
        print("Total", sum([len(i) for i in self.df.lnc]), "gene-lncRNA pairs")
        
        # 新加的一行，为了跳过bind去算cor
        self.df_TAD = pd.DataFrame(columns=['gene_id','lnc_id','gene_name','CDT'])
        for i in self.df.index:
            gene_id = self.df.loc[i,'gene_id']
            gene_name = self.df.loc[i,'gene_name']
            CDT= self.df.loc[i,'CDT']
            lnc = self.df.loc[i,'lnc']
            
            length = len(lnc)
            
            temp_df_TAD = pd.DataFrame({'gene_id':[gene_id]*length, 'lnc_id':lnc, 'gene_name':[gene_name]*length, 'CDT':[CDT]*length})
            self.df_TAD = pd.concat([self.df_TAD, temp_df_TAD], ignore_index=True)
            
            
            
    
    # 4.bind
    def get_fasta(self, fasta_file, outpath):
        fasta = pybedtools.BedTool(fasta_file)
        gene_bed = self.gene_bed[self.gene_bed.gene_id.isin(list(self.df.gene_id))]
        gene_bed.gene_name = gene_bed.gene_id
        lnc_bed = self.lnc_bed
        lnc_bed.gene_name = lnc_bed.gene_id
        
        if os.path.exists(outpath + 'gene_fa') or os.path.exists(outpath + 'lnc_fa'):
            print('gene_fa or lnc_fa exists!')
        else:
            os.mkdir(''.join([outpath,'gene_fa']))
            os.mkdir(''.join([outpath,'lnc_fa']))
        
            for i in range(self.df.shape[0]):
                tmp_gene_bed = pybedtools.BedTool.from_dataframe(gene_bed[gene_bed.gene_id == list(self.df.gene_id)[i]])
                tmp_gene_fa = tmp_gene_bed.sequence(fi=fasta, nameOnly=True, s=True)
                tmp_lnc_bed = pybedtools.BedTool.from_dataframe(lnc_bed[lnc_bed.gene_id.isin(list(self.df.lnc)[i])])
                tmp_lnc_fa = tmp_lnc_bed.sequence(fi=fasta, nameOnly=True, s=True)
                
                with open(''.join([outpath,'gene_fa/',list(self.df.gene_id)[i],'.fa']),'w') as f:
                    f.write(open(tmp_gene_fa.seqfn).read().upper())
            
                with open(''.join([outpath,'lnc_fa/', list(self.df.gene_id)[i],'.fa']),'w') as f:
                    f.write(open(tmp_lnc_fa.seqfn).read().upper())
        
        
    def triplexator(self, script_path, gene_path, data_path):
        os.system("cat %s | /home/rzy/software/rush -j 5 'bash %s {1} %s'" % (gene_path, script_path, data_path))
    
    def QmRLFSfinder(self, script_path, gene_path, data_path, out_dir):
        os.system("cat %s | /home/rzy/software/rush -j 20 'bash %s {1} %s %s'" % (gene_path, script_path, data_path, out_dir))
    
    def QmRLFSfinder_results_process(self, gene_name, out_dir):
        try:
            lnc_path = [out_dir, 'QmRLFS-finder/', gene_name, ".out.table.txt"]
            lnc = pd.read_table(("").join(lnc_path))
        except:
            print(gene_name,"no R-loop data")
        else:
            lnc_seq = lnc[lnc.strand == "+"].sequence_RIZ
            lnc_name = lnc[lnc.strand == "+"]['#Name']
        
            gene_path = [out_dir,"gene_fa/",gene_name,".fa"]
            gene = SeqIO.read(("").join(gene_path),"fasta")
        
            num1 = [] # +
            num2 = [] # -
            for lnc in set(lnc_name):
                bind_sites1 = 0
                bind_sites2 = 0
                for seq in lnc_seq[lnc_name == lnc]:
                    bind_sites1 = bind_sites1 + gene.seq.count(seq)
                    bind_sites2 = bind_sites2 + gene.seq.reverse_complement().count(seq)
                num1.append(bind_sites1)
                num2.append(bind_sites2)

    
            # 将以上得到的结果整理成一个gene、+_bind_sites、-_bind_sites、all_bind_sites开头的表格
            data = {"gene_id":[gene_name]*len(set(lnc_name)),
                    "lnc_id":list(set(lnc_name)),
               "+_bind_sites":num1,
               "-_bind_sites":num2,
               "all_bind_sites":pd.Series(num1) + pd.Series(num2)}
            results = pd.DataFrame(data, columns = ["gene_id","lnc_id","+_bind_sites","-_bind_sites","all_bind_sites"])
    
            return(results)
    
    def gene_lncRNA_bind(self, fasta_file, outpath='./data/pipeline/bind/'):
        # fasta process
        self.get_fasta(fasta_file, outpath)
        
        # get gene_id
        self.df.gene_id.to_csv(outpath+'gene_id.txt', index=False, header=None)
        
        # triplexator
        raw_path = os.getcwd()
        script_path_1 = ''.join([raw_path,'/triplexator.sh'])
        
        os.chdir(outpath)
        data_path = os.getcwd()
        
        if os.path.exists(data_path+'/Triplexator'):
            print('Triplexator exists!')
        else:
            os.mkdir('./Triplexator')
            os.chdir('./Triplexator')
            #triplex_res_path = os.getcwd()
            
            self.triplexator(script_path_1, data_path+'/gene_id.txt',data_path)
        os.chdir(raw_path)

        # QmRLFS-finder
        script_path_2 = ''.join([raw_path,'/QmRLFS-finder.sh'])
        
        if os.path.exists(data_path+'/QmRLFS-finder'):
            print('QmRLFS-finder exists!')
        else:
            os.chdir(outpath)
            os.mkdir('./QmRLFS-finder')
            os.chdir('./QmRLFS-finder')
        
            out_dir = os.getcwd()
        
            self.QmRLFSfinder(script_path_2, data_path+'/gene_id.txt', data_path, out_dir)
        os.chdir(raw_path)
        
        
    def gene_lncRNA_bind_porcess(self, outpath='./data/pipeline/bind/'):
        # triplexator results process
        if os.path.exists(outpath+'triplextor_results.txt'):
            print('triplextor_results.txt exists!')
        else:
            with open(outpath+'triplextor_results.txt','w') as f:
                file_list = os.listdir(outpath+'Triplexator')
                matchPattern = re.compile(r'#')
                for i in file_list:
                    file = open(outpath+'Triplexator/'+i+'/'+i+'.tpx.summary')
                    lineList = []
                    while 1:
                        line = file.readline()
                        if not line:
                            break
                        elif matchPattern.search(line):
                            pass
                        else:
                            lineList.append(line)
                    file.close()
                
                    for j in lineList:
                        f.writelines(j)
        
        os.system('wc -l %s' % (outpath+'triplextor_results.txt'))
        self.triplextor_results = pd.read_table(outpath+'triplextor_results.txt', 
                                               names=['gene_id','lnc_id','Total (abs)', 'Total (rel)', 'GA (abs)', 'GA (rel)', 'TC (abs)', 'TC (rel)', 'GT (abs)', 'GT (rel)', 'other'])
        self.triplextor_results['gene_id'] = [i.split('(')[0] for i in self.triplextor_results['gene_id']]
        self.triplextor_results['lnc_id'] = [i.split('(')[0] for i in self.triplextor_results['lnc_id']]
        self.triplextor_results = pd.merge(self.triplextor_results, self.df.iloc[:,0:3], on='gene_id')
        
        # QmRLFS-finder results process
        if os.path.exists(outpath+'QmRLFS-finder_results.txt'):
            print('QmRLFS-finder_results.txt exists!')
        else:
            inCDT_lnc =  pd.DataFrame(columns = ["gene_id","lnc_id","+_bind_sites","-_bind_sites","all_bind_sites"])
            for i in self.df.gene_id:
                inCDT_lnc = inCDT_lnc.append(self.QmRLFSfinder_results_process(i, outpath), ignore_index=True)
        
            inCDT_lnc[inCDT_lnc.all_bind_sites != 0].to_csv(outpath + "QmRLFS-finder_results.txt",sep="\t",index=False)
        
        os.system('wc -l %s' % (outpath+'QmRLFS-finder_results.txt'))
        self.QmRLFSfinder_results = pd.read_table(outpath+'QmRLFS-finder_results.txt')
        self.QmRLFSfinder_results['lnc_id'] = [i.split('(')[0] for i in self.QmRLFSfinder_results['lnc_id']]
        self.QmRLFSfinder_results = pd.merge(self.QmRLFSfinder_results, self.df.iloc[:,0:3], on='gene_id')
        
        self.df_TAD_bind = pd.merge(self.triplextor_results[['gene_id','lnc_id','Total (abs)', 'Total (rel)', 'gene_name', 'CDT']], 
                                    self.QmRLFSfinder_results[['gene_id','lnc_id','all_bind_sites', 'gene_name', 'CDT']],
                                    on=['gene_id','lnc_id','gene_name', 'CDT'], how='outer').fillna(0)
    
    # 5.cor
    def coexpression(self, exp_samples, trans=True, bind=True):
        "exp_samples: expression matrix, columns are genes, rows are samples. trans: if columns are samples, rows are genes."
        exp = pd.read_table(exp_samples, index_col=0)
        if trans:
            exp = exp.T
            
        if bind:
            df_TAD_bind_cor = self.df_TAD_bind.copy()
            df_TAD_bind_cor['pearson_cor'] = 0
            df_TAD_bind_cor['spearman_cor'] = 0
        
            for i in range(df_TAD_bind_cor.shape[0]):
                df_TAD_bind_cor.iloc[i,7] = exp[df_TAD_bind_cor.iloc[i,0]].corr(exp[df_TAD_bind_cor.iloc[i,1]], method='pearson')
                df_TAD_bind_cor.iloc[i,8] = exp[df_TAD_bind_cor.iloc[i,0]].corr(exp[df_TAD_bind_cor.iloc[i,1]], method='spearman')
                
            self.df_TAD_bind_cor = df_TAD_bind_cor.copy()
        else:
            df_TAD_cor = self.df_TAD.copy()
            df_TAD_cor['pearson_cor'] = 0
            df_TAD_cor['spearman_cor'] = 0
        
            for i in range(df_TAD_cor.shape[0]):
                df_TAD_cor.iloc[i,4] = exp[df_TAD_cor.iloc[i,0]].corr(exp[df_TAD_cor.iloc[i,1]], method='pearson')
                df_TAD_cor.iloc[i,5] = exp[df_TAD_cor.iloc[i,0]].corr(exp[df_TAD_cor.iloc[i,1]], method='spearman')
                
            
            self.df_TAD_cor = df_TAD_cor.copy()
        

# %% [code]
cosmic = read_gene('./data/raw_data/cosmic.txt')

# %% [code]
pancancer = CancerDriverLnc(cosmic,'gene_name')
pancancer.extract_gene_bed('./data/raw_data/protein_coding_gene.gtf')
pancancer.gene_TAD('./data/raw_data/TAD/solid_T/')
pancancer.gene_TAD_lncRNA('./data/raw_data/lncRNA_gene.gtf')

# %% [code]
pancancer.gene_lncRNA_bind('./data/raw_data/hg19.fa', outpath='./data/pipeline/bind/')

# %% [code]
pancancer.gene_lncRNA_bind_porcess()

# %% [code]
pancancer.coexpression('./data/raw_data/CCLE_geneTPM.tsv')

# %% [code]
pancancer.df_TAD_bind_cor

# %% [code]
cosmic_gene_info = pd.read_table('../0.raw_data/cosmic_genes_info.txt',sep=' ')

# %% [code]
cancer_driver_lncRNA = pd.merge(pancancer.df_TAD_bind_cor[np.abs(pancancer.df_TAD_bind_cor['spearman_cor']) > 0.5], cosmic_gene_info, on='gene_name')

CDL_type = cancer_driver_lncRNA.groupby(['lnc_id'])[['TSG','OG']].sum()
CDL_type['type'] = 'CDL'
CDL_type['type'][CDL_type['OG'] == 0] = 'Tumor suppressive lncRNA'
CDL_type['type'][CDL_type['TSG'] == 0] = 'Oncogenic lncRNA'
CDL_type['type'][(CDL_type['OG'] != 0) & (CDL_type['TSG'] != 0)]= 'Dual lncRNA'
CDL_type = pd.DataFrame({'lnc_id':list(CDL_type.index),'lnc_type':list(CDL_type.type)})

cancer_driver_lncRNA = pd.merge(cancer_driver_lncRNA, CDL_type, on='lnc_id')
#cancer_driver_lncRNA.to_csv('./data/pipeline/cancer_driver_lncRNAs.txt', sep='\t', index=False)

# %% [code]
CDL_type.drop_duplicates()['lnc_type'].value_counts()

# %% [code]
```
