## Supplemental Code for "Cancer Driver Topologically Associated Domains identify oncogenic and tumor suppressive lncRNAs": QmRLFS-finder.sh

```
#!/bin/bash

gene=$1
data_path=$2
out_dir=$3

#source ~/software/miniconda3/bin/activate
#conda activate py2

conda run -n py2 python /home/rzy/lncRNA/data_and_src/Figure_3/software/QmRLFS-finder.py -i $data_path/lnc_fa/${gene}.fa -o $out_dir/${gene}
```
