## Supplemental Code for "Cancer Driver Topologically Associated Domains identify oncogenic and tumor suppressive lncRNAs": triplexator.sh

```
#!/bin/bash

export PATH=$PATH:/home/rzy/lncRNA/data_and_src/Figure_3/software/triplexator-1.3.2-Linux/bin
export LD_LIBRARY_PATH=$LD_LIBRARY_PATH:/home/rzy/lncRNA/data_and_src/Figure_3/software/triplexator-1.3.2-Linux/bin


gene=$1
data_path=$2

mkdir $data_path/Triplexator/${gene}
cd $data_path/Triplexator/${gene}
triplexator -ss $data_path/lnc_fa/${gene}.fa -ds $data_path/gene_fa/${gene}.fa -o ${gene}.tpx -p 8
```
